# Detection of Fluorescent Protein Mechanical Switching in Cellulo

**DOI:** 10.1101/2024.01.10.575065

**Authors:** T. Curtis Shoyer, Kasie L. Collins, Trevor R. Ham, Aaron T. Blanchard, Juilee N. Malavade, Jennifer L. West, Brenton D. Hoffman

## Abstract

The ability of cells to sense and respond to mechanical forces is critical in many physiological and pathological processes. However, the mechanisms by which forces affect protein function inside cells remain unclear. Motivated by in vitro demonstrations of fluorescent proteins (FPs) undergoing reversible mechanical switching of fluorescence, we investigated if force-sensitive changes in FP function could be visualized in cells. Guided by a computational model of FP mechanical switching, we develop a formalism for its detection in Förster resonance energy transfer (FRET)-based biosensors and demonstrate its occurrence *in cellulo* in a synthetic actin-crosslinker and the mechanical linker protein vinculin. We find that *in cellulo* mechanical switching is reversible and altered by manipulation of cellular force generation as well as force-sensitive bond dynamics of the biosensor. Together, this work describes a new framework for assessing FP mechanical stability and provides a means of probing force-sensitive protein function inside cells.

**MOTIVATION:** The ability of cells to sense mechanical forces is critical in developmental, physiological, and pathological processes. Cells sense mechanical cues via force-induced alterations in protein structure and function, but elucidation of the molecular mechanisms is hindered by the lack of approaches to directly probe the effect of forces on protein structure and function inside cells. Motivated by in vitro observations of reversible fluorescent protein mechanical switching, we developed an approach for detecting fluorescent protein mechanical switching *in cellulo*. This enables the visualization of force-sensitive protein function inside living cells.

## INTRODUCTION

The ability of cells to sense and respond to mechanical forces is critical in many developmental and physiological processes, and its dysregulation is involved in the progression of several disease states, including fibrosis and cancer^1^. To sense mechanical stimuli, cells must convert forces into biochemically detectable signals, which occurs through a multi-step molecular process^1-4^. Forces are first transmitted across specific proteins (termed mechanotransmission). This results in force-induced changes in protein structure and function (termed mechanosensing), such as the unfolding of a domain to expose a cryptic binding site^3^. Such protein conformational changes are then recognized biochemically (termed mechanotransduction), often through the binding/unbinding of transducer proteins. These new protein complexes drive downstream alterations in cell signaling and gene expression (termed mechanoresponse)^3^. Despite significant progress in our understanding of the initial and final steps of this process, elucidating the molecular mechanisms of mechanosensing and mechanotransduction remains challenging^1,5^.

In vitro single molecule techniques have provided a physical understanding of mechanosensitive molecular mechanisms. Using these techniques, the extension and unfolding of mechanosensitive protein domains, such as those in talin and alpha-catenin, as well as the subsequent binding of transducer proteins, such as vinculin, have been directly characterized^6-8^. However, determining where and when these processes occur in cells is still challenging. Progress in the spatiotemporal regulation of mechanosensitive processes inside cells has largely come from the emergence of imaging techniques^9^. The development of molecular tension sensors (MTSs) to visualize loads across specific proteins inside cells has advanced our understanding of mechanotransmission, specifically which proteins transmit loads and how these loads vary across biological contexts^10^. Progress in mechanosensing has been enabled by techniques to label unfolded protein domains via the binding of secondary probes, including antibodies that recognize the extended conformations of p130Cas or alpha-catenin^11,12^, and STReTCh, which operates by the force-induced exposure of SpyTag and subsequent covalent binding of SpyCatcher^13^. Likewise, advances in mechanotransduction have been made by monitoring the localization of endogenous transducer proteins in response to molecular tension across a load-bearing protein^14^. However, labeling could affect protein function or compete with the binding of endogenous transducers for detection, and these tools are limited to fixation and/or depend on target protein-specific reagents that can be difficult to develop. Also, these *in cellulo* techniques for mechanosensing and mechanotransduction are based on the binding of a secondary probe (a synthetic marker or a labeled natural protein), which conflates the two steps. Currently, there are no approaches analogous to in vitro techniques that can measure mechanosensing (i.e. force-induced deformation of protein domains) inside cells.

To address this technological gap, we asked if the function of MTSs could be extended. MTSs were designed to measure the magnitude of loads on proteins, i.e. mechanotransmission. The largest class of MTSs used in cells are genetically encoded Förster resonance energy transfer (FRET)-based MTSs^10,15^.

They consist of two fluorescent proteins (FPs) separated by an extensible domain^16^. Load across the MTS deforms the extensible domain, altering the distance between the FPs and the FRET efficiency. The FPs inside MTSs are in the line of loading, but their photophysical properties have been assumed to be force-insensitive. However, recent in vitro experiments have demonstrated that GFP fluorescence can be switched on/off by cycles of mechanical loading^17^. This process is reversible and associated with an intermediate transition that is distinct from the complete unfolding or denaturation of the FP^17,18^. Therefore, FP mechanical switching is a reversible, force-sensitive transition between two structural/functional states. Additionally, FP mechanical switching is a kinetic process that inherently depends on both the magnitude and dynamics of loading (e.g. load duration or rate). This is important because both the magnitude and dynamics of mechanical forces are known to drive mechanosensing by endogenous proteins^5,8,19-21^, as well as cell-level responses to mechanical stimuli^3,4^. We thus hypothesized that FP mechanical switching within FRET-based MTSs could be used to measure how force affects protein structure/function inside cells, independent of secondary probe binding. At the same time, we also reasoned that the continued use and design of MTSs to measure mechanotransmission would require an understanding of how FP mechanical switching affects this.

Here, we investigated if FP mechanical switching occurs inside cells. To create a physical framework, we first developed a kinetic model of FP mechanical switching in the context of FRET-based MTSs and then simulated expected experimental readouts. This revealed the effect of FP mechanical switching on FRET-based MTSs and predicted unique data signatures for the detection of FP mechanical switching *in cellulo*. Guided by this framework, we found that a synthetic actin-binding domain tension sensor (ABDTS) exhibited strong signatures of FP mechanical switching. The effect was reversed by pharmacological disruption of F-actin, suggesting a lack of irreversible denaturation of the FPs. We also found less, but detectable, FP mechanical switching in a tension sensor for the mechanical linker protein vinculin (VinTS). FP mechanical switching in vinculin was sensitive to both manipulations of the vinculin-actin catch bond and mechanical stiffness of the external microenvironment. Together, this work describes a new paradigm for detecting the effect of mechanical loads on fluorescent protein function in cells. This enables the visualization of force-dependent protein structure/function independent of secondary probe binding, which can be leveraged to probe mechanosensitive processes *in cellulo*.

## RESULTS

### Development of a framework to assess FP mechanical switching *in cellulo*

To investigate FP mechanical switching in the context of fusion proteins in cells, we first considered a single FP within a load-bearing protein (Note S1 and Figure S1). Specifically, we modeled a load-bearing protein subject to binding/unbinding dynamics with a single FP in the line of loading. The FP can reversibly switch between functional and non-functional states in a force-sensitive manner. Informed by single molecule studies on the response of GFP to mechanical loading, FP mechanical switching is modeled as a fast, near-equilibrium transition at a characteristic force that permits a subsequent transition to a non-fluorescent state with a Bell model force-dependent rate constant^17,22^. As a large number of FPs with different structures, photophysical properties, and mechanical stabilities exist, and the integration of FPs into fusion proteins in the cellular environment can alter these properties, we assessed the extent of FP mechanical switching over a large parameter space^16,22-24^. The model identified regions of the mechanical switching parameter space in which FP mechanical switching was likely or unlikely to occur as a function of the load magnitudes and durations estimated for protein loading in cells^10,25^ (Note S1 and Figure S1). These analyses map the conditions where various FPs may function as sensors of mechanosensing (i.e. presence of mechanical switching) versus mechanotransmission (i.e. absence of mechanical switching) when incorporated into MTSs. Additionally, for a given FP, they indicate the sensitivities of FP mechanical switching to load magnitude and load duration (Note S1 and Figure S1).

To investigate the detection of FP mechanical switching in FRET-based MTS, we first extended the model of FP mechanical switching to an MTS. To do so, we considered two FPs within the line of loading. To mediate FRET these FPs must have distinct photophysical properties, where one FP (the donor) can non-radiatively transfer energy to the other FP (the acceptor)^26^. In our model, MTSs are loaded by an actin structure to which they bind/unbind, during which the acceptor and donor FP can reversibly switch between functional and non-functional states in a force-sensitive manner (Figure 1a, Note S1, and Figure S2). Therefore, within a population of MTSs, each sensor exists in one of four states (Figure 1b).

**Figure 1.**
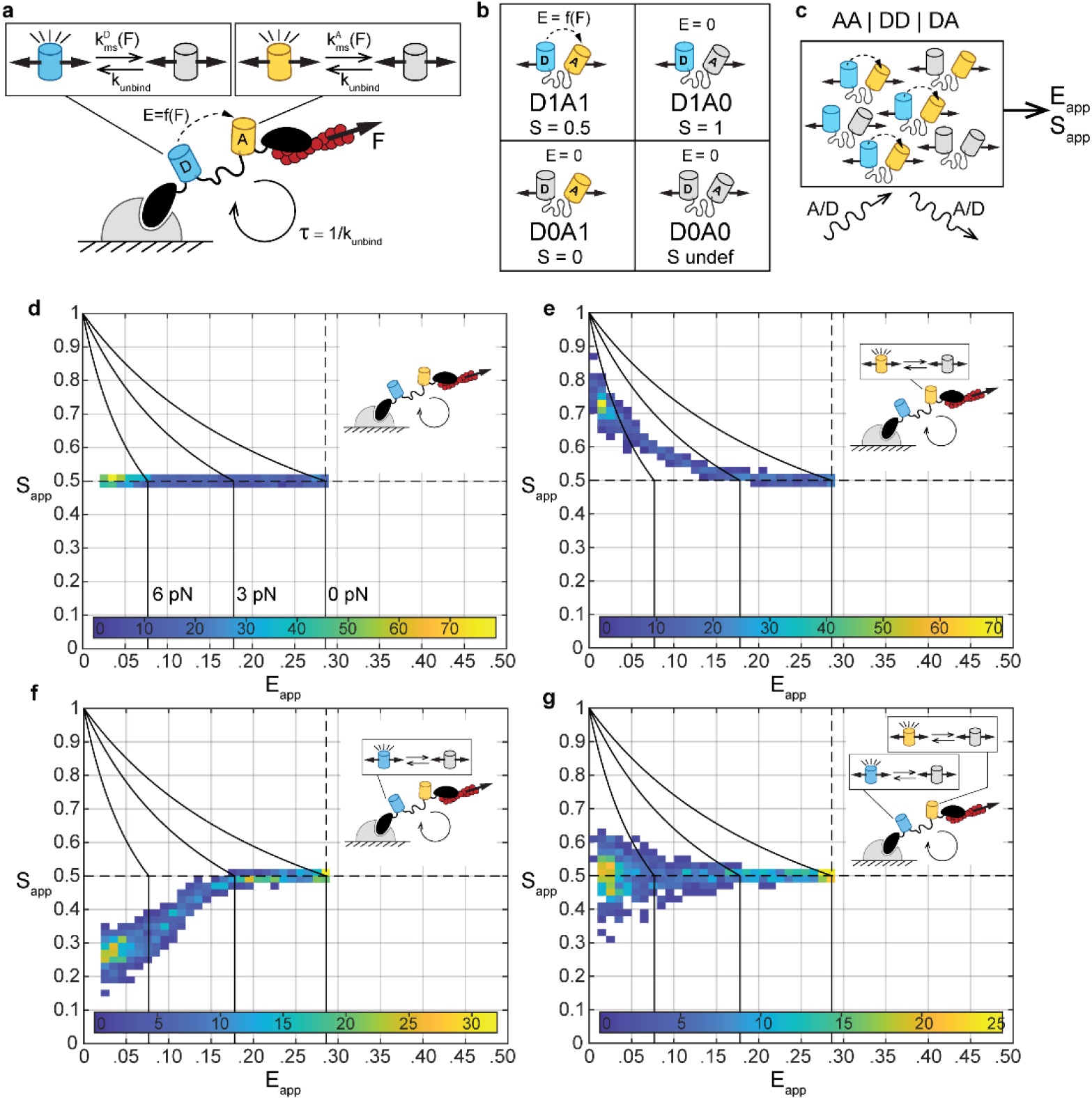
Computational model predicting data signatures for FP mechanical switching in MTSs. (a) Model of FP mechanical switching in MTSs. MTSs loaded by an actin structure to which they bind/unbind. Donor and acceptor FPs are in the line of loading and can reversibly switch between functional and non-functional states in a force-sensitive manner. (b) Four possible MTS states based on the status of the donor and acceptor FP. For each state, the FRET efficiency, E, and FP stoichiometry, S, are indicated. (c) Schematic of three-channel FRET measurements of a simulated population of MTSs. ES-histograms for populations of MTSs subject to forces ranging from 0 to 10 pN with (d) no FP mechanical switching, (e) acceptor mechanical switching only, (f) donor mechanical switching only, and (g) identical acceptor and donor mechanical switching, according to base model parameters given in Note S1, Table S2. Histograms show 1000 simulated MTS populations, each comprised of 50 sensors subject to the same load magnitude, *FF*, drawn from a uniform distribution from 0 to 10 pN. In all plots, reference black lines are tension isoclines for acceptor only or donor only mechanical switching at *FF* of 0, 3, and 6 pN (from right-to-left).

Sensitized emission is a widely used imaging modality for measuring FRET-based sensors^27-29^. By imaging in three channels, the apparent FRET efficiency, *E*_*app*_, and FP stoichiometry, *S*_*app*_ (relative abundance of donor to total FPs), can be determined using existing calibration approaches^27-29^. Using the FRET efficiency-force relationship, *E*(*F*), for the original TSMod^30^, we simulated three channel FRET measurements for populations of MTSs subject to a force, *FF*, that exhibit donor and/or acceptor mechanical switching (based on the signal contribution and number of MTS in each state) (Figure 1c). A recently developed approach for plotting calibrated three channel FRET data is a two-dimensional histogram of *E*_*app*_ and *S*_*app*_^29^. To generate data signatures of FP mechanical switching in MTSs, we extended this approach to include the force-sensitivity of FRET-based MTSs and FP mechanical switching (Note S1 and Figures S3-S4).

In the absence of FP mechanical switching, both acceptor and donor FPs are functional (D1A1 state), and the data is distributed along *S*_*app*_ = 0.5 (Figure 1d). This corresponds to the expected FP stoichiometry for a FRET construct with 1 donor and 1 acceptor, and the spread in *E*_*app*_ is due to the loading of the MTS. FP mechanical switching results in deviations in *S*_*app*_ from 0.5. Acceptor mechanical switching increases *S*_*app*_, with larger effects at lower *E*_*app*_ values, resulting in an up/left-slanting data signature (Figure 1e). This trend is conserved for a wide range of FP mechanical switching parameters and is distinct from constitutive acceptor loss-of-function, e.g. due to photobleaching (Note S1 and Figure S5). For a given tension magnitude, acceptor mechanical switching reduces *E*_*app*_ compared to its value in the absence of mechanical switching (Note S1 and Figure S4). This is consistent with the known effect of excess donors on FRET efficiency measurements^29^. This indicates that quantification of the force magnitude using *E*_*app*_ and the eff-force calibration is no longer possible. Furthermore, donor mechanical switching decreases *S*_*app*_, with larger effects at lower *E*_*app*_ values, resulting in a down/left-slanting data signature (Figure 1f). This trend is conserved for a wide range of FP parameters and is distinct from constitutive donor loss-of-function (Note S1 and Figure S6). Unlike acceptor mechanical switching, for a given tension magnitude, donor mechanical switching does not affect *E*_*app*_ (Note S1 and Figure S4), consistent with no effect of excess acceptor on FRET efficiency measurements^29^. As we had found that single FP mechanical switching was sensitive to load duration, we tested if MTS readouts also responded to changes in load duration and found that this was indeed the case for both acceptor and donor mechanical switching (Note S1 and Figures S7 and S9). Lastly, we considered when both acceptor and donor FPs undergo mechanical switching. If the acceptor and donor exhibit identical FP mechanical switching kinetics, *S*_*app*_ values deviate from 0.5 in both directions (Figure 1g), producing a signature that is distinct from absent, acceptor only, and donor only FP mechanical switching. Between the limits discussed here, both FPs may undergo mechanical switching but with different parameters. In this case, dominant mechanical switching of acceptor or donor remains detectable in the presence of lower levels of mechanical switching in the other species according to the signatures for acceptor or donor mechanical switching (Note S1 and Figure S8).

Taken together, the simulations demonstrate the effect of FP mechanical switching on three channel FRET measurements of MTSs and predict unique data signatures for the detection of FP mechanical switching *in cellulo*.

### Synthetic actin binding tension sensor exhibits FP mechanical switching *in cellulo*

Next, we investigated if FP mechanical switching could be detected *in cellulo*. To begin, we sought to assess this with a structurally simple MTS that is loaded directly by the actin cytoskeleton and is not subject to biochemical regulation. Therefore, we created the synthetic Actin Binding Domain Tension Sensor (ABDTS) by attaching the actin binding domain F-tractin to both ends of a tension sensor module (TSMod)^30^ (Figure 2h). As an unloaded control, we used a version possessing only one F-tractin domain (ABDTL) (Figure 2a). We performed three-channel FRET imaging of these constructs in NIH 3T3 cells. (Note that all experimental FRET measurements are inherently apparent FRET efficiency and FP stoichiometry, but are indicated as *E* and *S* without the subscript to match previous conventions^16,27,28,31^). Both constructs localized similarly to actin structures in NIH 3T3 cells (Figure 2b-d,i-k). ABDTL exhibited a spatially uniform FRET efficiency of ~0.285, corresponding to the unloaded value for TSMod^28^, and stoichiometry of ~0.5 (Figure 2e-f and Figure S13a-b). ES-histograms of single cells and the whole cell population contained a single major density centered on *E* ∼0.285 and *S*∼0.5 (Figure 2g,o), resembling the predicted signature for an unloaded MTS with no FP mechanical switching from the model (Note S1 and Figure S4b). In contrast, ABDTS exhibited regions of lower FRET efficiency (*E* < 0.285) and higher stoichiometry (*S* > 0.5), especially at the periphery of the cell in regions of positive curvature (Figure 2l-m and Figure S13c-d). These regions contained a gradient of increasing stoichiometry pointing outward, and at the very edge of these regions we observed that acceptor signal is nearly completely lost while donor signal is only partially lost (Figure S13e-h). We verified that these patterns were not due to an ABDTS-induced alteration or loss of actin in these regions by fixing and labeling ABDTL- and ABDTS-containing cells with phalloidin (Figure S14a-j). ES-histograms of a single cell (Figure 2n) and the whole cell population (Figure 2p) contained a major up/left-slanting density extending to lower FRET efficiencies (*E* < 0.285) and higher stoichiometries (*S* > 0.5). These data resemble the predicted signature of a loaded MTS with dominant acceptor mechanical switching from the model (Figure 1e). Additionally, at the lowest FRET efficiency values (coming mainly from pixels at the edge of the cell), the ES-histogram had a wider range of stoichiometries spanning between 0.5 and 1. This resembles the biphasic trends observed in the model for two different mechanisms: 1) more sensitive acceptor mechanical switching with less sensitive donor mechanical switching onset at higher forces (Note S1 and Figure S8c), or 2) acceptor mechanical switching with force-induced unbinding (Note S1 and Figure S10h,k). Our observation of partial loss of donor signal (in addition to near complete loss of acceptor signal) at the very edge of ABDTS-but not ABDTL-containing cells (Figure S13e-h) supports the first explanation. To quantitatively compare ABDTS versus ABDTL, we computed the fraction of pixels in each cell in a low *E*, high *S* bin (*E* < 0.15, *S* > 0.60) and found that ABDTS had a significantly higher fraction of pixels than ABDTL (Figure 2q). The trends in FRET efficiency and stoichiometry for ABDTL and ABDTS were also not altered by fixation (Figure S14k-m).

**Figure 2.**
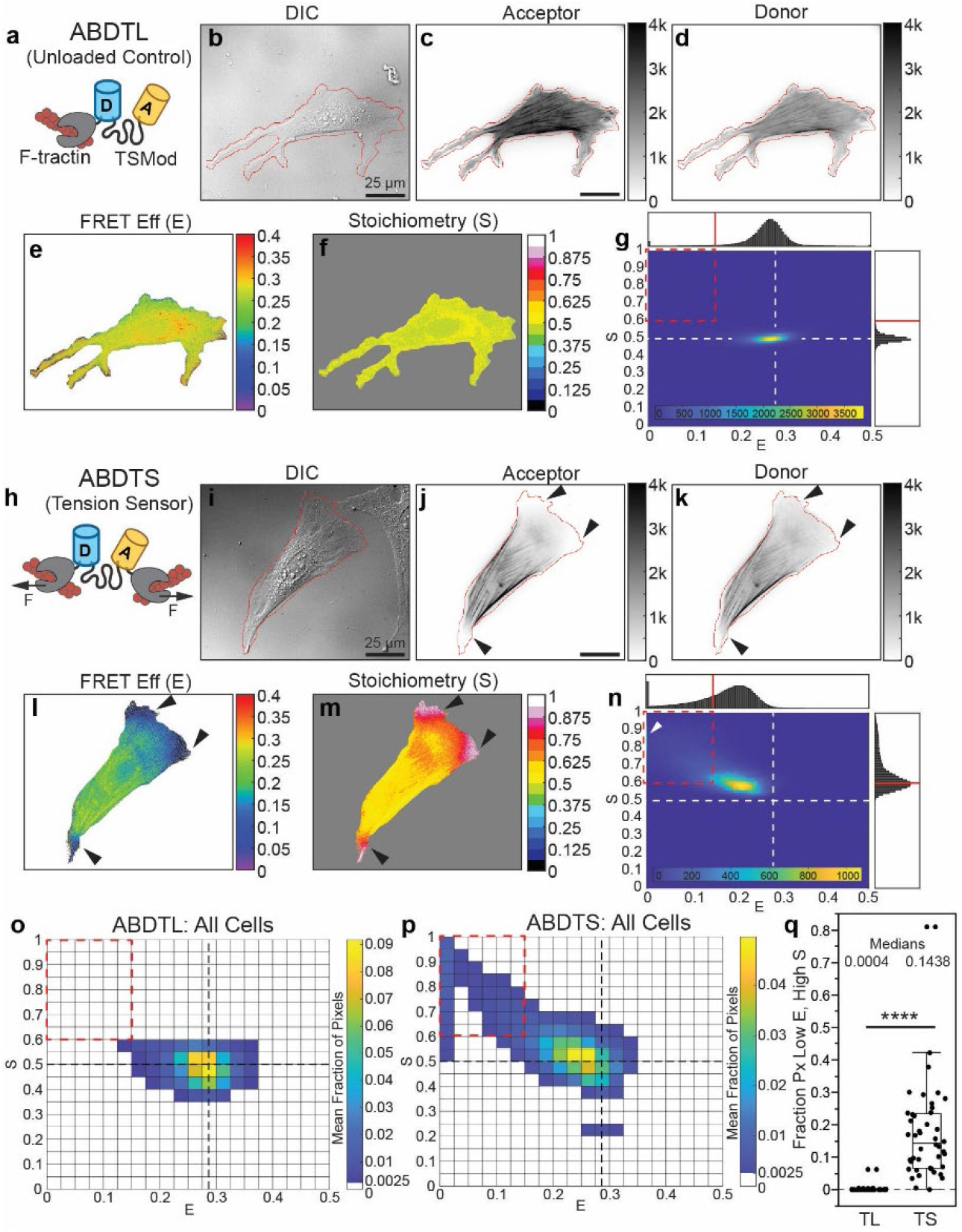
Synthetic actin binding tension sensor exhibits FP mechanical switching *in cellulo*. (a) Schematic of actin binding domain tailless control (ABDTL). (b-g) Representative NIH 3T3 cell expressing ABDTL, showing images of DIC used to create cell outline, acceptor and donor intensities with cell outline overlaid in red, FRET efficiency and Stoichiometry in cell mask, and ES-histogram of pixels in the cell. (h) Schematic of actin binding domain tension sensor (ABDTS). (i-n) Images and histogram for a representative NIH 3T3 cell expressing ABDTS, analogous to those in (b-g). ES-histograms for whole cell populations of ABDTL (o) and ABDTS (p), indicating the cell-averaged fraction of cell-masked pixels in each bin (N = 38/44 cells for ABDTL/ABDTS over 5 experimental days). (q) Box plot of fraction of pixels in each cell in the low *E*, high *S* bin (*E* < 0.15, *S* > 0.60) for ABDTL and ABDTS. Differences between groups were detected using the Steel-Dwass test, ****: p<0.0001.

The data signatures indicate that the mechanical switching of mVenus (acceptor) is dominant over mTFP1 (donor), suggesting differential mechanical stabilities for the two FPs *in cellulo*. Previously, it has been shown that even FPs derived from the same species, GFP and enhanced yellow fluorescent protein (EYFP), have different mechanical stabilities in vitro^22,23^. To test the feasibility that mVenus and mTFP1 had different mechanical stabilities, we performed steered molecular dynamics (SMD) simulaons on the two FPs, an approach previously applied to GFP^18^. In agreement with our experimental observaons, we found that mTFP1 had a higher mechanical stability than mVenus in the SMD simulaons (Note S2, Figures S15-S17, and Movies S1-S3).

To test if acceptor mechanical switching in ABDTS was manipulable, we used Latrunculin A to disrupt the actin cytoskeleton of fully spread cells (Figure S18). In Latrunculin A treated cells, ABDTS exhibited significantly increased FRET efficiency and decreased stoichiometry, indicating a reduction in acceptor mechanical switching. This demonstrates the long-timescale reversibility of this process in cells and is consistent with mechanical switching but not irreversible denaturation of the FP.

Taken together, these data demonstrate the presence of reversible FP mechanical switching in a synthetic actin binding sensor in living cells.

### Vinculin tension sensor exhibits weaker but detectable acceptor mechanical switching

We next asked if FP mechanical switching could be detected in an MTS within a naturally occurring protein. We focused on the mechanical linker protein vinculin, which couples the actin cytoskeleton at focal adhesions (FAs) to mediate adhesion reinforcement and stiffness sensing^30,32,33^. We focused on vinculin because it has been widely studied with an existing tension sensor (VinTS)^2,10,14,16,34,35^, and VinTS variants that unload vinculin^33,36^ as well as progressively disrupt vinculin catch-bonding^34^ have been recently developed. Applying our framework, we re-analyzed existing data sets of VinTS and these mutants expressed in vinculin -/- MEFs^33,34^. As the tension-insensitive control, we used VinTS-I997A, which contains a point mutation that disrupts vinculin’s binding to actin and has been shown to unload vinculin^33,36^. VinTS-I997A in FAs showed FRET efficiencies corresponding to unloaded TSMod and stoichiometries corresponding to no FP mechanical switching at both the single-cell (Figure 3a-d and Figure S19a-e) and cell population levels (Figure 3z), as expected for a tension-insensitive control. In contrast, VinTS exhibited a spectrum of behaviors. At one end of this spectrum, VinTS cells exhibited lower FRET efficiency (*E* < 0.285) with no apparent FP mechanical switching (*S*∼0.5) (Figure S19f-j), matching the model prediction for MTS loading without FP mechanical switching (Figure 1d). At the other end of this spectrum, VinTS cells exhibited both a lower FRET efficiency (*E* < 0.285) and a higher stoichiometry (*S* > 0.5) with an up/left sloping ES-histogram shape (Figure 3f-j and Figure S19k-o). This matches the prediction for MTS loading with acceptor mechanical switching (Figure 1e), albeit with distinctly less mechanical switching than observed in ABDTS (compare Figure 2p to Figure 3z). The heterogeneity in the VinTS data is likely attributable to cell-to-cell variability in contractility^37^. The presence of weaker (than ABDTS) but detectable acceptor mechanical switching in VinTS is also apparent at the cell population level (Figure 3z). To quantify this, we again looked at the fraction of pixels in each cell in a low *E*, high *S* bin (*E* < 0.15, *S* > 0.60), indicating a significant difference between VinTS and VinTS-I997A (Figure 3aa).

**Figure 3.**
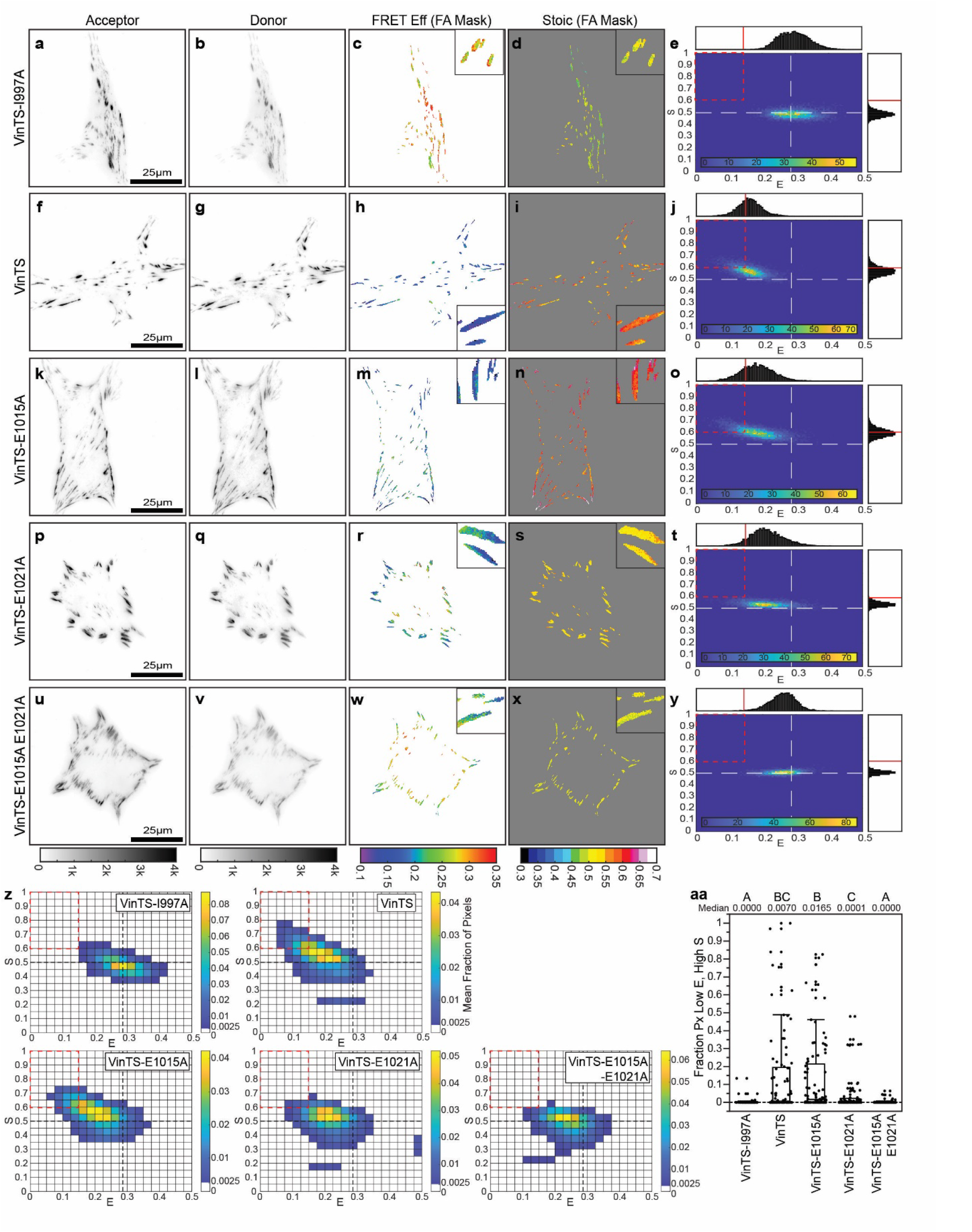
FP mechanical switching in VinTS is sensitive to manipulation of the vinculin-actin catch bond. Representative vinculin -/- MEFs expressing VinTS-I997A (a-e), VinTS (f-j), VinTS-E1015A (k-o), VinTS-E1021A (p-t), or VinTS-E1015A-E1021A (u-y) on FN-coated glass, showing images of acceptor intensity, donor intensity, FRET efficiency in the FA mask, Stoichiometry in the FA mask, and an ES-histogram of FA-masked pixels for the cell. (z) ES-histograms for whole cell populations, indicating the cell-averaged fraction of FA-masked pixels in each bin (N = 59/89/92/88/101 cells over 2/4/3/3/3 experimental days for VinTS-I997A/VinTS/VinTS-E1015A/VinTS-E1021A/VinTS-E1015A-E1021A). (aa) Box plot of fraction of pixels in each cell in the low *E*, high *S* bin (*E* < 0.15, *S* > 0.60) for each construct. Differences between groups were detected using the Steel-Dwass test. Levels not connected by the same letter are significantly different at p<0.05. P-values are given in Note S3. The data is a new analysis of the three-channel FRET images from an experiment in a previous publication^34^.

Furthermore, force-activated bond dynamics play an important role in mechanosensitive processes at FAs^2,4^, and vinculin is known to form a strong catch bond with F-actin, whose duration increases with force up to a certain point^38^. Our modeling indicated that FP mechanical switching can be sensitive to changes in load duration (Note S1 and Figures S1 and S7), as well as alterations to force-sensitive bond dynamics of the MTS (Note S1 and Figures S10-S11). To assess if manipulating vinculin catch-bonding affects FP mechanical switching, we re-analyzed the data of VinTS harboring single (VinTS-E1015A and VinTS-E1021A, Figure 3k-t) and double (VinTS-E1015A-E1021A, Figure 3u-y) point mutations that were previously shown to progressively disrupt vinculin catch-bonding while retaining other key aspects of vinculin function^34^. The ES-histograms at both single-cell and cell population levels demonstrate that the acceptor mechanical switching signature of VinTS was partially reduced in the single mutants and eliminated in the double mutant (Figure 3z). Quantifying the fraction of pixels in the low *E*, high *S* bin (*E* < 0.15, *S* > 0.60), VinTS-E1015A-E1021A was significantly lower than VinTS and similar to VinTS-I997A (Figure 3aa), indicating an elimination of acceptor mechanical switching in the double mutant. However, while VinTS-E1015A-E1021A and VinTS-I997A had no apparent FP mechanical switching (*SS*∼0.5), the ES-histogram of VinTS-E1015A-E1021A was shifted to a slightly lower FRET efficiency than VinTS-I997A (Figure 3z), consistent with the E1015A-E1021A double mutant remaining partially loaded as previously reported^34^. Together, these analyses indicate that FP mechanical switching is detectable in vinculin and requires vinculin catch-bonding.

Altering ECM stiffness is also thought to affect the loading dynamics of mechanical proteins within FAs^2,4^. In molecular clutch models of the FA, substrate stiffness is a major factor that controls protein loading rate, with lower loading rates typically occurring on softer substrates^4^. When we explicitly modeled the loading of MTS at different loading rates, our model also predicted a sensitivity of FP mechanical switching to loading rate (Note S1 and Figure S12). Therefore, we tested the hypothesis that changing the substrate stiffness would alter FP mechanical switching in vinculin. To do so, we seeded vinculin -/- MEFs stably expressing VinTS or VinTS-E1015A-E1021A on FN-coated PA gels (Figure S20).

Previous work indicated that the FRET efficiency of VinTS did not change on 10 kPa gels compared to glass but did not assess stoichiometry^16^. Therefore, we chose a softer gel (of approximately 3.5 kPa stiffness, Figure S21) and directly assessed stoichiometry. On PA gels, we observed a considerable change in the ES-histogram shape for VinTS, consistent with a loss of FP mechanical switching (Figure S20a). In contrast, the ES-histogram shape for VinTS-E1015A-E1021A on gels resembled that on glass and was consistent with little to no FP mechanical switching (Figure S20b). On these softer substrates, we found no statistical difference in the fraction of pixels in the low *E*, high *S* bin between the two constructs (Figure S20c), indicating that the softer substrate eliminated acceptor mechanical switching in VinTS.

Taken together, these results demonstrate that FP mechanical switching occurs in a naturally occurring protein, vinculin, and is sensitive to key cell-intrinsic (force-activated bond dynamics) and cell-extrinsic (substrate stiffness) parameters underlying mechanosensitive processes at FAs.

## DISCUSSION

Uncovering the molecular mechanisms of mechanosensitive signaling requires characterizing multiple steps: mechanotransmission, mechanosensing, mechanotransduction, and mechanoresponse^3^. A current challenge is understanding the spatiotemporal regulation of these steps. Progress has been limited by the lack of tools to probe the various steps inside cells, especially mechanosensing (e.g. protein domain unfolding). Motivated by the in vitro demonstration of reversible mechanical switching of GFP^17^, we use computational modeling to develop a formalism to detect FP mechanical switching in FRET-based biosensors *in cellulo* by three-channel imaging of sensitized emission. Guided by this formalism, we demonstrate that FP mechanical switching occurs *in cellulo* in both a synthetic actin crosslinking protein and the mechanical linker protein vinculin. *In cellulo* mechanical switching is reversible and sensitive to manipulations of cell-generated forces, force-sensitive bond dynamics of the biosensor, and external mechanical stiffness. Together, this work describes a new paradigm for detecting the effect of mechanical loads on FPs in cells and provides a method to visualize force-induced changes in protein structure/function independent of secondary probe binding, which can be leveraged to study mechanosensitive processes.

These findings demonstrate that genetically encoded FRET-based MTS can operate in two modes, measuring mechanotransmission in the absence of FP mechanical switching or measuring mechanosensing in the presence of FP mechanical switching. In the mechanotransmission mode, i.e. the original design intention, load magnitude is inferred from FRET efficiency, specifically using the eff-force relationship for calibrated MTSs^10,15^. If FP mechanical switching occurs, the calibration becomes inaccurate, with the severity of the effect depending on the amount of mechanical switching present in the donor and/or acceptor. To this end, our work provides a quality control for accurate measurements by MTSs in mechanotransmission mode. The first step is to rule out factors not related to FP mechanical switching by applying the criteria established by Coullomb et al.^29^ to the unloaded control, i.e. the unloaded control should have the expected *S*∼0.5 and unloaded *E* value. Secondly, using the framework developed here, it should be shown that the MTS maintains *S*∼0.5 across the full range of *E* values to be validated for quantitative measurements of load magnitude using the eff-force relationship. However, in the presence of most deviations in *S*, a reduction in *E* is still indicative of loading, meaning relative comparisons of mechanotransmission are still possible (although contributions from load magnitude versus load duration are not separable).

Furthermore, MTSs can also operate in a mechanosensing mode that leverages FP mechanical switching. In this mode, the MTS provides a readout of force-induced changes in FP function in response to the loads across the protein-of-interest. We note that FP mechanical switching depends on both the load magnitude and dynamics (e.g. load duration or rate), which is evidenced by the kinetics of GFP mechanical switching from in vitro experiments^17^, and which we incorporated into our mathematical model of this process inside MTSs. Thus, FP mechanical switching is not another means to measure force magnitude. Instead, it is an approach to assess whether mechanosensing is supported in a specific load-bearing protein and biological context. Compared to other imaging-based techniques for mechanosensing^11-13^, which rely on labeling unfolded protein domains with secondary binding probes, our approach based on FP mechanical switching has the advantage of being independent of secondary probe binding. Additionally, our approach is suited for dynamic measurements of mechanosensing in live cells. The main disadvantage is that the readout is not the endogenous domain itself, making the mechanical switching properties of the FPs an important design consideration. Our modeling indicated that FP mechanical switching is easiest to detect when restricted predominantly to one of the two FPs (Note S1). Choosing this designated FP to have certain mechanical switching properties might enable mimicking a variety of endogenous mechanosensitive domains.

A broad consequence of this work is that the identity of FPs (and thus mechanical properties), as well as their placement (with respect to the line of loading), are critical design elements of any probe or biosensor for a load-bearing protein. Even when different FPs or placements are tolerated biologically, altering either aspect could have unintended consequences on the performance of the sensor or probe when it is placed under mechanical load. Therefore, unless the lack of mechanical switching has been demonstrated in both systems, results from MTSs for the same protein but with different FPs may not be directly comparable even when using FRET efficiency measurements. This is one potential explanation for discrepancies in sets of vinculin^33,39^ or E-cadherin^40,41^ tension measurements, each based on MTSs with two different FP pairs. Additionally, the consequences of FP identity and placement apply to other probes and sensors that put FPs in the line of mechanical loading but were not explicitly designed to be force-sensitive. One example is synthetic cross-linkers harboring a single FP between binding domains, such as the membrane-actin cortex linkers that have been previously used to image membrane proximal actin or manipulate membrane-cortex adhesion^42,43^. Another example is conformation sensors that were designed for reporting the relief of head-tail inhibition (also commonly referred to as autoinhibition), which is often found in load-bearing proteins^44^. This includes the vinculin conformation sensor (VinCS), in which one FP is inserted in the line of force between the head and tail and the other is outside the line of force at the c-terminus. For instance, multiple versions of VinCS exist that use different FP FRET pairs and change if the donor or acceptor FP is placed in the loaded versus unloaded positions in the sensor^30,45,46^, suggesting that each version could be affected by FP mechanical switching differently.

An important direction from this work will be screening and engineering new FPs with different mechanical switching properties. Here, we found that mVenus mechanical switching was dominant over mTFP1, indicating that FPs can have different mechanical switching sensitivities *in cellulo*. Consistent with this, different mechanical stabilities have also been reported in vitro for GFP and EYFP, two FPs derived from the same species^22,23^. For MTSs in mechanotransmission mode and other probes and sensors with FPs in the line of force, proper sensor function requires the use of mechanically stable FPs. It was previously shown that both YPet and mCherry can be subjected to considerable load magnitudes and durations (e.g. 24 pN for >5 min) without unfolding^47^. This suggests that these could be well suited for sensors requiring mechanically stable FPs, like MTSs in mechanotransmission mode. However, we suggest that their mechanical switching properties be tested *in cellulo* when used in biosensors, as the insertion of FPs into various proteins can affect the function of the FP^24^. Furthermore, the possibility for FP mechanical switching should be evaluated when previously validated sensors are used in different subcellular structures or cell types, as little is known about the loading magnitudes, durations, and rates in various contexts. On the other hand, the development of new MTSs in mechanosensing mode will rely on FPs with various mechanical switching sensitivities. Finding FPs with desired mechanical switching properties would enable sensors that function as detectors of mechanosensing in response to a plethora of dynamic loading conditions and for pairing with specific endogenous mechanosensitive domains.

Determining the relationships between FP mechanical switching and force-sensitive conformational changes in endogenous mechanosensitive domains will also be a key topic for the future.

This work describes a new framework for assessing FP mechanical stability and provides a means of directly probing force-sensitive protein function *in cellulo*. First, it provides a quality control that will immediately improve the development and application of genetically-encoded FRET-based MTSs designed to quantitatively measure the first step in mechanosensitive signaling, mechanotransmission. Second, it provides a new approach leveraging FP mechanical switching inside MTSs to probe another key step in mechanosensitive signaling, mechanosensing. Together, this and existing tools form an overlapping continuum for probing the multi-step molecular mechanisms of mechanosensitive signaling.

## LIMITATIONS OF THE STUDY

When considering the application or extension of our framework and experimental methodology for detecting FP mechanical switching *in cellulo*, limitations pertaining to the modelling of the photophysical properties of FPs during mechanical switching, imaging modality, and MTS design should be considered.

Firstly, in our mathematical model, we made assumptions about the effect of FP mechanical switching on the photophysical properties of FPs to determine the signal contribution for each sensor state in three channel FRET measurements. To our knowledge, forced-induced changes in the excitation or emission wavelengths of FPs have not been described. Therefore, we assumed that donor FPs that have undergone mechanical switching cannot be excited by any excitation light in the optical system, and that acceptor FPs that have undergone mechanical switching cannot be excited by any excitation light in the optical system and also cannot accept energy from donor FPs. If this assumption is not met, a substantially more complex formalism is needed. Other, less critical assumptions associated with the mathematical model are detailed in Note S1.

Secondly, we focused on sensitized emission for the FRET imaging modality because both the acceptor and donor are readily observable, its ease of use and low cost, standard calibration methodologies to measure FRET efficiency, and, most importantly, existing frameworks for analyzing FP stoichiometry^27-29^. The framework we developed is not immediately adaptable to other FRET modalities, such as FLIM, which are also used to image MTSs^15^. However, we suggest that sensitized emission FRET can be used for quality control of new and existing MTSs designed for mechanotransmission mode before using them in other imaging modalities. For instance, extensive donor FP mechanical switching could bias FLIM-FRET results and would not be detected by most standard FLIM-FRET imaging protocols.

Lastly, we applied our framework to TSMod^30^, which contains a calibrated unstructured tension sensing element. The framework here can be applied immediately to TSMod in other endogenous and synthetic proteins. By modifying the eff-force relationship, the framework here can be applied immediately to other calibrated unstructured tension sensing elements (e.g. repeats of GGSGGS), whose eff-force relationships are loading rate-independent and have no hysteresis^10^. Additionally, the framework is adaptable to calibrated structured tension sensing elements that operate at equilibrium, whose eff- force relationships are also loading rate-independent and have no hysteresis, such as those based on a single rapid unfolding transition like HP35 and HP35st^15^. In contrast, the framework is not readily modifiable for structured tension sensing elements exhibiting loading rate-dependence or hysteresis.

## Methods

### Generation of DNA constructs

Construction of pcDNA3.1-VinTS, pcDNA3.1-VinTS-I997A, pcDNA3.1-VinTS-E1015A, pcDNA3.1-VinTS-E1021A, pcDNA3.1-VinTS-E1015A-E1021A, pRRL-VinTS, and pRRL-VinTS-E1015A-E1021A have been described previously^30,33,34^. ABDTS is comprised of F-tractin (actin-binding peptide from rat neuronal inositol 1,4,5-triphosphate 3-kinase A), followed by a 9 amino acid linker (GLALPVATM, hereafter called “Linker 1”), the original TSMod, an 11 amino acid linker (GGSGSDPPVAT, hereafter called “Linker 2”), and a second F-tractin. F-tractin and Linker 1 were derived from pEGFP-C1 F-tractin-EGFP (Addgene Plasmid #5847)^48^. The original TSMod was derived from pcDNA3.1-TSMod (Addgene Plasmid #26021)^30^. Linker 2 (GGSGSDPPVAT) was used previously in another construct containing F-tractin^49^. Gibson Assembly (with Gibson Assembly Master Mix, E2611S; NEB, Ipswich, MA) was used to generate pcDNA3.1-ABDTS from pcDNA3.1 vector digested with EcoRI-HF (Cat #: R310S; NEB)/NotI-HF (Cat #: R3189S; NEB,) and the following three fragments containing complementary regions: (1) F-tractin and Linker 1, (2) TSMod, and (3) Linker 2 and F-tractin. The fragment containing F-tractin and Linker 1 was generated by Polymerase Chain Reaction (PCR) using pEGFP-C1 F-tractin-EGFP and the oligonucleotide primer sequences (5’ to 3’) ccactagtccagtgtggtggATGGCGCGACCACGGGGC (forward) and tgctcaccatCATGGTGGCGACCGGTAGCG (reverse). The fragment containing TSmod was generated by PCR using pcDNA3.1-TSMod and the oligonucleotide primer sequences (5’ to 3’) cgccaccatgATGGTGAGCAAGGGCGAG (forward) and cgctgccgccCTTGTACAGCTCGTCCATGC (reverse). The fragment containing Linker 2 and F-tractin was generated by PCR using pEGFP-C1 F-tractin-EGFP and the oligonucleotide primer sequences (5’ to 3’) gctgtacaagggcggcagcggcagcgatccccccgtggccaccATGGCGCGACCACGGGGC (forward) and cgggccctctagactcgagcTTACCCTGCGGCCGCTGC (reverse). ABDTL is comprised of F-tractin, Linker 1, and the original TSMod, i.e. only the part of ABDTS before Linker 2 and the second F-tractin. pcDNA3.1-ABDTL was generated via PCR from pcDNA3.1-ABDTS and inserted into pcDNA3.1 via BamHI-HF (Cat #: R3136S; NEB)/EcoRI-HF (Cat #: R310S; NEB) digestion and subsequent ligation (Cat #: M0202S; NEB). All newly generated constructs were verified by DNA sequencing (GENEWIZ from Azenta).

### Cell Culture and Expression of DNA constructs

Vinculin -/- MEFs (kindly provided by Dr. Ben Fabry and Dr. Wolfgang H. Goldmann, Friedrich-Alexander-Universität Erlangen-Nürnberg)^50^ were maintained at 37 °C in a humidified 5% CO_2_ atmosphere in Dulbecco′s Modified Eagle′s Medium (DMEM) high glucose with sodium pyruvate (D6429; Sigma Aldrich, St. Louis, MO) supplemented with 10% FBS (HyClone SH30071.03; Cytivia, Marlborough, MA), 1% v/v non-essential amino acids (11140050; Thermo Fisher Scientific, Waltham, MA), and 1% v/v penicillin-streptomycin solution (15140122; Thermo Fisher). The generation of cell lines stably expressing VinTS and VinTS-E1015A-E1021A via lentiviral transduction were described previously^33,34,51^. NIH-3T3 cells were obtained from ATCC (CRL-1658) and maintained in the same conditions, using the same media formulation except for the omission of non-essential amino acids. For transient expression of ABDTS or ABDTL, NIH 3T3s were transfected at 50–75% confluence in 6-well tissue culture plates (25-105; Genesee Scientific, El Cajon, CA) using Lipofectamine 2000 (11668019; Thermo Fisher) and OptiMEM (31985070; Thermo Fisher) following the manufacturer’s instructions. For imaging, glass bottom dishes with no. 1.5 coverslips (FD35-100; World Precision Instruments, Sarasota, FL) or no. 1.5 glass coverslips mounted in reusable metal dishes (30-1313-03192; Bioptechs, Butler, PA) were incubated with 10 µg/ml fibronectin (33016015; Thermo Fisher) in PBS at room temperature for 1 hour and rinsed once with PBS. Approximately 3000 cells/cm ^2^ were plated on the dishes. Cells were incubated for 4 hours to enable sufficient spreading.

### Analysis of Previous VinTS Data Sets

Analyses were conducted on previously obtained three-channel FRET images of Vinculin -/- MEFs expressing VinTS or VinTS-I997A on FN-coated glass that were part of the data in the study by Rothenberg et al. ^33^, and the three-channel FRET images of Vinculin -/- MEFs expressing VinTS, VinTS-I997A, VinTS-E1015A, VinTS-E1021A, or VinTS-E1015A-E1021A on FN-coated glass that were part of the data in the study by Chirasani, Khan, Malavade, et al. ^34^.

### Fixation & Phalloidin Labeling of Actin

For fixation, cells were washed twice with PBS, fixed with 4% v/v EM-grade paraformaldehyde (Cat #: 15700; Electron Microscopy Sciences, Hatfield, PA) in PBS for 10 minutes and then rinsed with PBS. To label actin, cells were permeabilized with 0.1% Triton-X in PBS for 5 min, blocked with 2% bovine serum albumin (BSA, A7906-100G; Sigma-Aldrich) in PBS for 30 min, treated with Alexa Fluor 647-conjugated phalloidin (A22287; Thermo Fisher) at a 1:100 dilution for 60 min, and then rinsed three times with PBS. Cells were imaged in PBS.

### Pharmacological Inhibitors

Cells were seeded on fibronectin-coated glass coverslips as described above. A solution of 1 µM Latrunculin A (L5163; Sigma-Aldrich) were prepared from a 2 mM stock solution in DMSO (D2650; Sigma-Aldrich) by dilution in complete growth medium. Vehicle only controls were conducted with DMSO in complete growth medium. Cells were treated for 15 minutes at 37°C prior to fixation.

### Imaging of FRET-based Sensors and Immunofluorescence

An Olympus inverted fluorescent microscope (IX83; Olympus, Tokyo, Japan) was used to image samples as described previously^33^. Images were acquired at 60x magnification (UPlanSApo 60X/NA1.35 Objective, Olympus) and illuminated by a Lambda LS equipped with a 300W ozone-free xenon bulb (Sutter Instrument, Novato, CA). The images were captured using a sCMOS ORCA-Flash4.0 V2 camera (Hamamatsu Photonics, Hamamatsu, Japan). The FRET images were acquired using a custom filter set comprised of an mTFP1 excitation filter (ET450/30x; Chroma Technology Corp, Bellows Falls, VT), mTFP1 emission filter (FF02-485/20-25, Semrock, Rochester, NY), Venus excitation filter (ET514/10x; Chroma Technology Corp), Venus emission filter (FF01-571/72; Semrock), and dichroic mirror (T450/514rpc; Chroma Technology Corp). For sensitized emission FRET microscopy, three images are acquired to calculate FRET efficiency^51^. These include imaging the acceptor (I_AA_, Venus excitation, Venus emission), FRET (I_DA_, mTFP1 excitation, Venus emission), and donor (I_DD_, mTFP1 excitation, mTFP1 emission). For immunofluorescent imaging, we utilized the DA/FI/TR/Cy5-4X4 M-C Brightline Sedat filter set (Semrock) and the associated dichroic mirror (FF410/504/582/669-Di01). The motorized filter wheels (Lambda 10-3; Sutter Instrument), automated stage (H117EIX3; Prior Scientific, Rockland, MA), and image acquisition were controlled through MetaMorph Advanced software (Molecular Devices, San Jose, CA). Differential interference contrast (DIC) images were acquired at 60x magnification using a polarizer (IX-LWPO, Olympus) mounted above the condenser, an adjustable DIC slider (U-DICT, Olympus) containing one of the Nomarski prisms, and a mirror unit equipped with the analyzer (IX3-FDICT, Olympus). The Nomarski prism within the DIC slider was adjusted to achieve optimum contrast prior to each experiment.

For live cell imaging of NIH-3T3 cells, growth media was replaced with DMEMgfp-2 live cell visualization media (MC102; Sapphire North America, Ann Arbor, MI), supplemented with 10% FBS, 30 minutes before imaging. For live cell imaging of MEFs, growth media was replaced with the same live cell visualization media and supplemented with 10% FBS and 1% NEAA, 30 minutes before imaging. A constant temperature was maintained across the sample using an objective heater (Objective Heater Medium 150819-13; Bioptechs, Butler, PA) in conjunction with a stage and lid heater (Stable Z System 403-1926; Bioptechs). A humidified CO_2_ perfusion system (130708; Bioptechs) was used to maintain a stable pH. All components were brought to thermal equilibrium prior to imaging.

### Calculation of FRET Efficiency and Stoichiometry from Sensitized Emission

FRET was detected through measurement of sensitized emission^27^ and calculated using custom written code in MATLAB (Mathworks)^51^. All analyses were conducted on a pixel-by-pixel basis. Prior to FRET calculations, all images were first corrected for dark current, uneven illumination, background intensity, and three-dimensional offsets caused by chromatic aberrations and minute hardware misalignments (registration) as previously described^28^. Spectral bleedthrough coefficients were determined through FRET-imaging of cells expressing only donor or only acceptor FPs. The donor bleedthrough coefficient (*dbt*) was calculated for mTFP1 as:

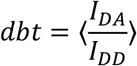

where *I*_*DD*_ is the intensity in the FRET-channel, *I*_*DD*_ is the intensity in the donor-channel, and data were binned by donor-channel intensity. Similarly, the acceptor bleedthrough coefficient (*abt*) was calculated for Venus (A206K) as:

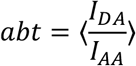

where *I*_*DD*_ is the intensity in the acceptor-channel, and data were binned by acceptor-channel intensity. For the mTFP1-Venus (A206K) FP pair on our microscope setup, the cross-talk between donor and acceptor channels (signal from donor in acceptor channel and vice-versa) was determined to be negligeable. To correct for spectral bleedthrough in experimental data, pixel-by-pixel FRET corrections were performed according to the equation:

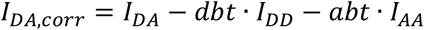

where *II*_*DA, corr*_ is the corrected FRET intensity (also defined in the literature as *FF*_*cc*_). After bleedthrough correction, FRET efficiency was calculated. Through imaging donor-acceptor fusion constructs of differing, but constant, FRET efficiencies, it is possible to calculate two proportionality constants that enable the calculation of FRET efficiencies for a given FRET pair^27^. These proportionality constants are *G*:

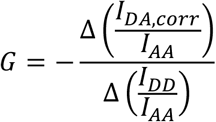

where Δ indicates the change between two donor-acceptor fusion proteins, and *kk*:

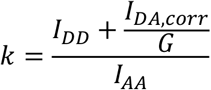

Using published methods^28^, the calibration factors were experimentally determined for mTFP1 and Venus (A206K). With these two proportionality constants, it is possible to calculate both the apparent FRET efficiency (*E*_*app*_):

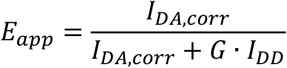

and FP stoichiometry^29^ (*S*_*app*_):

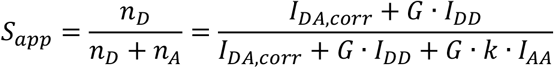

The calibration constants *G* and *k* were monitored over the course of this work to control for changes in lamp and filter performance. Note, these formulas are equivalent to the formulas from Coulomb et al.^29^ with *α*^*BT*^ = *dbt, δδ*^*DE*^ = *abt, γ*^*M*^ = *G*, and 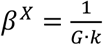:

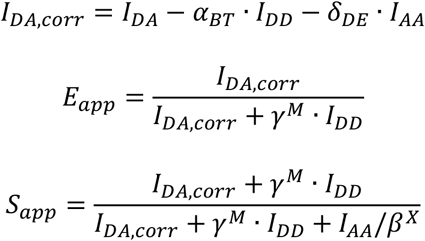

Note that all experimental FRET measurements are inherently apparent FRET efficiency (*E*_*app*_) and FP stoichiometry (*S*_*app*_), but are indicated for experimental data in this work using *E* and *S* without the subscript to match previous conventions^16,27,28,31^.

To analyze *E* and *S*, the following approaches were used to generate analysis masks. For ABDTS and ABDTL, analysis masks were constructed for whole cells. Cell masks were manually drawn using DIC images (for live imaging) or phalloidin-labeled actin images (for fixed imaging). Cell masks were refined by excluding pixels in which the intensity of both *I*_*AA*_ and *I*_*DD*_ was below a small threshold value (exclude px if *I*_*DD*_ < 50 and *I*_*AA*_ < 50). This condition preserves regions containing signal from both or a single FP. For VinTS variants, analysis masks contained all pixels in the FAs of single cells. Segmentation of FAs was done as previously reported using a water-based algorithm^51^. Segmentation and manual cell mask generation for VinTS variants were performed using the acceptor channel, which is independent of FRET, as previously described^33^.

For ES-histogram analyses, all pixels in the above-described analysis mask were used. *E* outside [0.000, 0.500] and *S* outside [0.000, 1.000] were set to the nearest limit. ES-histograms were constructed with fixed bin widths. For single cells, pixel counts were plotted, and bin widths of 0.005 for *E* and 0.010 for *S* were used. For cell populations, histograms show the average of the relative fraction of each bin across cells in the indicated group, which gives each cell in the population an equal waiting regardless of its size (number of pixels in analysis mask). Here, bin widths of 0.025 for *E* and 0.050 for *S* were used. To quantitatively compare the extent of acceptor FP mechanical switching between groups or conditions, the fraction of pixels in a larger bin containing low *E* values and high *S* values was quantified for each cell. Specifically, we quantified the fraction of pixels in each cell with *E* < 0.15 and *S* > 0.60.

### Preparation of Methacrylated Coverslips and Hydrophobic Glass Substrate

18×18 No. 2 glass coverslips (48368-040; VWR, Radnor, PA) were exposed to air plasma for 5 minutes. Immediately before addition to glass coverslips, 500 μL of glacial acetic acid (ACROS Organics AC124040010; Thermo Fisher) was added to a solution of 3-(Trimethoxysilyl) propyl methacrylate (M6514-25ML; Sigma-Aldrich) in ethanol (E7023-500ML; Sigma-Aldrich; 30 μL in 9.5 mL). This solution was applied to the coverslips and incubated for 5 minutes. The coverslips were then washed twice with ethanol and dried via compressed air. Coverslips were prepared < 1h before fabrication of polyacrylamide gels.

A hydrophobic glass substrate was prepared using by first rinsing with DI H_2_O, then 70% ethanol in DI H_2_O (Sigma-Aldrich), and finally 100% isopropyl-alcohol (BDH-11334LP; VWR International). This was dried via compressed air. Rain-X® Original Glass Water Repellent (ITW Global Brands, Houston, TX) was applied to the pre-cleaned glass substrate according to the manufacturer’s instructions.

### Preparation, Imaging, and Mechanical Testing of PA Gels

Polyacrylamide (PA) gels were fabricated using a methodology to improve ECM protein attachment described previously^52^. Varying ratios of 40% acrylamide (1610140; Bio-Rad, Hercules, CA) and 2% bis-acrylamide solutions (1610142; Bio-Rad) were mixed with DI H_2_O at the ratio shown in Table 1. This solution was de-gassed for 10 minutes via sonication under vacuum. The de-gassed solution was mixed with a 1.5M solution of 2-aminoethyl methacrylate (516155-5G; Sigma-Aldrich) in DI H_2_O at a ratio of 1:100. This solution was passed through a 0.2 μm syringe filter (VWR) to remove any particulate. This solution was mixed with a 100 mg/mL solution of Irgacure-2959 (410896-10G; Sigma-Aldrich) in methanol (VWR) at a ratio of 1:100. 240 μL of this solution was pipetted onto a hydrophobic glass substrate (see above). A methacrylated coverslip (see above) was inverted onto the droplet of PA solution and exposed to UV light for 15 min. Following polymerization, the glass substrate was flooded with PBS and the gel-attached coverslips were gently removed. Gels were placed in a 6-well plate and washed twice with PBS. Following washing, a working solution of 0.83 mg/mL solution of sulfosuccinimidyl 6-(4′-azido-2′-nitrophenylamino)hexanoate (sulfo-SANPAH, 22589; Thermo Fisher) in PBS was prepared from a stock solution of 83 mg/mL sulfo-SANPAH in DMSO (Sigma-Aldrich). The working solution was pipetted onto the surface of each gel, ensuring complete coverage. The gels were then exposed to UV light for 5 min. The gels were rinsed twice with PBS and covered with a 10 μg/mL solution of fibronectin in PBS (for gels to be used for cell attachment) or 10 μg/mL BSA in PBS (to reduce non-specific adhesion for gels to be used in mechanical testing) prior to incubation overnight at 4 °C.

**Table 1:**
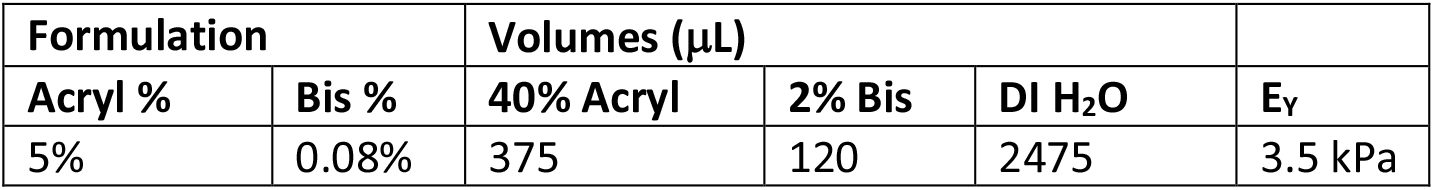
Formulation, Solution Volumes, and Mechanical Testing for PA Gel.

| Formulation | | Volumes (μL) | | | |
| --- | --- | --- | --- | --- | --- |
| Acryl % | Bis % | 40% Acryl | 2% Bis | DI H<sub>2</sub>O | E<sub>y</sub> |
| 5% | 0.08% | 375 | 120 | 2475 | 3.5 kPa |

For imaging, gels were kept in a 6-well dish. The fibronectin in PBS solution was changed to full MEF growth media prior to seeding cells. MEFs (stably expressing VinTS or VinTS-E1015A-E1021A) were seeded at approximately 5700 cells/cm ^2^ and allowed to adhere for 24h prior to fixation. Following fixation, the gels were stored in PBS until imaging. No. 1.5 glass coverslips were mounted in reusable metal dishes (Bioptechs, Butler, PA). A 50 μL droplet of PBS was added to the coverslip and the gel was carefully inverted onto the droplet. Imaging was performed in this configuration.

Mechanical testing of PA gels was performed via atomic force microscopy (Asylum Research MFP 3D, Santa Barbara, CA). Hydrogels were immersed in PBS and indented with a spherical cantilever (10 μm radius, borosilicate glass, Novascan Technologies, Ames, IA) with a spring constant of 188.39 pN/nm at an indentation speed of 0.8 μm/s. Young’s modulus was determined via Hertzian model fit.

### Statistics

Statistical analyses were performed using JMP Pro (SAS, Cary, NC) software. Normality of each dataset was assessed using Q-Q plots and Shapiro-Wilks test. For normal datasets, comparisons of data with equal variances, as determined with Levene’s test, were analyzed with an ANOVA and, if necessary, Tukey’s Honest Significant Difference (HSD) tests. Datasets with unequal variances were analyzed with a non-parametric Welch’s ANOVA and, if necessary, the Steel-Dwass multiple comparisons test. For non-normal datasets, comparisons of data were analyzed with a Kruskal-Wallis test and, if necessary, Steel-Dwass multiple comparisons test. A p value of p<0.05 was considered statistically significant. In figures, a single asterisk (*), double asterisk (**), triple asterisk (***), and quadruple asterisk (****) indicate p-values less than 0.05, 0.01, 0.001, and 0.0001, respectively, and ns indicates a p-value greater than or equal to 0.05. Cells were treated as independent observations. Sample sizes for each experiment can be found in the figure legends. Where used, standard box plots were created using JMP Pro, where the bottom and top of the box indicate the first and third quartiles, respectively, the middle line indicates the median, the whiskers extend to the outermost data points below the first quartile and above the third quartile that are within 1.5 times the interquartile range, and data outside the whiskers are indicated as points.

### Mathematical Models of FP Mechanical Switching in MTS

The formulation and implementation of the mathematical models of FP mechanical switching in load-bearing proteins and MTSs are provided in Note S1.

### Steered Molecular Dynamics Simulations of FPs

The methods for SMD simulations of FPs are provided in Note S2.

## Resource Availability

### Lead Contact

Further information and requests for resources and reagents should be directed to and will be fulfilled by the lead contact, Brenton D. Hoffman.

### Materials availability

Plasmids generated in this study will be shared up request to the lead contact, until made publicly available on Addgene.

### Data and code availability

1. All data reported in this paper will be shared by the lead contact upon request.
2. MATLAB codes used to perform image pre-processing (https://gitlab.oit.duke.edu/HoffmanLab-Public/image-preprocessing) and three channel sensitized emission FRET image analysis (https://gitlab.oit.duke.edu/HoffmanLab-Public/fret-analysis) are publicly available on GitLab. MATLAB codes used to analyze and simulate the mathematical models of FP mechanical switching will be made publicly available on GitLab before publication.
3. Any additional information required to reanalyze the data reported in this paper is available from the lead contact upon request.

## Acknowledgements and Funding

We thank Dr. Stefano Di Talia and Dr. Victoria Deneke (Duke University) for their assistance with molecular cloning to generate ABDTS and ABDTL constructs. We also acknowledge support from the Duke Compute Cluster. Images of molecular dynamics simulations in this work were made with VMD & NAMD software support. VMD & NAMD were developed with NIH support by the Theoretical and Computational Biophysics group at the Beckman Institute, University of Illinois at Urbana-Champaign. This research was supported by the National Institute of Health (1R01GM121739 to B.D.H. and 5T32GM008555-26 to K.L.C.), the National Science Foundation (GRFP DGE 2139754 to T.C.S. and GRFP DGE 2139754 to J.N.M.), and the National Cancer Institute K00 Fellowship (K00CA245789 to A.T.B.).

## Author Contributions

B.D.H. and J.L.W. conceived the project. K.L.C. and J.N.M. conducted experiments. T.C.S., T.R.H., K.L.C., and J.N.M. performed data analyses. T.C.S. designed and analyzed mathematical models. A.T.B. performed and analyzed SMD simulations. T.C.S., T.R.H., and B.D.H. wrote and edited the paper.

## Supplementary Information

Figures S1-S21, Captions for Movies S1-S3, Tables S1-S9, and Notes S1-S3 are included in the Supplementary Information document. Movies S1-S3 are provided separately.

## References

1 Wolfenson, H., Yang, B. & Sheetz, M. P. Steps in Mechanotransduction Pathways that Control Cell Morphology. Annu Rev Physiol 81, 585–605, doi:10.1146/annurev-physiol-021317-121245 (2019).

2 Hoffman, B. D., Grashoff, C. & Schwartz, M. A. Dynamic molecular processes mediate cellular mechanotransduction. Nature 475, 316–323, doi:10.1038/nature10316 (2011).

3 Hoffman, B. D. & Yap, A. S. Towards a Dynamic Understanding of Cadherin-Based Mechanobiology. Trends Cell Biol 25, 803–814, doi:10.1016/j.tcb.2015.09.008 (2015).

4 Elosegui-Artola, A., Trepat, X. & Roca-Cusachs, P. Control of Mechanotransduction by Molecular Clutch Dynamics. Trends Cell Biol 28, 356–367, doi:10.1016/j.tcb.2018.01.008 (2018).

5 Sharma, S., Subramani, S. & Popa, I. Does protein unfolding play a functional role in vivo? FEBS J 288, 1742–1758, doi:10.1111/febs.15508 (2021).

6 del Rio, A. et al. Stretching single talin rod molecules activates vinculin binding. Science 323, 638–641, doi:10.1126/science.1162912 (2009).

7 Yao, M. et al. Force-dependent conformational switch of alpha-catenin controls vinculin binding. Nat Commun 5, 4525, doi:10.1038/ncomms5525 (2014).

8 Alegre-Cebollada, J. Protein nanomechanics in biological context. Biophys Rev 13, 435–454, doi:10.1007/s12551-021-00822-9 (2021).

9 LaCroix, A. S., Rothenberg, K. E. & Hoffman, B. D. Molecular-Scale Tools for Studying Mechanotransduction. Annu Rev Biomed Eng 17, 287–316, doi:10.1146/annurev-bioeng-071114-040531 (2015).

10 Ham, T. R., Collins, K. L. & Hoffman, B. D. Molecular Tension Sensors: Moving Beyond Force. Curr Opin Biomed Eng 12, 83–94, doi:10.1016/j.cobme.2019.10.003 (2019).

11 Sawada, Y. et al. Force sensing by mechanical extension of the Src family kinase substrate p130Cas. Cell 127, 1015–1026, doi:10.1016/j.cell.2006.09.044 (2006).

12 Yonemura, S., Wada, Y., Watanabe, T., Nagafuchi, A. & Shibata, M. alpha-Catenin as a tension transducer that induces adherens junction development. Nat Cell Biol 12, 533–542, doi:10.1038/ncb2055 (2010).

13 Zhong, B. L., Vachharajani, V. T. & Dunn, A. R. Facile detection of mechanical forces across proteins in cells with STReTCh. Cell Rep Methods 2, 100278, doi:10.1016/j.crmeth.2022.100278 (2022).

14 Tao, A. et al. Identifying constitutive and context-specific molecular-tension-sensitive protein recruitment within focal adhesions. Dev Cell 58, 522–534 e527, doi:10.1016/j.devcel.2023.02.015 (2023).

15 Fischer, L. S., Rangarajan, S., Sadhanasatish, T. & Grashoff, C. Molecular Force Measurement with Tension Sensors. Annu Rev Biophys 50, 595–616, doi:10.1146/annurev-biophys-101920-064756 (2021).

16 LaCroix, A. S., Lynch, A. D., Berginski, M. E. & Hoffman, B. D. Tunable molecular tension sensors reveal extension-based control of vinculin loading. Elife 7, doi:10.7554/eLife.33927 (2018).

17 Ganim, Z. & Rief, M. Mechanically switching single-molecule fluorescence of GFP by unfolding and refolding. Proc Natl Acad Sci U S A 114, 11052–11056, doi:10.1073/pnas.1704937114 (2017).

18 Saeger, J., Hytonen, V. P., Klotzsch, E. & Vogel, V. GFP’s mechanical intermediate states. PLoS One 7, e46962, doi:10.1371/journal.pone.0046962 (2012).

19 Dudko, O. K., Hummer, G. & Szabo, A. Intrinsic rates and activation free energies from singlemolecule pulling experiments. Phys Rev Lett 96, 108101, doi:10.1103/PhysRevLett.96.108101 (2006).

20 Evans, E. Probing the relation between force--lifetime--and chemistry in single molecular bonds. Annu Rev Biophys Biomol Struct 30, 105–128, doi:10.1146/annurev.biophys.30.1.105 (2001).

21 Wang, Y., Yan, J. & Goult, B. T. Force-Dependent Binding Constants. Biochemistry 58, 4696–4709, doi:10.1021/acs.biochem.9b00453 (2019).

22 Dietz, H. & Rief, M. Exploring the energy landscape of GFP by single-molecule mechanical experiments. Proc Natl Acad Sci U S A 101, 16192–16197, doi:10.1073/pnas.0404549101 (2004).

23 Perez-Jimenez, R., Garcia-Manyes, S., Ainavarapu, S. R. & Fernandez, J. M. Mechanical unfolding pathways of the enhanced yellow fluorescent protein revealed by single molecule force spectroscopy. J Biol Chem 281, 40010–40014, doi:10.1074/jbc.M609890200 (2006).

24 Snapp, E. L. Fluorescent proteins: a cell biologist’s user guide. Trends Cell Biol 19, 649–655, doi:10.1016/j.tcb.2009.08.002 (2009).

25 Roca-Cusachs, P., Iskratsch, T. & Sheetz, M. P. Finding the weakest link: exploring integrinmediated mechanical molecular pathways. J Cell Sci 125, 3025–3038, doi:10.1242/jcs.095794 (2012).

26 Algar, W. R., Hildebrandt, N., Vogel, S. S. & Medintz, I. L. FRET as a biomolecular research tool - understanding its potential while avoiding pitfalls. Nat Methods 16, 815–829, doi:10.1038/s41592-019-0530-8 (2019).

27 Chen, H., Puhl, H. L., Koushik, S. V., Vogel, S. S. & Ikeda, S. R. Measurement of FRET efficiency and ratio of donor to acceptor concentration in living cells. Biophys J 91, L39–L41, doi:10.1529/biophysj.106.088773 (2006).

28 Gates, E. M., LaCroix, A. S., Rothenberg, K. E. & Hoffman, B. D. Improving Quality, Reproducibility, and Usability of FRET-Based Tension Sensors. Cytometry A 95, 201–213, doi:10.1002/cyto.a.23688 (2019).

29 Coullomb, A. et al. QuanTI-FRET: a framework for quantitative FRET measurements in living cells. Sci Rep 10, 6504, doi:10.1038/s41598-020-62924-w (2020).

30 Grashoff, C. et al. Measuring mechanical tension across vinculin reveals regulation of focal adhesion dynamics. Nature 466, 263–266, doi:10.1038/nature09198 (2010).

31 LaCroix, A. S., Rothenberg, K. E., Berginski, M. E., Urs, A. N. & Hoffman, B. D. Construction, imaging, and analysis of FRET-based tension sensors in living cells. Methods Cell Biol 125, 161–186, doi:10.1016/bs.mcb.2014.10.033 (2015).

32 Bays, J. L. & DeMali, K. A. Vinculin in cell-cell and cell-matrix adhesions. Cell Mol Life Sci 74, 2999–3009, doi:10.1007/s00018-017-2511-3 (2017).

33 Rothenberg, K. E., Scott, D. W., Christoforou, N. & Hoffman, B. D. Vinculin Force-Sensitive Dynamics at Focal Adhesions Enable Effective Directed Cell Migration. Biophys J 114, 1680–1694, doi:10.1016/j.bpj.2018.02.019 (2018).

34 Chirasani, V. R. et al. Molecular basis and cellular functions of vinculin-actin directional catch bonding. Nat Commun 14, 8300, doi:10.1038/s41467-023-43779-x (2023).

35 Shoyer, T. C. et al. Coupling during collective cell migration is controlled by a vinculin mechanochemical switch. Proc Natl Acad Sci U S A 120, e2316456120, doi:10.1073/pnas.2316456120 (2023).

36 Thompson, P. M. et al. Identification of an actin binding surface on vinculin that mediates mechanical cell and focal adhesion properties. Structure 22, 697–706, doi:10.1016/j.str.2014.03.002 (2014).

37 Rajagopalan, P., Marganski, W. A., Brown, X. Q. & Wong, J. Y. Direct comparison of the spread area, contractility, and migration of balb/c 3T3 fibroblasts adhered to fibronectin- and RGD-modified substrata. Biophys J 87, 2818–2827, doi:10.1529/biophysj.103.037218 (2004).

38 Huang, D. L., Bax, N. A., Buckley, C. D., Weis, W. I. & Dunn, A. R. Vinculin forms a directionally asymmetric catch bond with F-actin. Science 357, 703–706, doi:10.1126/science.aan2556 (2017).

39 Kanoldt, V. et al. Metavinculin modulates force transduction in cell adhesion sites. Nat Commun 11, 6403, doi:10.1038/s41467-020-20125-z (2020).

40 Cai, D. et al. Mechanical feedback through E-cadherin promotes direction sensing during collective cell migration. Cell 157, 1146–1159, doi:10.1016/j.cell.2014.03.045 (2014).

41 Eder, D., Basler, K. & Aegerter, C. M. Challenging FRET-based E-Cadherin force measurements in Drosophila. Sci Rep 7, 13692, doi:10.1038/s41598-017-14136-y (2017).

42 Bisaria, A., Hayer, A., Garbett, D., Cohen, D. & Meyer, T. Membrane-proximal F-actin restricts local membrane protrusions and directs cell migration. Science 368, 1205–1210, doi:10.1126/science.aay7794 (2020).

43 Bergert, M. et al. Cell Surface Mechanics Gate Embryonic Stem Cell Differentiation. Cell Stem Cell 28, 209–216 e204, doi:10.1016/j.stem.2020.10.017 (2021).

44 Khan, R. B. & Goult, B. T. Adhesions Assemble!-Autoinhibition as a Major Regulatory Mechanism of Integrin-Mediated Adhesion. Front Mol Biosci 6, 144, doi:10.3389/fmolb.2019.00144 (2019).

45 Chen, H., Cohen, D. M., Choudhury, D. M., Kioka, N. & Craig, S. W. Spatial distribution and functional significance of activated vinculin in living cells. J Cell Biol 169, 459–470, doi:10.1083/jcb.200410100 (2005).

46 Case, L. B. et al. Molecular mechanism of vinculin activation and nanoscale spatial organization in focal adhesions. Nat Cell Biol 17, 880–892, doi:10.1038/ncb3180 (2015).

47 Austen, K. et al. Extracellular rigidity sensing by talin isoform-specific mechanical linkages. Nat Cell Biol 17, 1597–1606, doi:10.1038/ncb3268 (2015).

48 Belin, B. J., Goins, L. M. & Mullins, R. D. Comparative analysis of tools for live cell imaging of actin network architecture. Bioarchitecture 4, 189–202, doi:10.1080/19490992.2014.1047714 (2014).

49 Spracklen, A. J., Fagan, T. N., Lovander, K. E. & Tootle, T. L. The pros and cons of common actin labeling tools for visualizing actin dynamics during Drosophila oogenesis. Dev Biol 393, 209–226, doi:10.1016/j.ydbio.2014.06.022 (2014).

50 Mierke, C. T. et al. Vinculin facilitates cell invasion into three-dimensional collagen matrices. J Biol Chem 285, 13121–13130, doi:10.1074/jbc.M109.087171 (2010).

51 Rothenberg, K. E., Neibart, S. S., LaCroix, A. S. & Hoffman, B. D. Controlling Cell Geometry Affects the Spatial Distribution of Load Across Vinculin. Cell Mol Bioeng 8, 364–382, doi:10.1007/s12195-015-0404-9 (2015).

52 Lee, S., Stanton, A. E., Tong, X. & Yang, F. Hydrogels with enhanced protein conjugation efficiency reveal stiffness-induced YAP localization in stem cells depends on biochemical cues. Biomaterials 202, 26–34, doi:10.1016/j.biomaterials.2019.02.021 (2019).

