## Supplemental Information for "Detection of Fluorescent Protein Mechanical Switching in Cellulo"

This document contains the following supplementary information:

- Supplemental Figures S1-S21
- Supplemental Movies S1-S3 Captions
- Supplemental Note S1 (containing Tables S1-S4)
- Supplemental Note S2
- Supplemental Note S3 (containing Tables S5-S9)
- Supplemental References

### Supplemental Figures and Legends

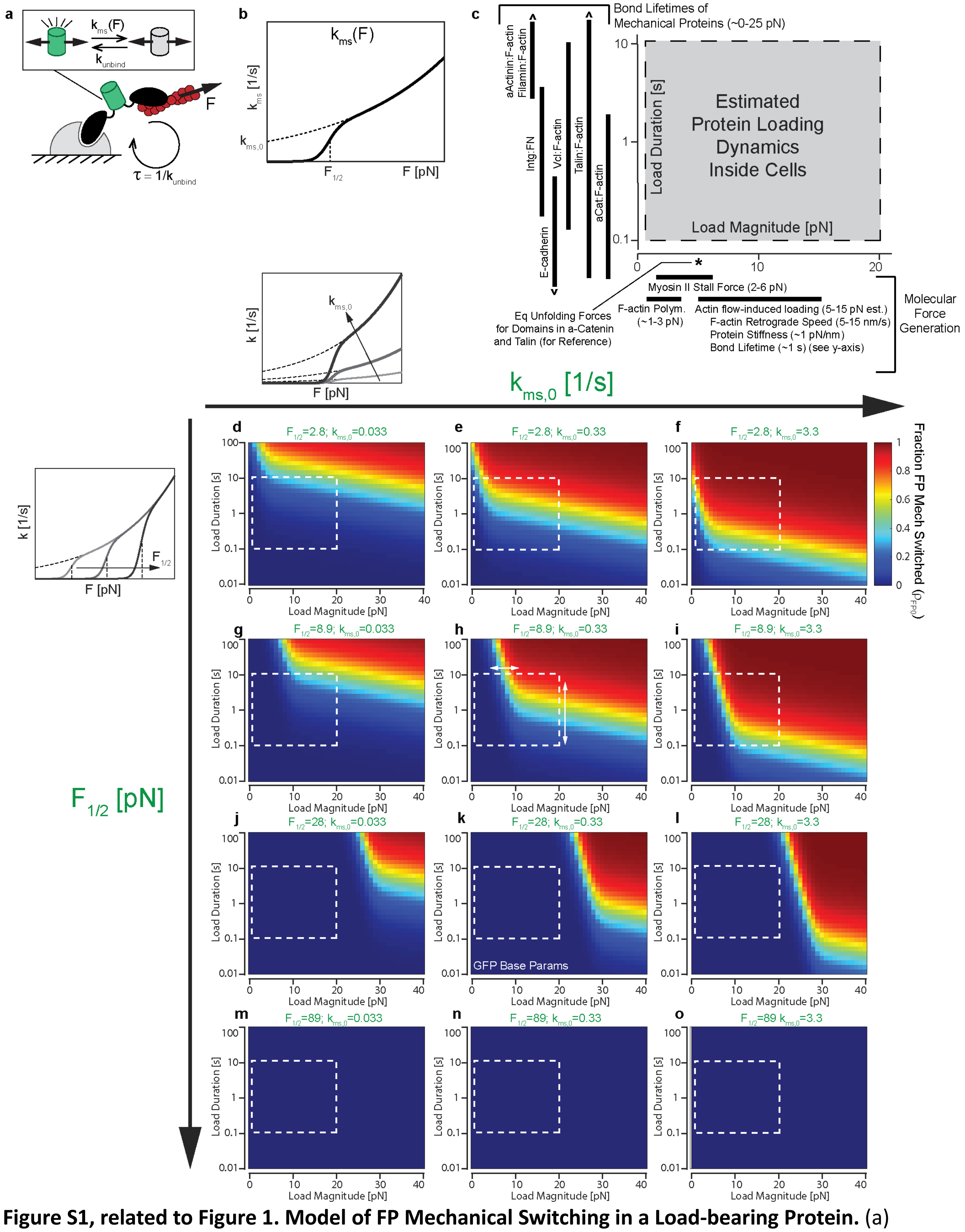


**Figure S1, related to Figure 1. Model of FP Mechanical Switching in a Load-bearing Protein.** (a) Schematic of model of a single FP integrated into a dynamic load-bearing protein. The protein is loaded by an actin structure to which it binds/unbinds. The FP is in the line of loading and can reversibly switch between functional (state 1) and non-functional (state 0) states in a force-sensitive manner. (b) Force-dependent rate constant for mechanical switching. (c) Estimated range of protein loading magnitude (x-axis) and duration (y-axis) inside cells, based on literature values for molecular force generation and bond lifetimes of mechanical proteins, respectively. See the text of Note S1 for citations. (d-o) Contour plots of the steady state fraction of FPs that have undergone mechanical switching ($\rho_{FP0}$) as a function of load magnitude ($F$) and load duration ($\tau$). Each plot corresponds to the indicated FP mechanical switching parameter combination ($F_{1/2}$ and $k_{MS,0}$). All other parameters were set to the base values indicated in Table S1.

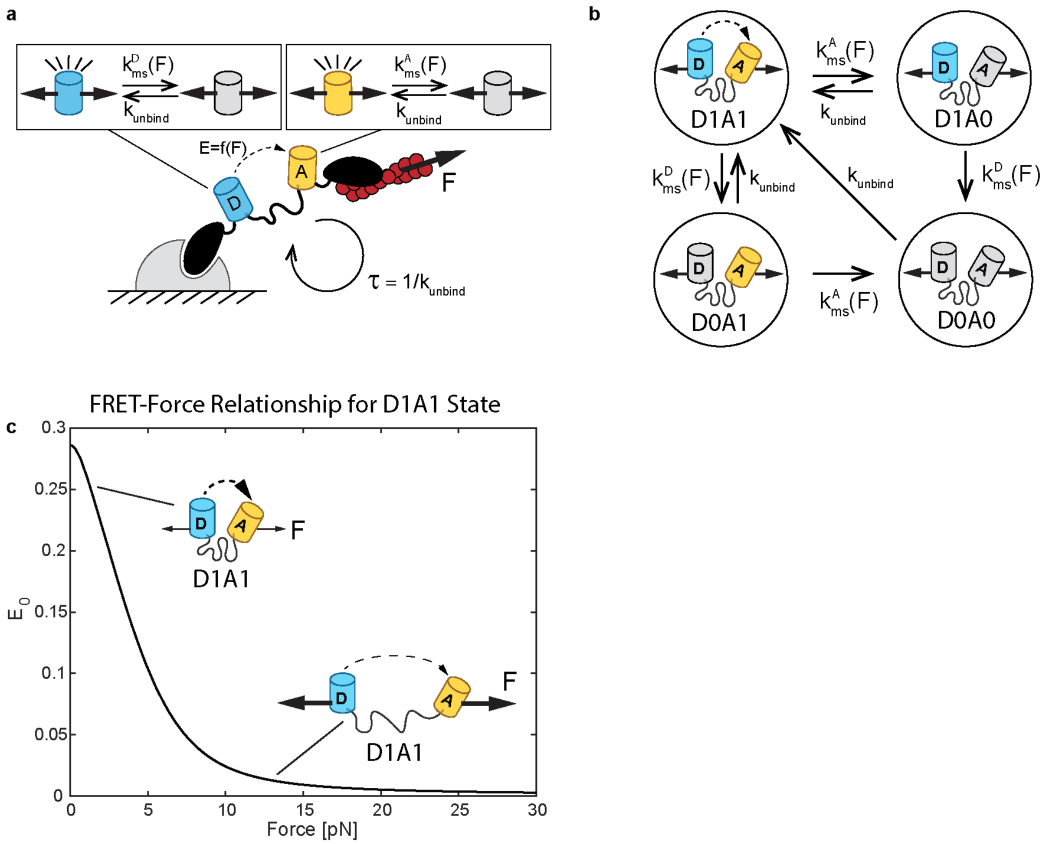


**Figure S2, related to Figure 1. Model Formulation of FP Mechanical Switching in MTS.** (a) Schematic of model of FP mechanical switching inside an MTS. The MTS is loaded by an actin structure to which it binds/unbinds. Donor and acceptor FPs are in the line of loading and can reversibly switch between functional (state 1) and non-functional (state 0) states in a force-sensitive manner. (b) Four possible MTS states based on the status of the donor and acceptor FP, with arrows indicating state transitions and the associated rate constant. (c) FRET-force relationship, $E_i = f(F_i)$, for an MTS in the D1A1 state.

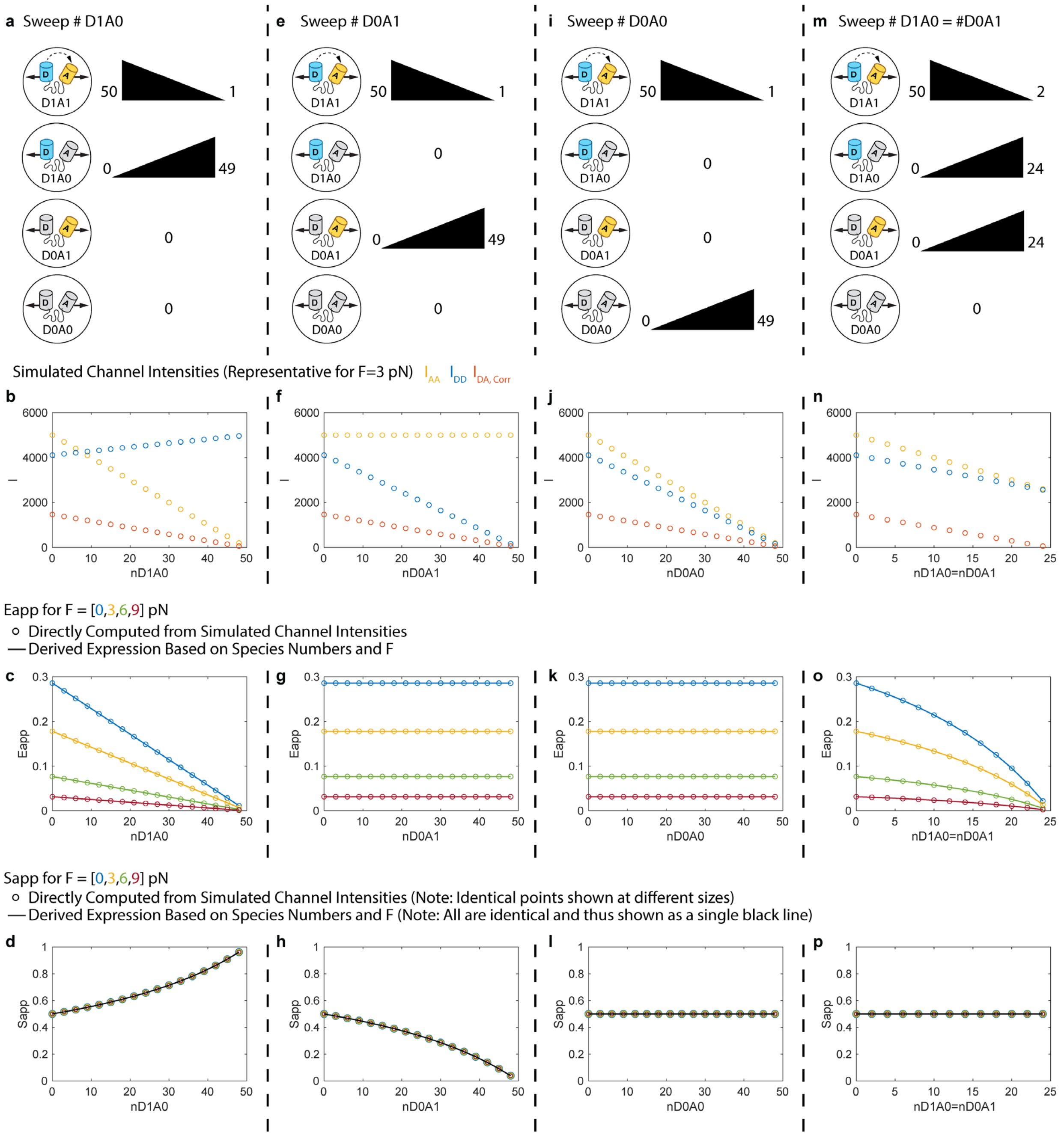


**Figure S3, related to Figure 1. Validation of Derived Expressions for $E_{app}$ and $S_{app}$ by Comparison to Direct Simulation of Channel Intensities.** The derived expressions for $E_{app}$ and $S_{app}$ (Note S1, Equations S18 and S19) for a population of MTSs comprised of a distribution of sensor states and all under the same force magnitude ($F$) were validated by comparison to direct computations from simulated channel intensities (Note S1, Equations S12-S17. (a-d) Effect of increasing the number of sensors in the D1A0 state ($n_{D1A0}$) from 0 to 49, with the remaining sensors in the D1A1 state (to a total of 50). (b) Intensities for the acceptor channel ($I_{AA}^{TOT}$; Equation S12), donor channel ($I_{DD}^{TOT}$; Equation S13), and corrected FRET channel ($I_{DA\,corr}^{TOT}$; Equation S15) versus $n_{D1A0}$ for $F$ of 3 pN. (c) Plot of $E_{app}$ computed directly from channel intensities (dots; Equation S16) or derived expression (line; Equation S18) versus $n_{D1A0}$ for four values of $F$. (d) Plot of $S_{app}$ computed directly from channel intensities (dots; Equation S17) or derived expression (line; Equation S19) versus $n_{D1A0}$ for four values of $F$. (e-h) Same set of plots for increasing the number of sensors in the D0A1 state ($n_{D0A1}$) from 0 to 49 with remaining sensors in the D1A1 state. (i-l) Same set of plots for increasing the number of sensors in the D0A0 state ($n_{D0A0}$) from 0 to 49 with remaining sensors in the D1A1 state. (m-q) Same set of plots for jointly increasing the number of sensors in the D1A0 and D0A1 states ($n_{D1A0} = n_{D0A1}$) from 0 to 24 with remaining sensors in the D1A1 state. Simulated channel intensities and direct computations from simulated channel intensities were performed with the following three-channel FRET parameters: $\widehat{\alpha_{BT}} = 0.75, \widehat{\delta^{DE}} = 0.25, \widehat{\gamma^{M}} = 1.65, \widehat{\beta^{X}} = 0.6061$, and $C_{AA} = 100$.

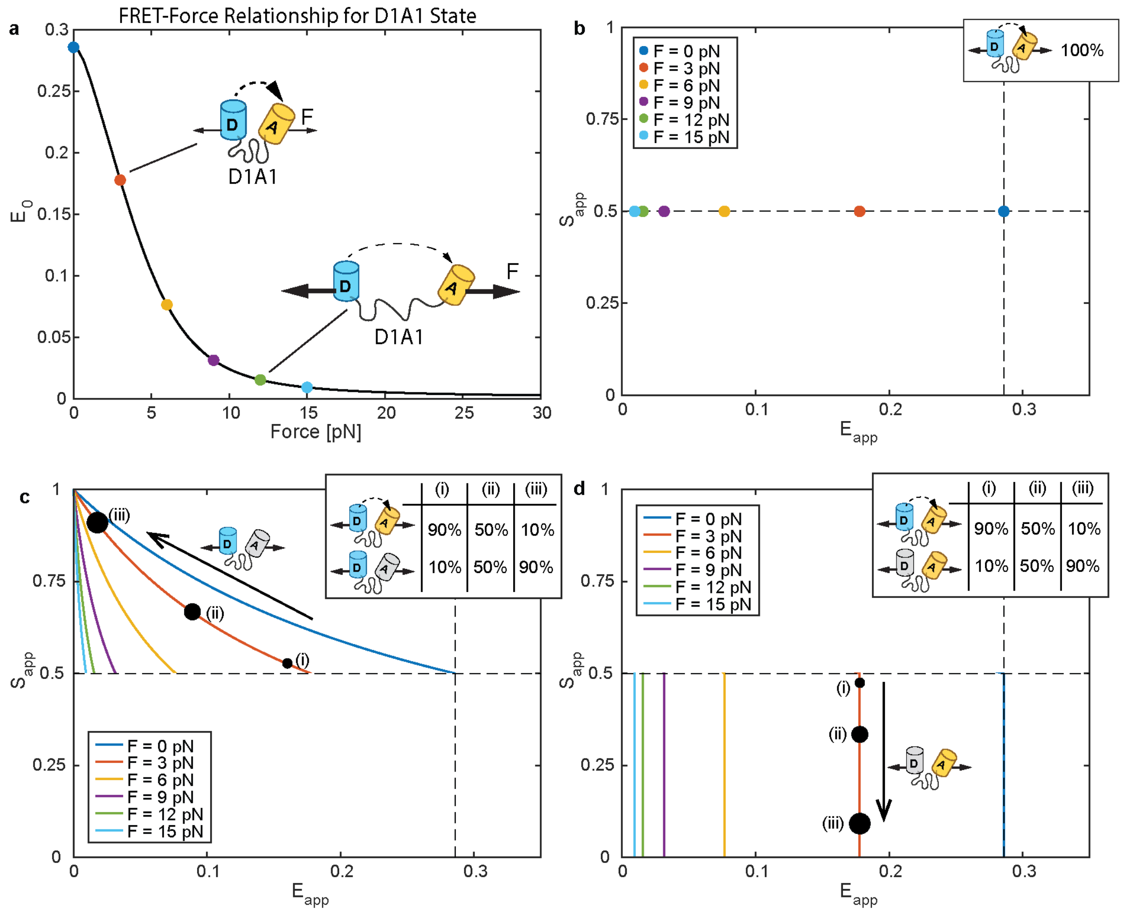


**Figure S4, related to Figure 1. Extension of ES-Histogram Framework to MTSs with FP Mechanical Switching.** (a) FRET-force relationship, $E_i = f(F_i)$, for an MTS in the D1A1 state, with dots indicating force magnitudes ($F$) of 0, 3, 6, 9, 12, and 15 pN. These force magnitudes are also indicated in the subsequent panels. (b) ES-Histogram for MTS ensemble comprised of 100% sensors in the D1A1 state and all sensors subject to the indicated $F$. (c) ($E_{app}$, $S_{app}$)-curves ("tension isoclines") for MTS ensembles comprised of various amounts of sensors in the D1A1 and D1A0 state (i.e. varying levels of acceptor mechanical switching only) with all sensors subject to the indicated $F$. Bottom right of each solid line (tension isocline) corresponds to 100% D1A1 state and top left approaches the limit of 100% D1A0 state. Three dots provide references indicating 90%:10%, 50%:50%, and 10%:90% sensors in D1A1:D1A0 state, respectively. (d) ($E_{app}$, $S_{app}$)-curves for MTS ensembles comprised of various amounts of sensors in the D1A1 and D0A1 state (i.e. varying levels of donor mechanical switching only) with all sensors subject to the indicated $F$. Top of each solid line (tension isocline) corresponds to 100% D1A1 state and bottom approaches the limit of 100% D0A1 state. Three dots provide references indicating 90%:10%, 50%:50%, and 10%:90% sensors in D1A1:D0A1 state, respectively.

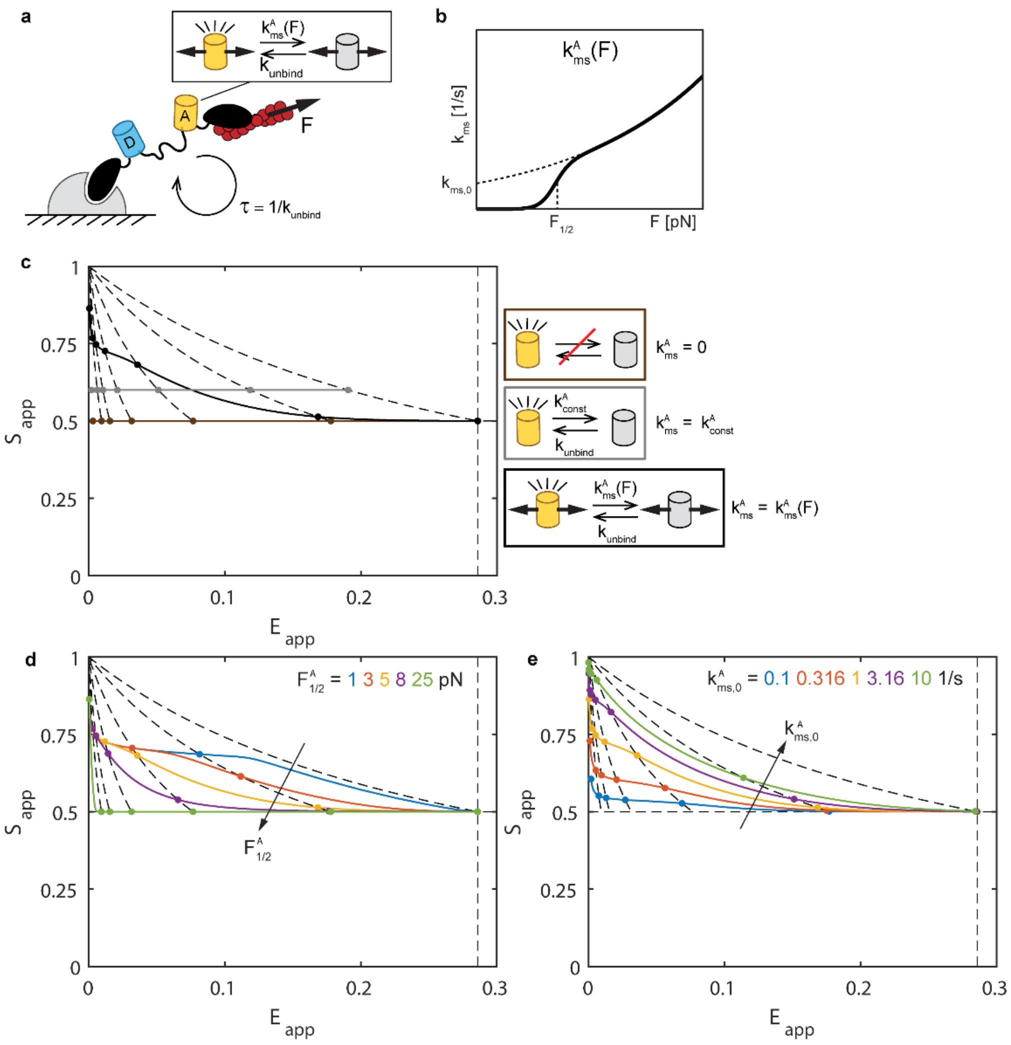


**Figure S5, related to Figure 1. Effect of Acceptor Mechanical Switching in MTS.** (a) Schematic of MTS with acceptor mechanical switching only. (b) Force-dependent rate constant for acceptor mechanical switching. (c) ($E_{app}$, $S_{app}$)-curves for MTS ensembles without acceptor mechanical switching [$k^A_{MS}(F) = 0$], force-independent acceptor loss-of-function [$k^A_{MS}(F) = k^A_{MS,0} = 1/s$ ; e.g. due to photobleaching], and acceptor mechanical switching [$k^A_{MS}(F)$ with base parameter values in Table S2] for a range of $F$ from 0 to 30 pN. (d) ($E_{app}$, $S_{app}$)-curves for different $F^A_{1/2}$ (and corresponding $m^A$ in Table S2). (e) ($E_{app}$, $S_{app}$)-curves for different $k^A_{MS,0}$. In all plots, reference dots indicate $F$ of 0, 3, 6, 9, 12, 15, and 30 pN (from right-to-left) and reference black dashed lines are tension isoclines for acceptor only mechanical switching at $F$ of 0, 3, 6, 9, 12, and 15 pN (from right-to-left). Except where indicated, all parameters are set to base values indicated in Table S2 and there is no donor mechanical switching [$k^D_{MS}(F) = 0$].

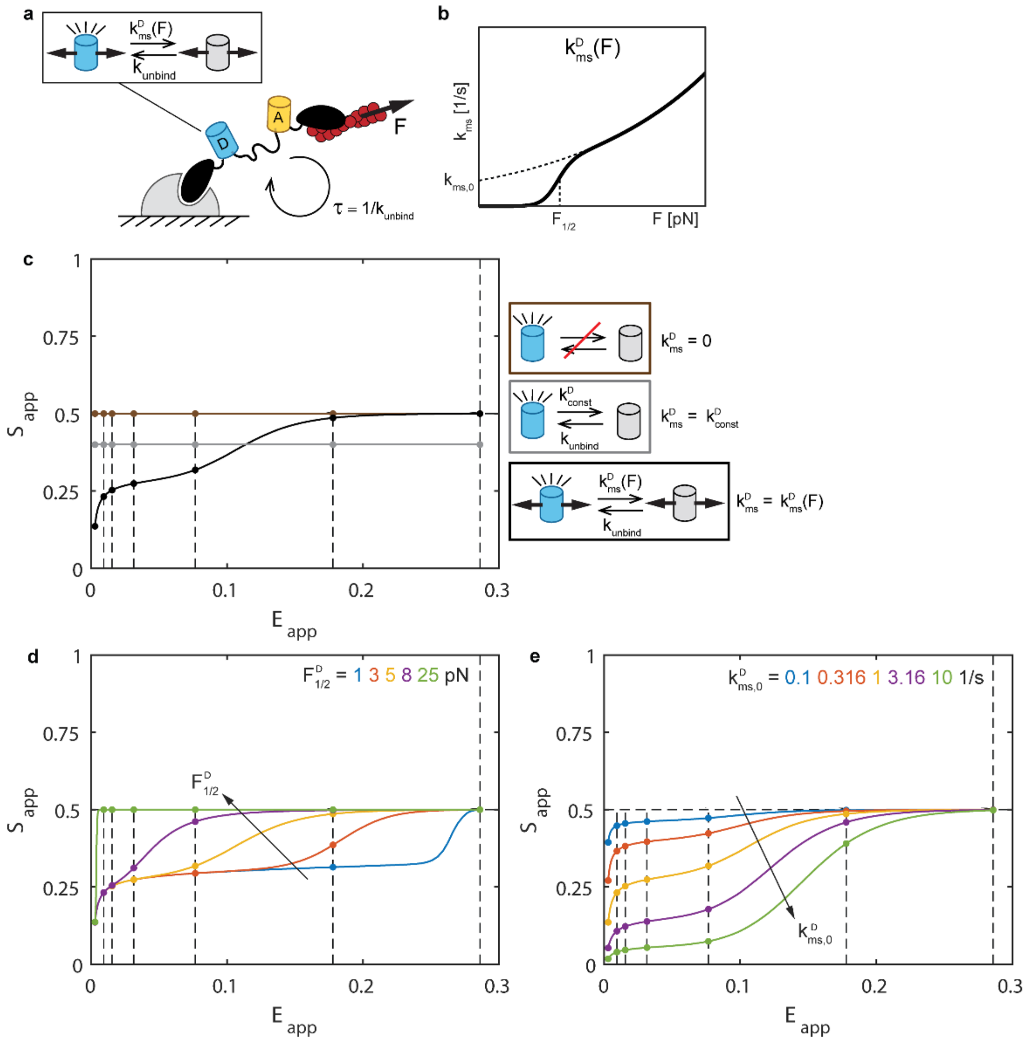


**Figure S6, related to Figure 1. Effect of Donor Mechanical Switching in MTS.** (a) Schematic of MTS with donor mechanical switching only. (b) Force-dependent rate constant for donor mechanical switching. (c) $(E_{app}, S_{app})$-curves for MTS ensembles without donor mechanical switching $[k^D_{MS}(F) = 0]$, force-independent donor loss-of-function $[k^D_{MS}(F) = k^D_{MS,0} = 1/s$ ; e.g. due to photobleaching], and donor mechanical switching $[k^D_{MS}(F)$ with base parameter values in Table S2] for a range of $F$ from 0 to 30 pN. (d) $(E_{app}, S_{app})$-curves for different $F^D_{1/2}$ (and corresponding $m^D$ in Table S2). (e) $(E_{app}, S_{app})$-curves for different $k^D_{MS,0}$. In all plots, reference dots indicate $F$ of 0, 3, 6, 9, 12, 15, and 30 pN (from right-to-left) and reference black dashed lines are tension isoclines for donor only mechanical switching at $F$ of 0, 3, 6, 9, 12, and 15 pN (from right-to-left). Except where indicated, all parameters are set to base values indicated in Table S2 and there is no acceptor mechanical switching $[k^A_{MS}(F) = 0]$.

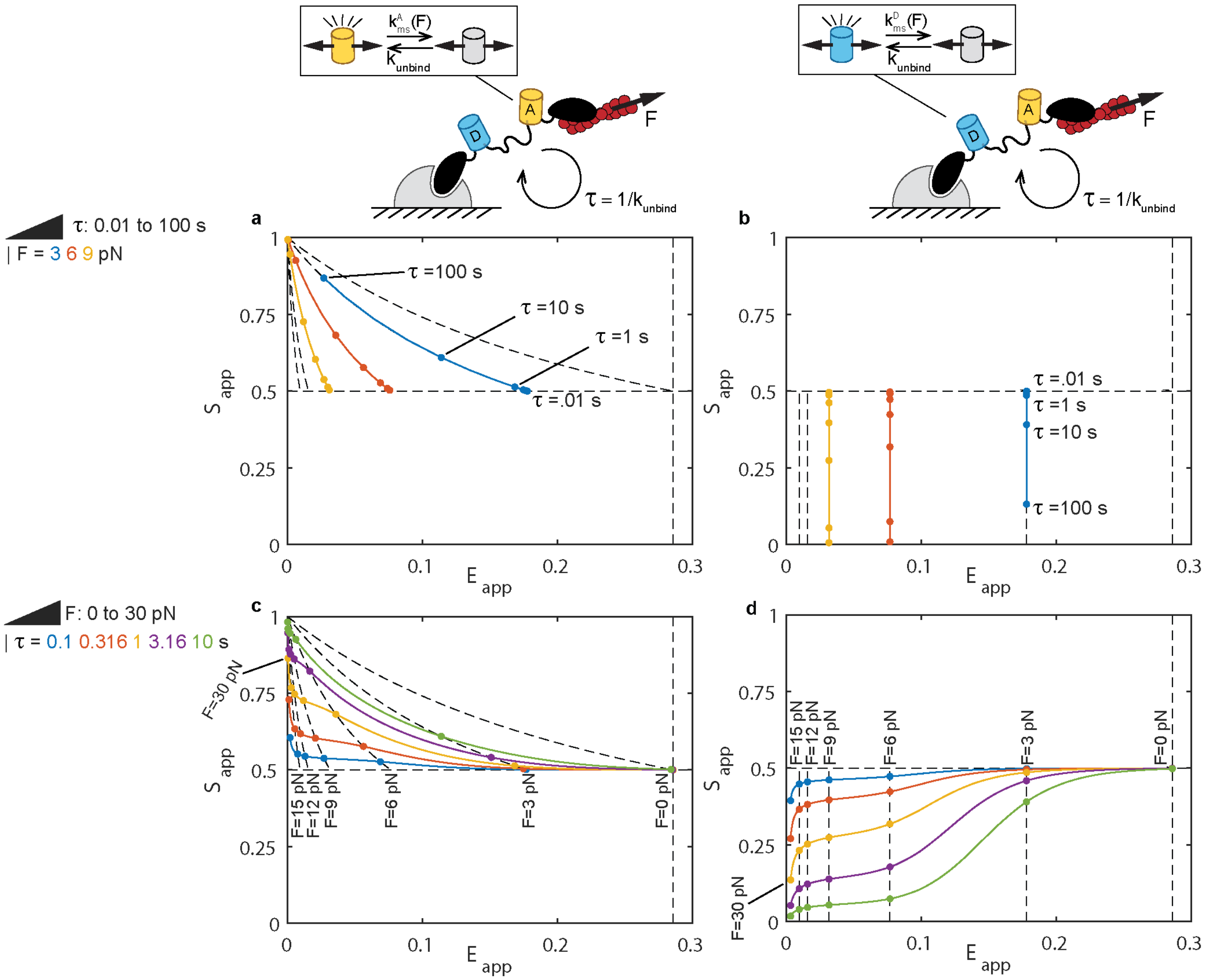


**Figure S7, related to Figure 1. Comparison on Variations in Load Magnitude versus Load Duration.** (a) For acceptor mechanical switching only [$k_{MS}^{A}(F)$ with base parameter values in Table S2 and $k_{MS}^{D}(F) = 0$], ($E_{app}$, $S_{app}$)-curves for variation of load duration $\tau$ from 0.01 to 100 seconds at three different $F$ values. Dots indicate select $\tau$ values for reference. (b) Same for donor mechanical switching only [$k_{MS}^{D}(F)$ with base parameter values in Table S2 and $k_{MS}^{A}(F) = 0$]. (c) For acceptor mechanical switching only [$k_{MS}^{A}(F)$ with base parameter values in Table S2 and $k_{MS}^{D}(F) = 0$], ($E_{app}$, $S_{app}$)-curves for variation of $F$ from 0 to 30 pN at five different $\tau$ values. Dots indicate select $F$ values for reference. (d) Same for donor mechanical switching only [$k_{MS}^{D}(F)$ with base parameter values in Table S2 and $k_{MS}^{A}(F) = 0$]. In all plots, black dashed lines are tension isoclines for acceptor only or donor only mechanical switching at $F$ of 0, 3, 6, 9, 12, and 15 pN. Except where indicated, all parameters are set to base values indicated in Table S2.

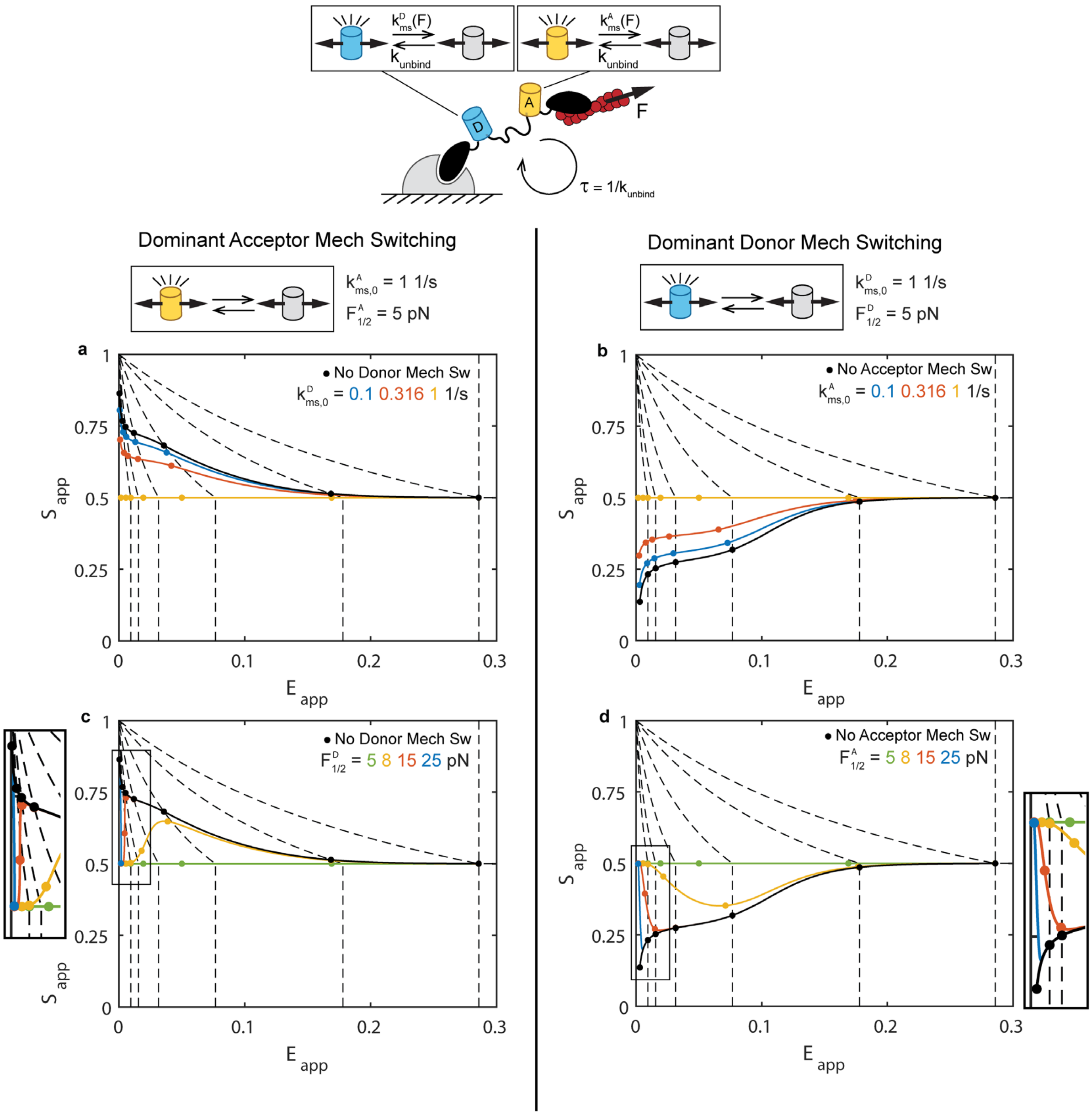


**Figure S8, related to Figure 1. Dominant Mechanical Switching in One FP is Detectable In Presence of Weaker Mechanical Switching in the Other FP.** (a) For base acceptor mechanical switching [$k^A_{MS}(F)$ with base parameter values in Table S2], ($E_{app}$, $S_{app}$)-curves for donor mechanical switching at different values of $k^D_{MS,0}$. (b) For base donor mechanical switching [$k^D_{MS}(F)$ with base parameter values in Table S2], ($E_{app}$, $S_{app}$)-curves for acceptor mechanical switching at different values of $k^A_{MS,0}$. (c) For base acceptor mechanical switching [$k^A_{MS}(F)$ with base parameter values in Table S2], ($E_{app}$, $S_{app}$)-curves for donor mechanical switching at different values of $F^D_{1/2}$ (and corresponding $m^D$ in Table S2). (d) For base donor mechanical switching [$k^D_{MS}(F)$ with base parameter values in Table S2], ($E_{app}$, $S_{app}$)-curves for acceptor mechanical switching at different values of $F^A_{1/2}$ (and corresponding $m^A$ in Table S2). In all plots, reference dots indicate $F$ of 0, 3, 6, 9, 12, 15, and 30 pN (from right-to-left) and reference black dashed lines are tension isoclines for $F$ of 0, 3, 6, 9, 12, and 15 pN (from right-to-left). Except where indicated, all parameters are set to base values indicated in Table S2.

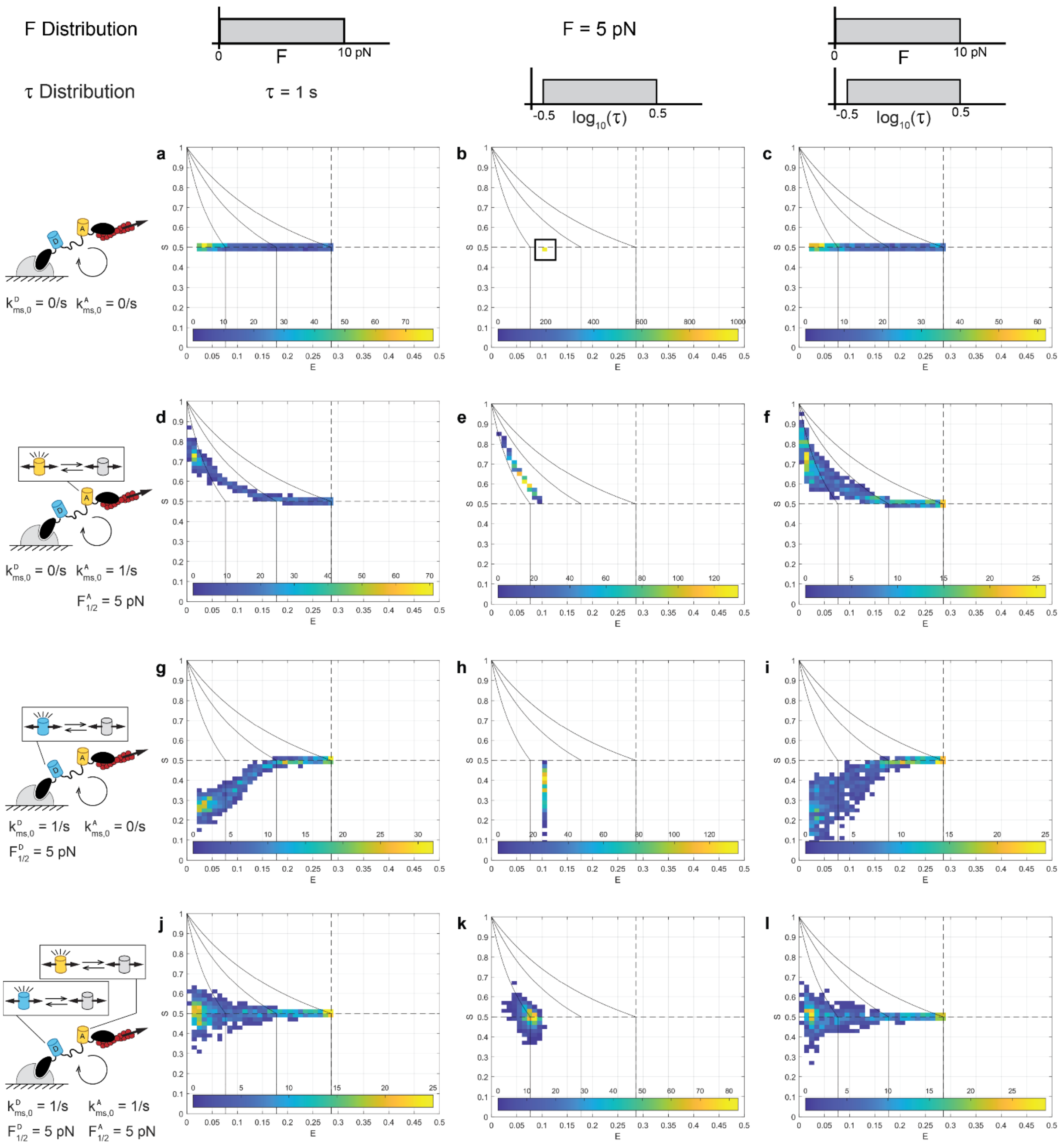


**Figure S9, related to Figure 1. Representative ES-Histograms for Stochastic Simulations.** For no FP mechanical switching [$k^A_{MS}(F) = k^D_{MS}(F) = 0$], ES-Histogram of 1000 simulated MTS ensembles ($N_{sims} = 1000$), each comprised of 50 total sensors ($n_{sensors} = 50$) subject to (a) a single $F$ value drawn from a uniform distribution between 0 and 10 pN and a single $\tau$ value of 1 s, (b) a single $F$ value of 5 pN and a single $\tau$ value drawn from a log-uniform distribution from $10^{-0.5}$ to $10^{0.5}$ s, or (c) a single $F$ value drawn from a uniform distribution between 0 and 10 pN and a single $\tau$ value drawn from a log-uniform distribution from $10^{-0.5}$ to $10^{0.5}$ s. (d-f) Same for acceptor only mechanical switching [$k^A_{MS}(F)$ with base parameter values in Table S2 and $k^D_{MS}(F) = 0$]. (g-i) Same for donor only mechanical switching [$k^D_{MS}(F)$ with base parameter values in Table S2 and $k^A_{MS}(F) = 0$]. (j-l) Same for acceptor and donor mechanical switching with identical parameters [$k^A_{MS}(F)$ and $k^D_{MS}(F)$ with base parameter values in Table S2]. In all plots, reference black lines are tension isoclines for acceptor only or donor only mechanical switching at $F$ of 0, 3, and 6 pN (from right-to-left). Note that within single MTS ensembles, all sensors are subject to the same $F$ and $\tau$ values.

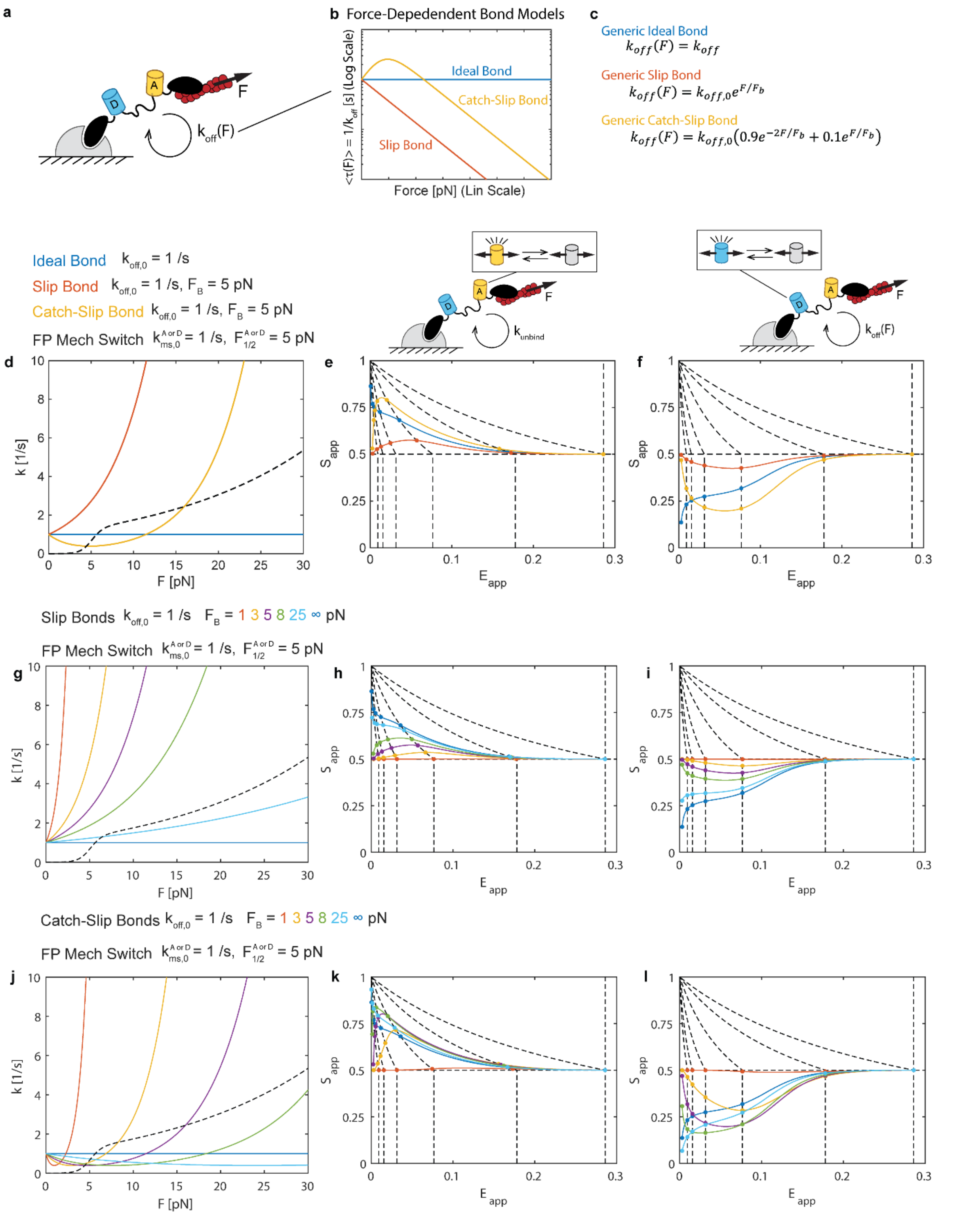


**Figure S10, related to Figure 1. Effect of Force-Dependent Bonds on FP Mechanical Switching in MTS.** (a) Schematic of MTS with turnover driven by force-dependent unbinding. (b) Mean lifetime versus force and (c) force-dependent unbinding rate constant expressions for generalized ideal, slip, and catch-slip bonds. (d) Force-dependent unbinding rate constants for three bond types at indicated parameters (matched intrinsic rate constant and exponential parameters), shown with the FP mechanical switching rate constant for comparison. (e) For acceptor mechanical switching only [$k_{MS}^{A}(F)$ with base parameter values in Table S2 and $k_{MS}^{D}(F) = 0$], ($E_{app}$, $S_{app}$)-curves for the different unbinding rate constants in (d) for a range of $F$ from 0 to 30 pN. (f) Same for donor mechanical switching only [$k_{MS}^{D}(F)$ with base parameter values in Table S2 and $k_{MS}^{A}(F) = 0$]. (g) Force-dependent unbinding rate constants for slip bonds with different $F_B$ values and the FP mechanical switching rate constant shown for comparison. For (h) acceptor or (i) donor only mechanical switching, ($E_{app}$, $S_{app}$)-curves for the different unbinding rate constants in (g) for a range of $F$ from 0 to 30 pN. (j) Force-dependent unbinding rate constants for catch-slip bonds with different $F_B$ values and the FP mechanical switching rate constant shown for comparison. For (k) acceptor or (l) donor only mechanical switching, ($E_{app}$, $S_{app}$)-curves for the different unbinding rate constants in (j) for a range of $F$ from 0 to 30 pN. In (e-f,h-i,k-l), reference dots indicate $F$ of 0, 3, 6, 9, 12, 15, and 30 pN (from right-to-left) and reference black dashed lines are tension isoclines for acceptor or donor only mechanical switching at $F$ of 0, 3, 6, 9, 12, and 15 pN (from right-to-left). Except where indicated, all parameters are set to base values indicated in Table S2.

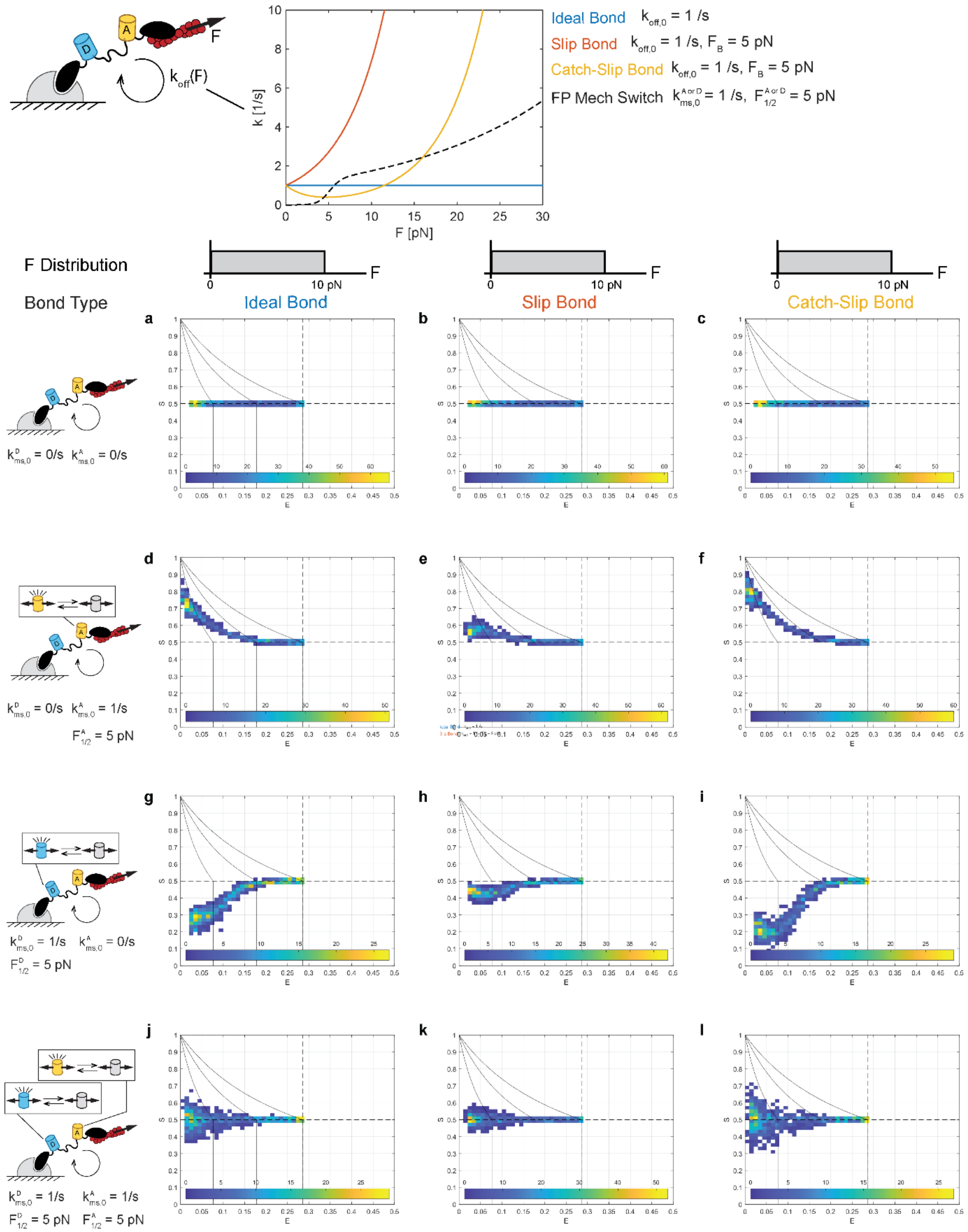


**Figure S11, related to Figure 1. ES-Histograms for Stochastic Simulations with Force-dependent Bonds.** (a-c) For no FP mechanical switching [$k^A_{MS}(F) = k^D_{MS}(F) = 0$], ES-Histogram of 1000 simulated MTS ensembles ($N_{sims} = 1000$), each comprised of 50 total sensors ($n_{sensors} = 50$) subject to single $F$ value drawn from a uniform distribution between 0 and 10 pN and having (a) ideal, (b) slip, or (c) catch-slip unbinding rate constant. (d-f) Same for acceptor only mechanical switching [$k^A_{MS}(F)$ with base parameter values in Table S2 and $k_{MS}^{D}(F) = 0$]. (g-i) Same for acceptor only mechanical switching [$k_{MS}^{D}(F)$ with base parameter values in Table S2 and $k_{MS}^{A}(F) = 0$]. (j-l) Same for acceptor and donor mechanical switching with identical parameters [$k_{MS}^{A}(F)$ and $k_{MS}^{D}(F)$ with base parameter values in Table S2]. In all plots, reference black lines are tension isoclines for acceptor only or donor only mechanical switching at $F$ of 0, 3, and 6 pN (from right-to-left). Note that within single MTS ensembles, all sensors are subject to the same $F$ value.

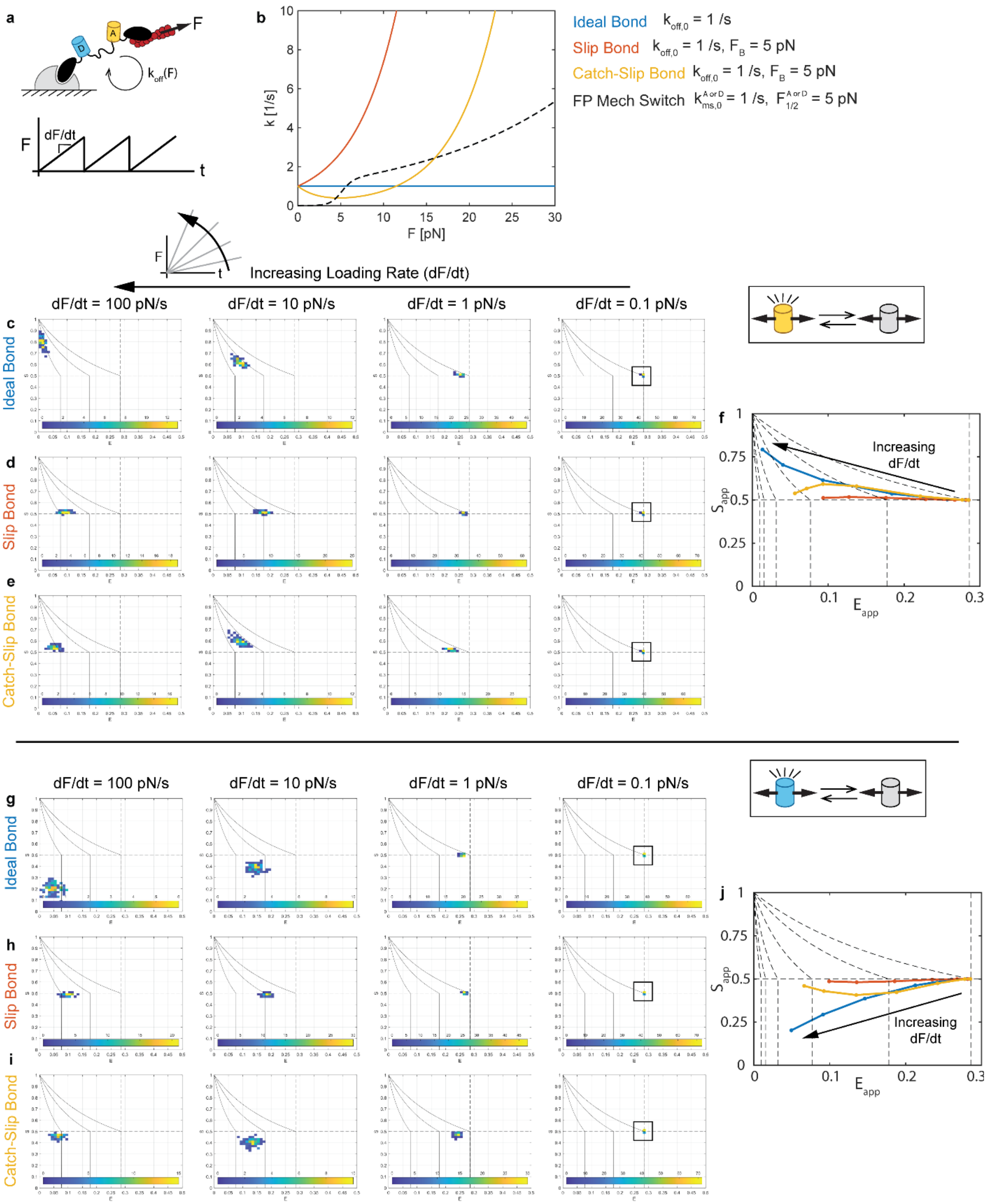


**Figure S12, related to Figure 1. Effect of Loading Rate on FP Mechanical Switching in MTS.** (a) Schematic of MTS subject to linear ramp (constant rate) loading. (b) Force-dependent unbinding rate constants for ideal, slip, and catch-slip bond types at indicated parameters (matched intrinsic rate constant and exponential parameters) with the FP mechanical switching rate constant shown for comparison. (c-e) For acceptor only mechanical switching [$k^{A}_{MS}(F)$ with base parameter values in Table S2 and $k^{D}_{MS}(F) = 0$] and MTS with an (c) ideal, (d) slip, or (e) catch-slip bond, ES-Histogram of 100 simulated MTS ensembles ($N_{sims} = 100$), each comprised of 50 total sensors ($n_{sensors} = 50$) subject to loading rates 100, 10, 1, or 0.1 pN/s. (f) For acceptor mechanical switching and the three bond types, ($E_{app}$, $S_{app}$)-curves for a range of loading rate $dF/dt$ from 0.1 to 100 pN/s constructed from the mean values from the stochastic simulations at each loading rate. Reference dots indicate loading rates of 0, 0.32, 1, 3.2, 10, 32, and 100 pN/s (from right-to-left) and reference black dashed lines are tension isoclines for acceptor or donor only mechanical switching at $F$ of 0, 3, 6, 9, 12, and 15 pN (from right-to-left). (g-i) Same as (c-e) but for donor only mechanical switching [$k_{MS}^{D}(F)$ with base parameter values in Table S2 and $k_{MS}^{A}(F) = 0$] and MTS with (g) ideal, (h) slip, or (i) catch-slip bond. (j) Same as (f) but for donor only mechanical switching.

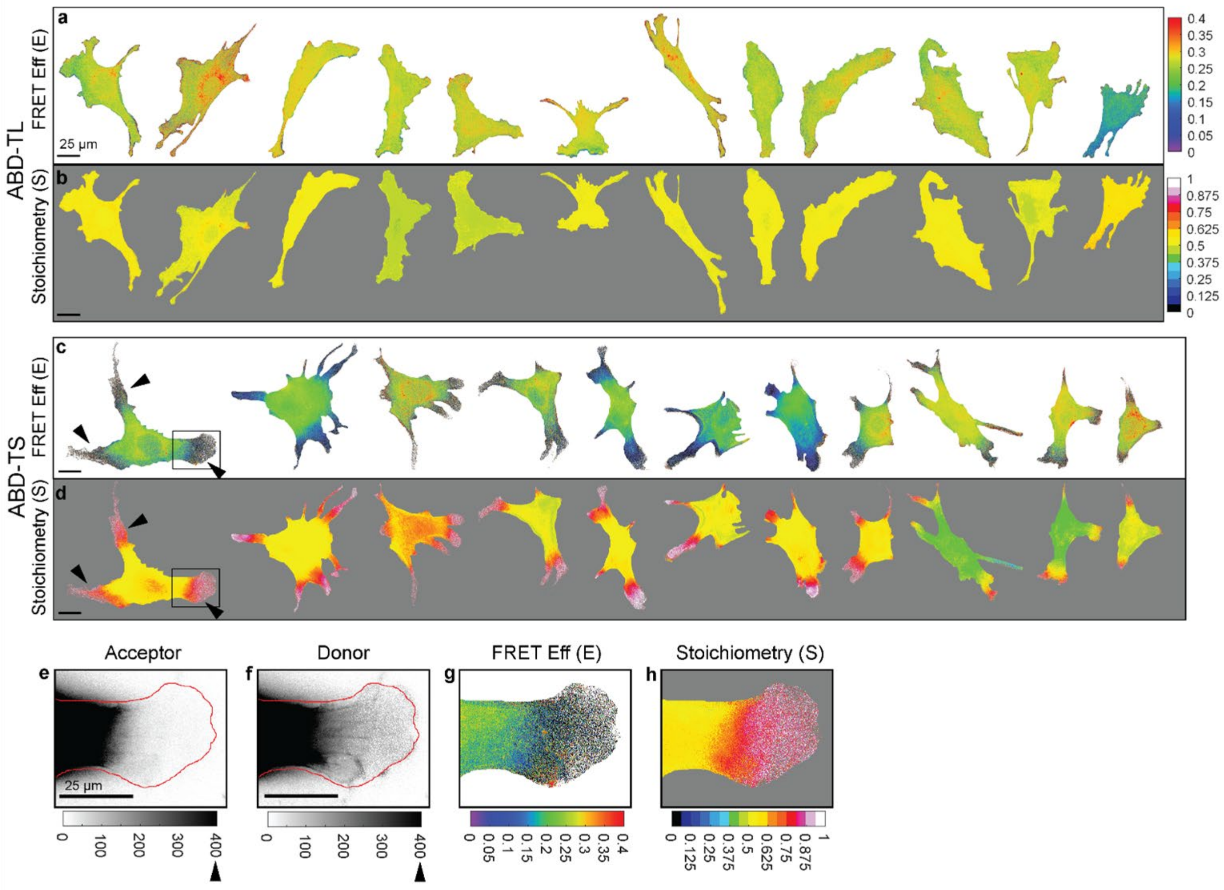


**Figure S13, related to Figure 2. Signatures of FP mechanical switching in ABDTS are consistent and exhibit a distinct spatial pattern**. Montage of >10 NIH 3T3 cells expressing (a-b) ABDTL or (c-d) ABDTS from a single experimental day, showing FRET efficiency and stoichiometry. (e-h) Zoom of indicated region at periphery of a cell expressing ABDTS, showing acceptor and donor intensities with cell outline overlaid in red and FRET efficiency and Stoichiometry in cell mask.

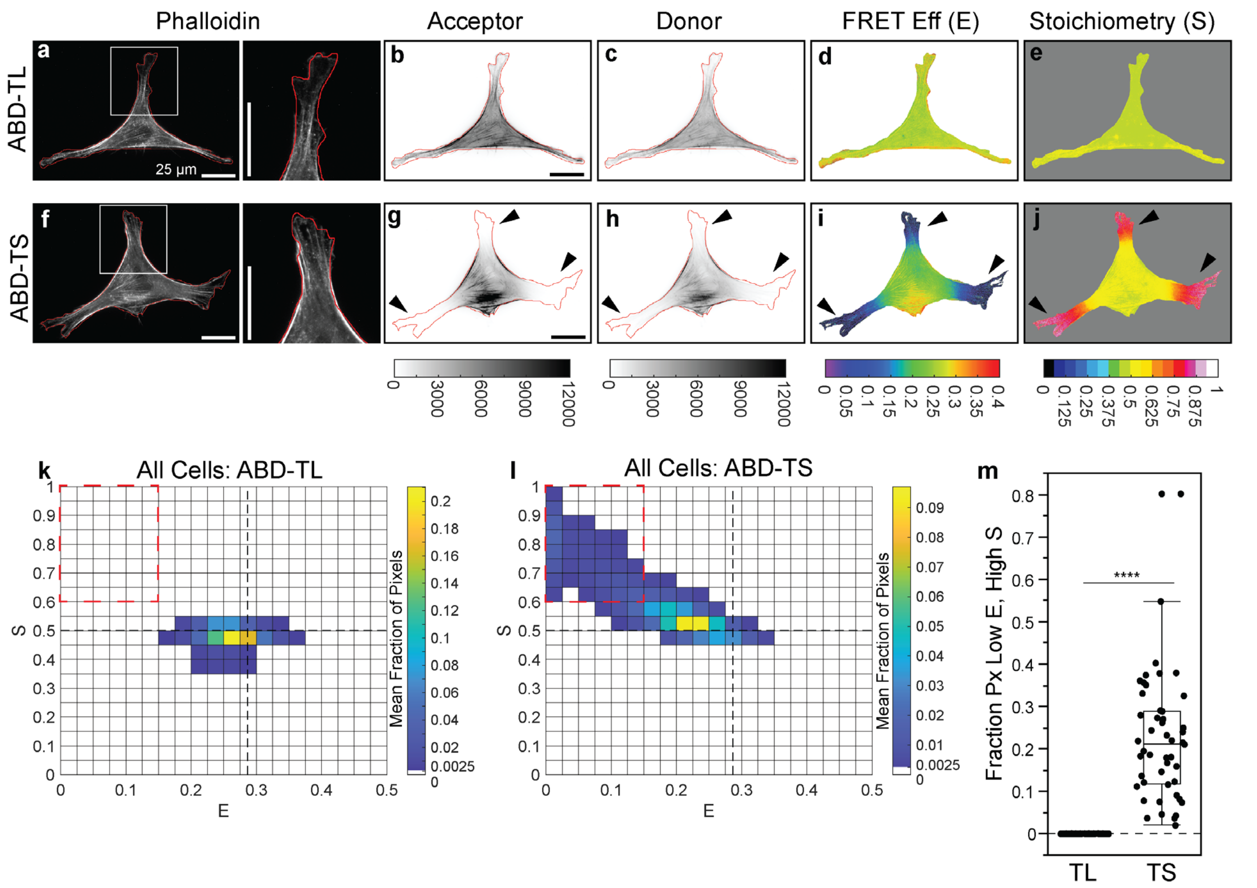


**Figure S14, related to Figure 2. Fixation and phalloidin labeling of F-actin in ABDTL- and ABDTS-expressing cells.** Representative NIH 3T3 cells expressing ABDTL (a-e) or ABDTS (f-j) fixed and labeled with phalloidin. Images are phalloidin intensity used to create cell outline, acceptor and donor intensities with cell outline overlaid in red, and FRET efficiency and Stoichiometry in cell mask. ES-histograms for whole cell populations of ABDTL (k) or ABDTS (l), indicating the cell-averaged fraction of cell-masked pixels in each bin (N = 64/48 cells for ABDTL/ABDTS over 2 experimental days). (m) Box plot of fraction of pixels in each cell in the low $E$, high $S$ bin ($E < 0.15,\ S > 0.60$). Differences between groups were detected using the Steel-Dwass test, ****: p<0.0001.

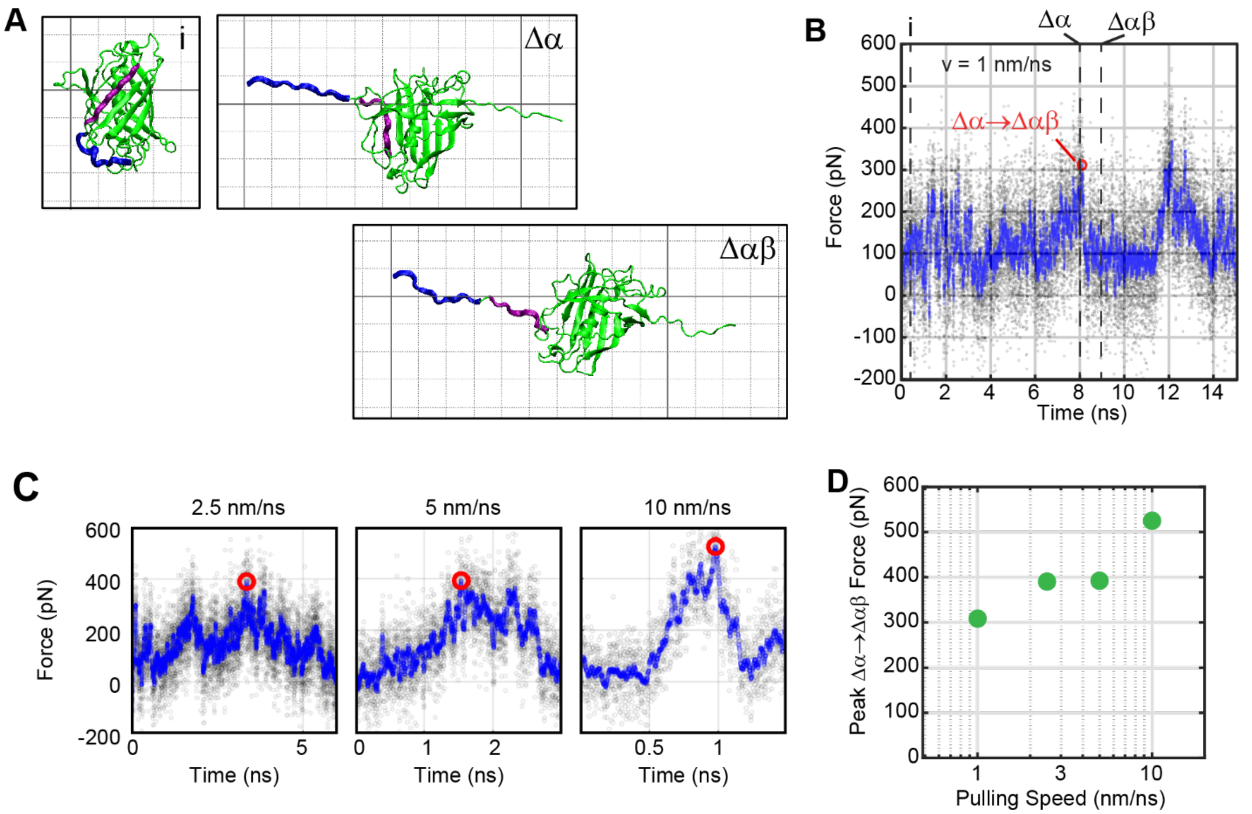


**Figure S15. Steered molecular dynamics simulations of alpha-GFP.** (A) Three snapshots of a representative simulation of GFP unfolding (at a pulling speed of 1 nm/ns) are shown: The initial state just before the application of force (i), the first intermediate state wherein GFP's N-terminal handle (blue) has been removed from the structure (Δα), and what we believe to be the mechanically switched intermediate wherein a β-strand (purple) has been removed from GFP's β-barrel structure (Δαβ). (B) A force vs. time plot from the simulation in (A) showing the force-extension curve (black translucent circles) and the force-extension curve smoothed by 250 points (blue circles). Vertical dashed lines show the timepoints of the snapshots in (A). A red circle shows the maximum force, which coincides with the transition from the Δα state to the Δαβ state. (C) Force-extension curves for additional loading rates, with the peak force before the Δα to Δαβ transition shown with red circles. (D) The peak force before the Δα to Δαβ transition, shown as a function of the pulling speed. Each point represents the maximum from a single simulation.

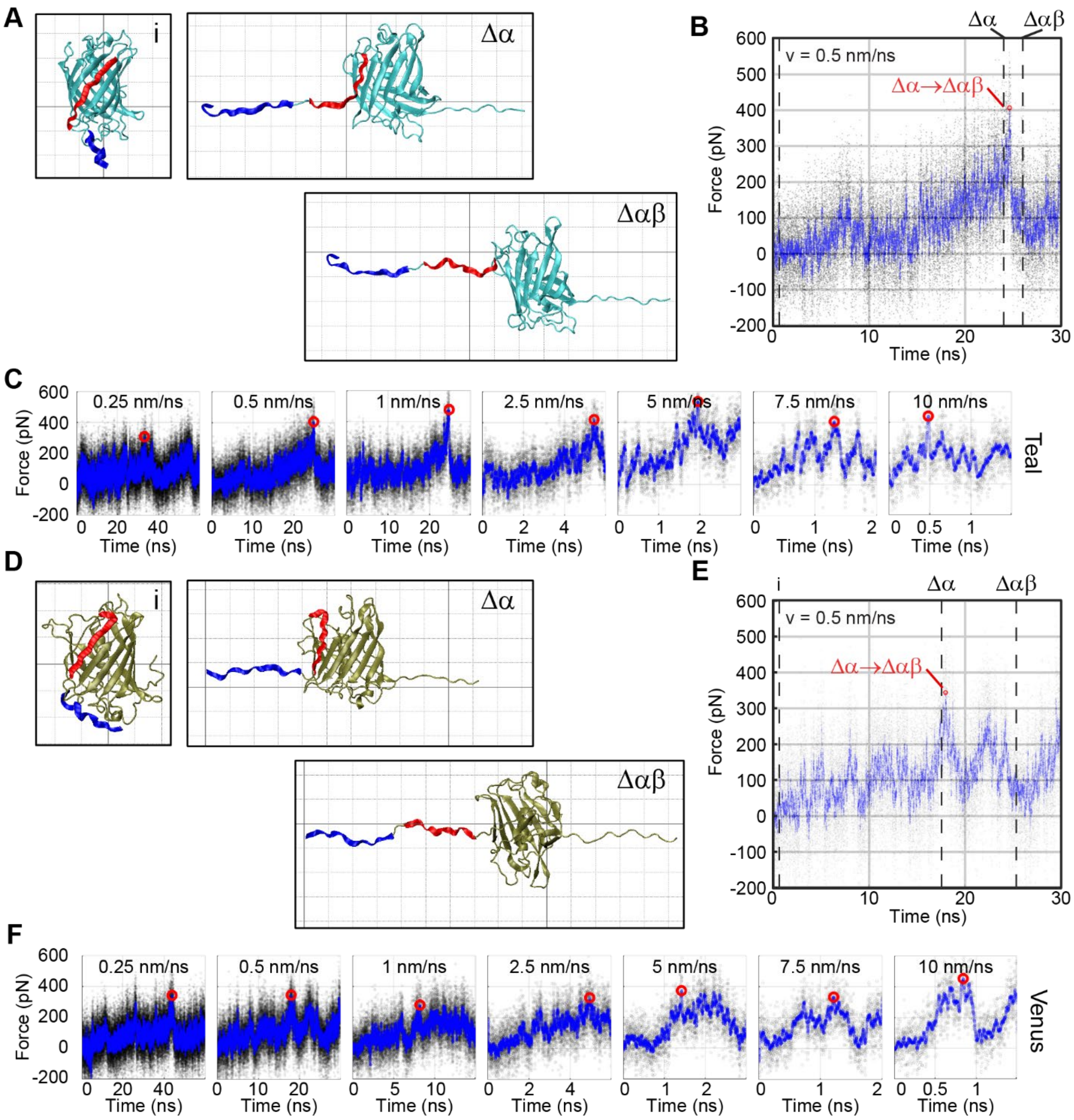


**Figure S16. Steered molecular dynamics simulations of mTFP1 and mVenus**. (A-C) Like Figure S15A-C, but for simulations of mTFP1. (D-F) Like Figure S15A-C, but for simulations of mVenus.

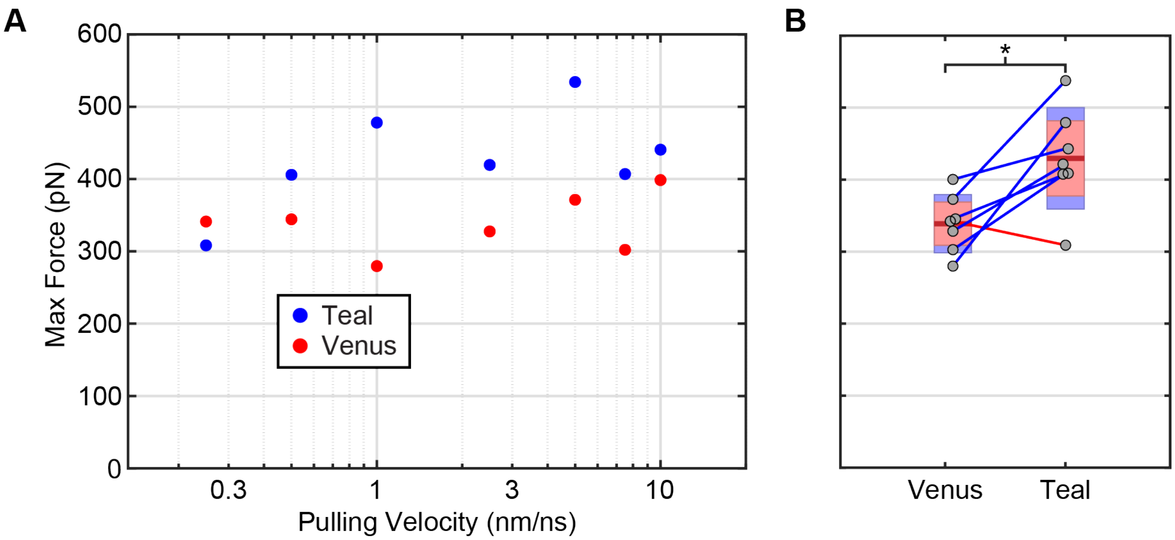


**Figure S17. Comparison of peak forces from SMD simulations for mVenus and mTFP1.** (A) The peak force before the Δα to Δαβ transition, shown as a function of the pulling speed for simulations of mTFP1 (blue) and mVenus (red) from Figure S16. B) Boxplots showing the maximum forces from (A) pooled for a paired comparison between mVenus and mTFP1. Boxplots are shown with average (maroon line), standard error of the mean (pink), and 95% confidence intervals (purple), as well as all individual datapoints (gray circles). Pairings between datapoints are shown via lines, where blue line color denotes that the maximum force for mTFP1 is higher than that of mVenus and red line color denotes the reverse. A statistical comparison between the two groups (paired T-test) shows that mTFP1 has a significantly higher rupture force than mVenus ($p$ = 0.047). These findings are consistent with our experimental observations.

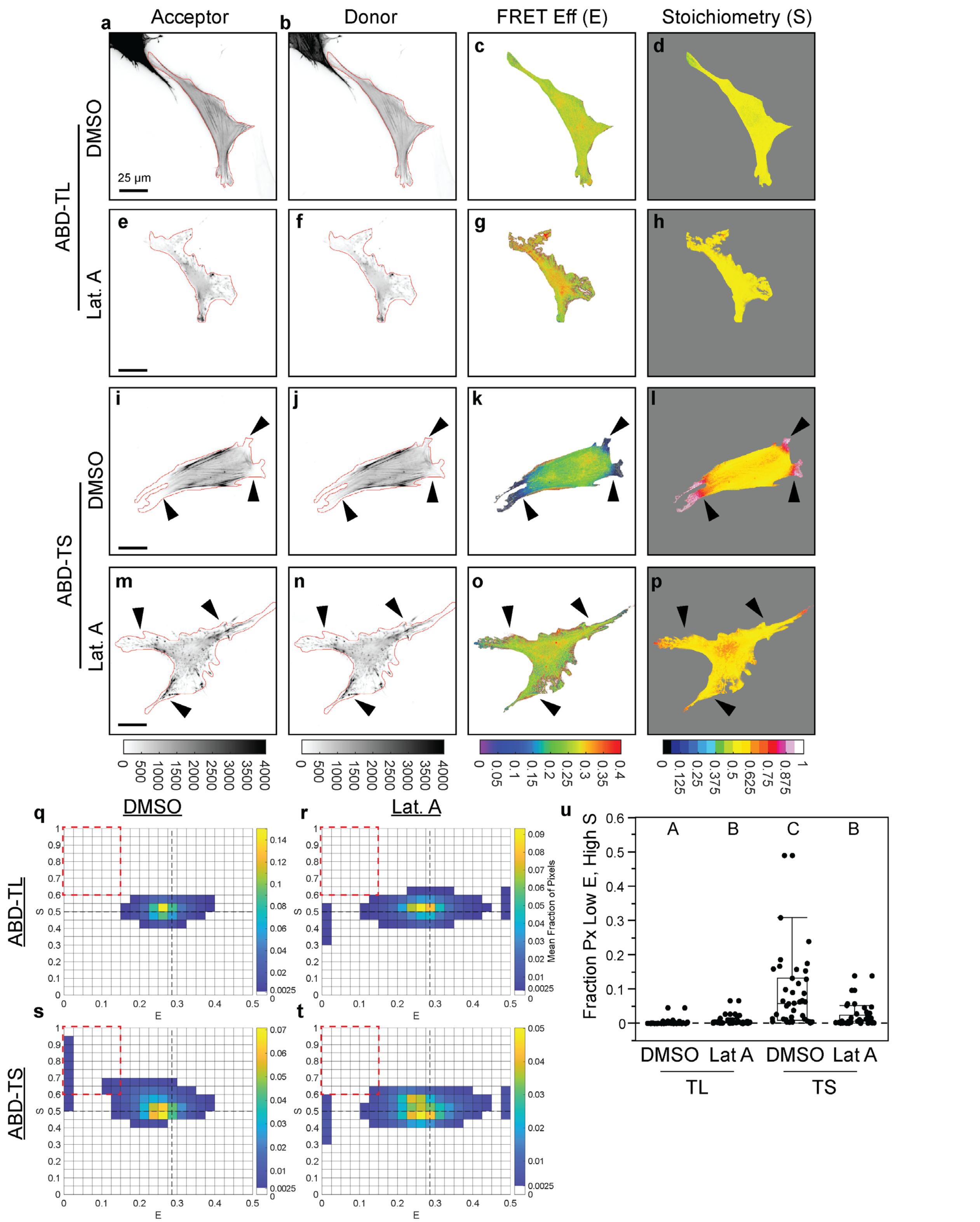


**Figure S18, related to Figure 2. Pharmacological disruption of actin reverses FP mechanical switching in ABDTS.** Representative NIH 3T3 cells expressing ABDTL with vehicle control (a-d), ABDTL treated with Latrunculin A (e-h), ABDTS with vehicle control (i-l), or ADBTS treated with Latrunculin A (m-p). Images are acceptor and donor intensities with cell outline overlaid in red and FRET efficiency and

Stoichiometry. (q-t) ES-histograms for whole cell populations of ABDTL with vehicle control (q, N=43 cells), ABDTL treated with Latrunculin A (r, N=40 cells), ABDTS with vehicle control (s, N=39 cells), and ADBTS treated with Latrunculin A (t, N=40 cells) over 3 experimental days. (u) Box plot of fraction of pixels in each cell in the low $E$, high $S$ bin ($E < 0.15,\ S > 0.60$) for the indicated conditions. Differences between groups were detected using the Steel-Dwass test. Levels not connected by the same letter are significantly different at p<0.05.

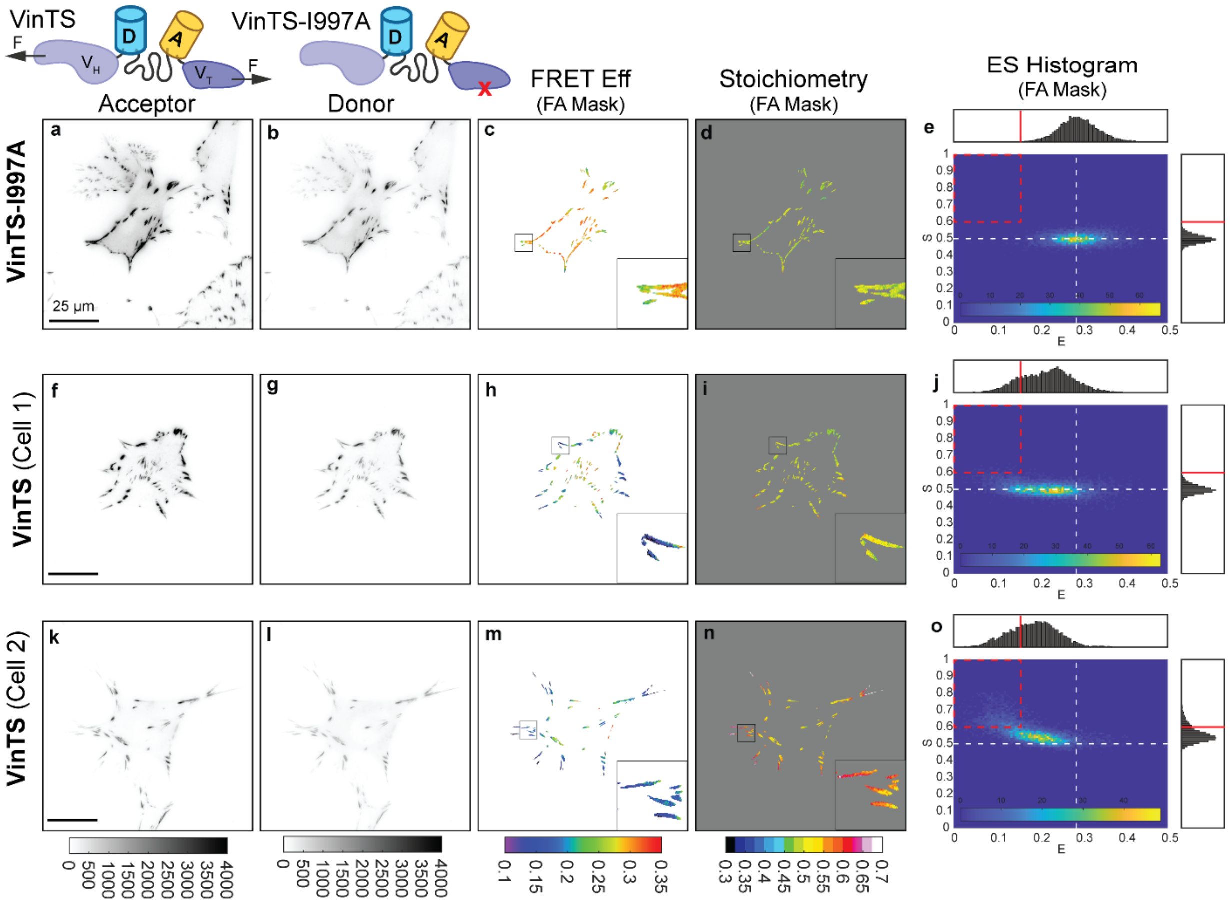


**Figure S19, related to Figure 3. Signatures of weaker, but detectable, FP mechanical switching in VinTS.** Representative vinculin -/- MEFs expressing VinTS-I997A (a-e) or VinTS (f-o) on FN-coated glass, showing images of acceptor intensity, donor intensity, FRET efficiency in the FA mask, Stoichiometry in the FA mask, and an ES-histogram of FA-masked pixels for the cell. For VinTS, the two representative cells correspond to the two ends of the spectrum of behaviors observed. Cell 1 (f-j) matches model predications for MTS loading without FP mechanical switching. Cell 2 (k-o) matches model predications for MTS loading with acceptor mechanical switching. The data is a re-analysis of three-channel FRET images from an experiment in a previous publication[1].

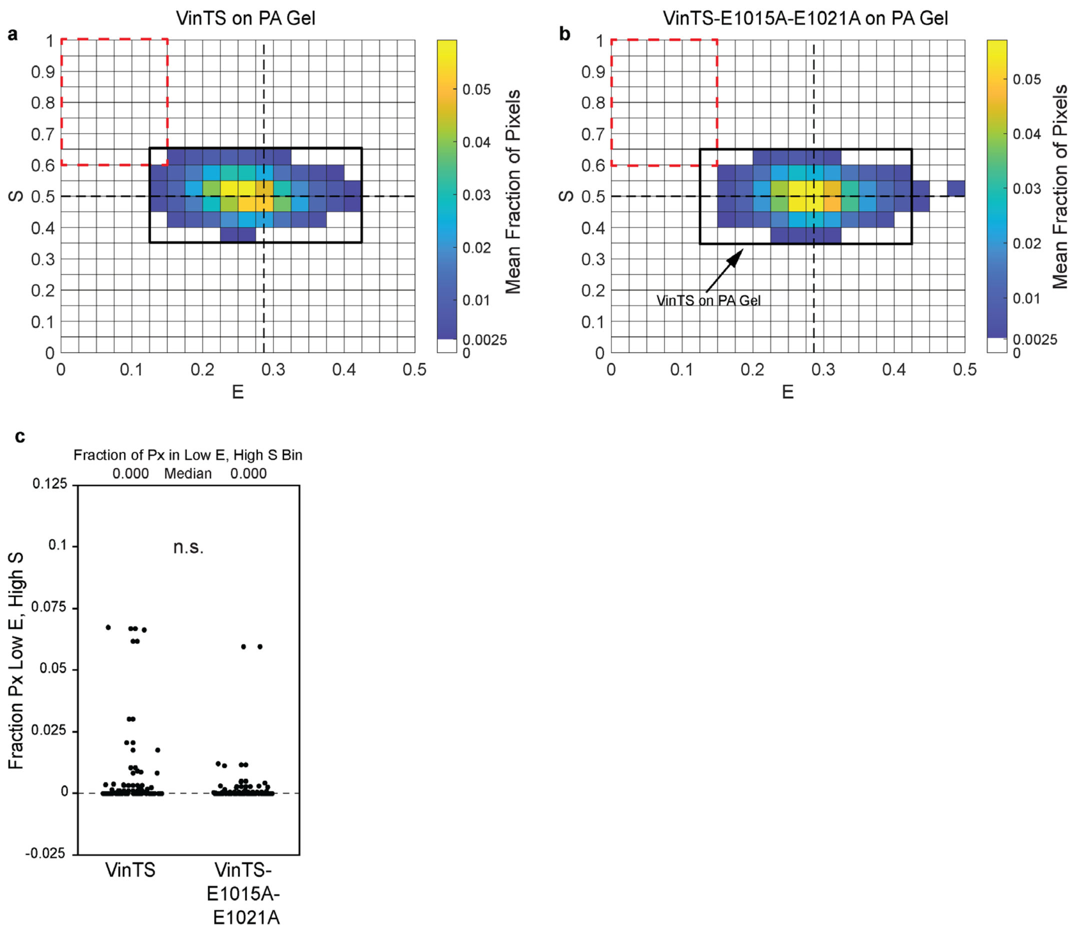


**Figure S20, related to Figure 3. Acceptor mechanical switching in VinTS is eliminated on soft substrates**. (a-b) ES-histograms for whole cell populations of vinculin -/- MEFs expressing VinTS or VinTS-E1015A-E1021A on PA Gels, indicating the cell-averaged fraction of FA-masked pixels in each bin (N = 87/92 cells over 3 experimental days for VinTS/VinTS-E1015A-E1021A). The black outline for VinTS from (a) is overlaid on the histogram for VinTS-E1015A-E1021A in (b) as a guide for the eye. (c) Box plot of fraction of FA-masked pixels in each cell in the low $E$, high $S$ bin ($E < 0.15,\ S > 0.60$).

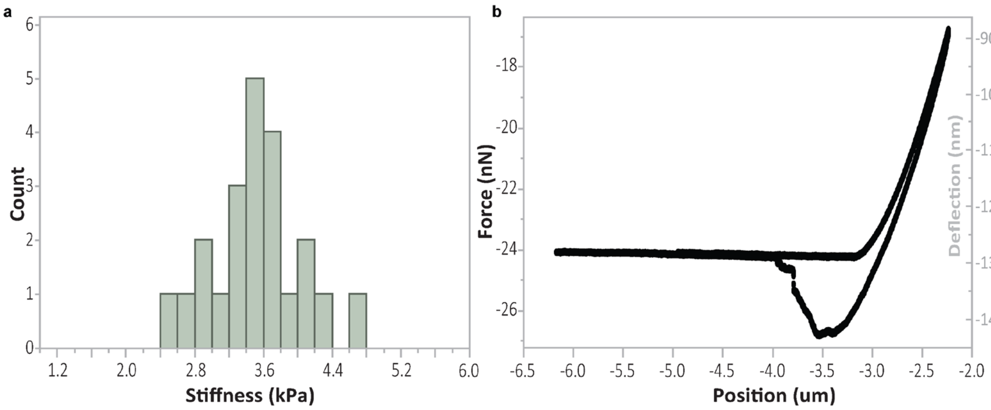


**Figure S21, related to Figure 3. Mechanical testing of the PA gel.** (a) Histogram of Young's modulus estimates for the PA gel formulation used in the experiment in Figure S20. Young's modulus was determined via Hertzian model fit from mechanical tests by atomic force microscopy (AFM). (b) Representative trace from mechanical testing. See methods for more information on mechanical testing.

### Supplemental Movie Legends

**Movie S1.** Timelapse showing unfolding of alpha-GFP at a pulling speed of 1 nm/ns (left) and a force vs. time plot from the same simulation (right). Grid ticks in the timelapse correspond to 0.1 nm. The N-terminal handle of the FP is colored blue, while the first beta-strand is colored purple and the remainder of the FP is colored green. The simulation timepoint is indicated via a vertical dashed line in the force vs. time plot. Black translucent circles in the force vs. time plot show the raw data, while the blue circles show the force vs. time plot smoothed by 250 points. A red circle shows the maximum force value before the pullout of the beta strand, which coincides with the transition from the Δα state to the Δαβ state.

**Movie S2.** Timelapse showing unfolding of mTFP1 at a pulling speed of 1 nm/ns (left) and a force vs. time plot from the same simulation (right). Grid ticks in the timelapse correspond to 0.1 nm. The N-terminal handle of the FP is colored blue, while the first beta-strand is colored red and the remainder of the FP is colored teal. The simulation timepoint is indicated via a vertical dashed line in the force vs. time plot. Black translucent circles in the force vs. time plot show the raw data, while the blue circles show the force vs. time plot smoothed by 250 points. A red circle shows the maximum force value before the pullout of the beta strand, which coincides with the transition from the Δα state to the Δαβ state.

**Movie S3.** Timelapse showing unfolding of mVenus at a pulling speed of 1 nm/ns (left) and a force vs. time plot from the same simulation (right). Grid ticks in the timelapse correspond to 0.1 nm. The N-terminal handle of the FP is colored blue, while the first beta-strand is colored red and the remainder of the FP is colored yellow. The simulation timepoint is indicated via a vertical dashed line in the force vs. time plot. Black translucent circles in the force vs. time plot show the raw data, while the blue circles show the force vs. time plot smoothed by 250 points. A red circle shows the maximum force value before the pullout of the beta strand, which coincides with the transition from the Δα state to the Δαβ state.

### Note S1: Mathematical Models of FP Mechanical Switching in Load-bearing Proteins and FRET-based Molecular Tension Sensors

### I. Introduction

This supplemental note covers the formulation, results, and analysis of mathematical models of fluorescent protein (FP) mechanical switching in load-bearing proteins and Förster resonance energy transfer (FRET)-based molecular tension sensors (MTS). To investigate the process of FP mechanical switching in the context of protein loading in cells, we first modeled the reversible mechanical switching of a single FP integrated into a load-bearing protein and assessed the extent of FP mechanical switching across load magnitudes and durations estimated for protein loading in cells (Section II). Then, to assess how FP mechanical switching affects FRET-based MTSs, we extended the model of FP mechanical switching to an MTS containing a donor and an acceptor FP (Section III). Using simulated three channel FRET measurements analogous to experimentally accessible readouts, we developed and applied a framework for detecting FP mechanical switching in MTSs. To facilitate data visualization, we conducted simulations that account for phenomena expected in real experimental data, including variability in protein loading dynamics and intrinsic noise in FP mechanical switching (Section IV). Lastly, the effect of two additional aspects of protein loading dynamics, force-sensitive bonds and loading rate control, on FP mechanical switching in MTSs was assessed (Section V). Assumptions and limitations of the models are discussed in Section VI, and the main conclusions are summarized in Section VII.

### II. Model of FP Mechanical Switching in a Load-bearing Protein

In this section, we investigate the process of FP mechanical switching in the context of load-bearing proteins inside cells. To do so, we develop and use a model describing the reversible mechanical switching of an FP integrated into a load-bearing protein subject to protein loading dynamics.

#### A. Model Formulation

##### 1. Kinetic Model

Mechanical forces applied externally to cells or generated internally by cells are transmitted through proteins that form mechanical linkages within the cytoskeleton or between the cytoskeleton and other subcellular structures, including adhesions and the plasma membrane[2-5]. Proteins in these linkages bear dynamic loads, exhibit binding/unbinding dynamics, and turnover[2-5]. Therefore, we modeled an FP integrated into a load-bearing protein subject to dynamic loading parameterized by a load magnitude $F$ and a characteristic load duration $\tau$, which is governed by unbinding from a loading source with rate constant $k_{unbind}$ (where $\tau \equiv 1/k_{unbind}$) (Figure S1a).

Our model of FP mechanical switching is motivated by single molecule experiments demonstrating that GFP fluorescence can be reversibly switched on/off by repeated cycles of mechanical tension[6]. Inside the load-bearing protein, the FP is modeled as a two-state system, with fluorescence on (FP1) or off (FP0). The FP experiences the force $F$ across the load-bearing protein and can undergo a mechanical switching transition (FP1 → FP0) according to a force-dependent mechanical switching rate constant, $k_{MS}(F)$. When the load-bearing protein unbinds from the loading source with rate constant $k_{unbind}$, an FP in the off state immediately recovers (FP0 → FP1) before the load-bearing protein rebinds and is reloaded. This is consistent with the rapid recovery of unloaded FPs and/or the exchange of load-bearing proteins with a cytosolic pool upon unbinding[6,7].

As we intended to interpret experiments with different types of FPs, we modeled a generalized force-dependent mechanical switching rate constant, $k_{MS}(F)$, that was informed from single molecule experimental observations of the response of GFP structure and fluorescence to mechanical loading[6,8]. Briefly, when GFP is mechanically loaded, it undergoes a fast, near-equilibrium transition from the native state to the first intermediate state that occurs at a characteristic force magnitude[8]. This first intermediate is characterized by the unfurling of GFP's flexible handle region, while all the beta strands in the beta-barrel remain fully intact. From this first intermediate state, GFP can transition to a second intermediate state, corresponding to beta-barrel disruption without complete denaturation[6]. A force-dependent rate constant was previously determined for this second transition by fitting to a Bell model[6]. Lastly, recovery of fluorescence requires complete unloading of GFP and return to the native state, with no recovery of fluorescence observed in the intermediate states along the refolding trajectory[6]. Taken together, this suggests that FP mechanical switching occurs through two subsequent transitions: a fast, near-equilibrium transition at a characteristic force, followed by a slower transition with a rate constant characterized by a Bell model. As recovery of fluorescence required complete unloading, we chose to model the second transition as not reversible and only permit recovery of fluorescence through protein unbinding and unloading (see above). Under these assumptions, we can approximate a single effective forward rate constant for FP mechanical switching. This consists of the probability of existing in the intermediate state produced from the first near-equilibrium sub-transition (modeled as a logistic function) multiplied by the rate constant of the second sub-transition (modeled as a Bell model). As such, we define the force-dependent rate constant of mechanical switching for an FP subject to a tension, $F$, as follows:

$$k_{MS}(F) = \left[1 + e^{-m\cdot(F-F_{1/2})}\right]^{-1} \cdot \left[k_{MS,0} \cdot e^{F\cdot\Delta x_{MS}/(k_BT)}\right] \quad \text{(S1)}$$

where the logistic function component is parameterized by a characteristic force $F_{1/2}$ and a steepness $m$, and the Bell model component is parametrized by an intrinsic rate constant $k_{MS,0}$ and an exponential parameter $\Delta x_{MS}$ (Figure S1b). For the logistic component motivated by the fast, near-equilibrium sub-transition, if we consider a system with two states (A and B) with a difference in free energy minima $\Delta G_{BA}^0$ and a mechanical reaction coordinate separation $\Delta x_{BA}^0$ that is assumed unaffected by applied force, then the equilibrium probability of existing in state B as function of $F$ is $P_{B,eq}(F) = \left[1 + e^{(\Delta G_{BA}^0 - F\Delta x_{BA}^0)/(k_BT)}\right]^{-1}$, derived from Bustamante et al.[9]. In this case, we have $F_{1/2} = \frac{\Delta G_{BA}^0}{\Delta x_{BA}^0}$ and $m = \frac{\Delta x_{BA}^0}{k_BT}$. Using state A to represent native GFP and state B to represent the first unfolding intermediate of GFP (GFPΔα), $\Delta G_{BA}^0$ would be approximately $22 k_BT$ as previously reported[8]. If we also estimate $\Delta x_{BA}^0$ from the previously reported contour length increase for this transition ($3.2\ nm$)[8], we get $F_{1/2} = 28.3\ pN$ and $m = 0.778\ pN^{-1}$ for GFP. The parameters for the single FP model are given in Table S1.

**Table S1: Parameters for Single FP Model**

| Param | Units | GFP Estimate | Single FP Model | Rationale |
|---|---|---|---|---|
| $F_{1/2}$ | pN | 28.3 | [2.83, 283] sweep | Sweep centered on estimated parameter for GFP from data in Dietz et al[8]. |
| $m$ | 1/pN | 0.778 | 0.778 | Estimated parameter for GFP from data in Dietz et al[8]. |
| $k_{MS,0}$ | 1/s | 0.33 | [0.033, 3.3] sweep | Sweep centered on estimated parameter for GFP from Ganim et al[6]. |
| $\Delta x_{MS}$ | nm | 0.23 | 0.23 | Estimated parameter for GFP from Ganim et al[6]. |
| $F$ | pN | N/A | [0,40] sweep | Encompasses estimated range of load magnitudes (~1-20 pN) based on molecular-scale force generation in the actin cytoskeleton driven by Myosin II motors[10,11] or F-actin polymerization[12], as well as estimates for the extension of molecular clutches (with characteristic protein stiffnesses) by actin retrograde flow[3,13] (Figure S1c, x-axis). Full range of parameter sweep extends to load magnitudes twice as high for completeness. |
| $\tau$ | s | N/A | [.01,100] sweep | Encompasses estimated range of load durations (~0.1-10 seconds) based on the unbinding lifetimes of bonds in mechanical linker proteins and transmembrane proteins under this range of load magnitudes, including alpha-catenin:F-actin[14], vinculin:F-actin[15], talin:F-actin[16], integrin:FN[17], E-cadheren trans-dimer[18], alpha-actinin:F-actin[19] and filamin:F-actin[19] (Figure S1c, y-axis). Full range of parameter sweep extends below and above estimated range by 1 order of magnitude for completeness. |

#### 2. Steady State Analysis of Model

To characterize the kinetic model for FP mechanical switching in a load-bearing protein, a steady state analysis in the deterministic limit was performed using 2 ordinary differential equations describing the rate of change of concentration of each species and 1 conservation equation for total concentration of protein.

$$\frac{d[FP1]}{dt} = -k_{MS}(F) \cdot [FP1] + k_{unbind}(F) \cdot [FP0]$$

$$\frac{d[FP0]}{dt} = -k_{unbind}(F) \cdot [FP0] + k_{MS}(F) \cdot [FP1]$$

$$[Total\ FP] = [FP1] + [FP0]$$

Setting each ODE to zero and solving for the steady state concentrations of each species yields expressions for the steady state fraction ($\rho$) of each species:

$$\rho_{FP1}(F) = \frac{[FP1]_{ss}}{[FP1]_{ss} + [FP0]_{ss}} = \frac{k_{unbind}}{k_{unbind} + k_{MS}(F)} \quad \text{(S2)}$$

$$\rho_{FP0}(F) = \frac{[FP0]_{ss}}{[FP1]_{ss} + [FP0]_{ss}} = \frac{k_{MS}(F)}{k_{unbind} + k_{MS}(F)} \quad \text{(S3)}$$

#### B. Results

To characterize the process of FP mechanical switching inside a dynamic load-bearing protein, we first investigated the range of load magnitudes ($F$) and durations ($\tau$) over which FP mechanical switching is likely to occur. We focused on an estimated range of load magnitudes (~1-20 pN) based on molecular-scale force generation in the actin cytoskeleton driven by Myosin II motors[10,11] or F-actin polymerization[12], as well as estimates for the extension of molecular clutches (with characteristic protein stiffnesses) by actin retrograde flow[3,13] (Figure S1c, x-axis). We also note that this range encompasses the equilibrium unfolding forces of multiple endogenous mechanosensitive protein domains, including those in alpha-catenin and talin[20,21]. We focused on an estimated range of load durations (~0.1-10 seconds) based on the unbinding lifetimes of bonds in mechanical linker proteins and transmembrane proteins under this range of load magnitudes, including alpha-catenin:F-actin[14], vinculin:F-actin[15], talin:F-actin[16], integrin:FN[17], E-cadheren trans-dimer[18], alpha-actinin:F-actin[19] and filamin:F-actin[19] (Figure S1c, y-axis). As a large set of natural and engineered FPs with different structures, photophysical properties, and mechanical stabilities exist, and the integration of FPs into a fusion protein as well as the use of FPs in the cellular environment can alter these properties[8,22-24], we assessed FPs with a range of force thresholds ($F_{1/2}$) and kinetic timescales ($1/k_{ms,0}$) (Figure S1d-o).

For a given set of FP parameters, FP mechanical switching occurs at load magnitudes near or above the force threshold for FP mechanical switching ($F \geq F_{1/2}$) and load durations of the same order of magnitude or longer than the kinetic timescale for FP mechanical switching ($\tau \geq 1/k_{ms,0}$) (for example see Figure S1h). If the load magnitude is too low, then FP mechanical switching is not permitted regardless of load duration. Likewise, when the load duration is too short, the protein unbinds before FP mechanical switching occurs, regardless of load magnitude.

Comparing FPs with different parameters, we find that decreasing the force threshold ($F_{1/2}$) expands the range of forces over which FP mechanical switching occurs with little effect on the range of load durations (for example compare Figure S1e,h,k,n). Alternatively, increasing the kinetic rate constant ($k_{ms,0}$) expands the range of load durations that support FP mechanical switching without affecting the load magnitude range (for example compare Figure S1g-i). Therefore, for two FPs with different sets of FP mechanical switching parameters, there exists load magnitudes and/or durations for which neither, just one, or both FPs are mechanically switched.

Lastly, this model also demonstrates differential sensitivity of FP mechanical switching to load magnitude and duration. For a given set of FP parameters, the fraction of mechanically switched FPs changes rapidly with respect to force near the force threshold, increasing steeply from 0 to 1 over a small range of a few pN in load magnitude (less than an order of magnitude; exact value depends on the parameters $m$ and $\Delta x_{MS}$; for an example see the horizontal arrow in Figure S1h). In contrast, the fraction of mechanically switched FPs changes gradually with respect to load duration, increasing gradually from 0 to 1 over almost 2 orders of magnitude in load duration (for an example see the vertical arrow in Figure S1h).

Taken together, the model of FP mechanical switching in a load-bearing protein indicates that FP mechanical switching can occur at the load magnitudes and durations estimated for protein loading in cells for a range of FP parameters. For a given FP, the loads must be high enough ($F \geq F_{1/2}$) and long enough ($\tau \geq 1/k_{ms,0}$) to support mechanical switching. Lastly, the process of FP mechanical switching exhibits sensitivity to both load magnitude and duration.

### III. Model of FP Mechanical Switching in FRET-based Tension Sensor

In this section we investigate how FP mechanical switching affects FRET-based MTSs. To do so, we extend the model of FP mechanical switching in a load-bearing protein to a FRET-based MTS containing a donor and an acceptor FP. Then, we compute three channel FRET measurements of ensembles of MTSs exhibiting donor and/or acceptor mechanical switching.

#### A. Model Formulation

##### 1. Kinetic Model

An MTS consists of a tension sensor module composed of two FPs (donor and acceptor) separated by an extensible domain that is integrated into a load-bearing protein. The MTS is subject to the same loading dynamics previously described for load-bearing proteins in Section II.A (Figure S2a). As the FPs are in series in the line of loading, the load applied across the MTS is experienced by both the donor and acceptor FPs, each of which can undergo FP mechanical switching as previously described in Section II.A. As a large set of natural and engineered FPs with different structures, photophysical properties, and mechanical stabilities exist[8,22], we model separate FP mechanical switching kinetic parameters for the acceptor and donor. The force-dependent rate constant of mechanical switching for donor and acceptor FPs subject to a load, $F$, are thus defined as follows:

$$k_{MS}^{D}(F) = \left[1 + e^{-m^{D}\cdot(F - F_{1/2}^{D})}\right]^{-1} \cdot \left[k_{MS,0}^{D} \cdot e^{F\cdot \Delta x_{MS}^{D}/(k_B T)}\right] \quad \text{(S4)}$$

$$k_{MS}^{A}(F) = \left[1 + e^{-m^{A}\cdot(F - F_{1/2}^{A})}\right]^{-1} \cdot \left[k_{MS,0}^{A} \cdot e^{F\cdot \Delta x_{MS}^{A}/(k_B T)}\right] \quad \text{(S5)}$$

Because the donor and acceptor FP can each exist in two states, i.e. functional/on (FP1) and non-functional/off (FP0), an MTS can exist in one of four states: functional donor and functional acceptor (D1A1), functional donor and non-functional acceptor (D1A0), non-functional donor and functional acceptor (D0A1), and non-functional donor and non-functional acceptor (D0A0). The transitions between these four states are summarized in the transition rate diagram (Figure S2b). Note that the absence of reverse transitions from D0A0 to D1A0 or from D0A0 to D0A1 is a consequence of the simple assumption that FP fluorescence does not recover under load. The parameters for the MTS model are given in Table S2. Values of $m^D$ and $m^A$ were adjusted based on the values of $F_{1/2}^{D}$ and $F_{1/2}^{A}$, respectively, to match the experimental observation of negligeable FP mechanical switching in unloaded FPs[6] as well as MTSs (see the following load-insensitive experimental controls from this work: ABDTL in

Figure 2 and Figure S13, and VinTS-I997A in Figure 3 and Figure S19). Specifically, $m^D$ and $m^A$ were set to limit the fraction of mechanically switched FPs in unloaded MTSs to negligeable values of $\rho_{D0}(F=0) < 0.0005$ and $\rho_{A0}(F=0) < 0.0005$ without reducing $m^D$ and $m^A$ below a lower limit of $1\ pN^{-1}$ based on the estimated parameter for GFP[8] (see Section II.A). Mathematically, this means that $m = max\left\{1,\ \frac{1}{F_{1/2}}\ln\left(\frac{1}{.0005}-1\right)\right\}\ pN^{-1}$.

**Table S2: Parameters for MTS Model**

| Param | Units | Base Value | Param Sweep | Rationale |
|---|---|---|---|---|
| $F_{1/2}^D$ $F_{1/2}^A$ | pN | 5 | 1, 3, 5, 8, 25 | Base value is permissive of FP mechanical switching in the estimated range of load magnitudes for molecular-scale forces. Sweep covers range below and above base value, with highest value similar to estimated parameter for GFP from Dietz et al.[8] (see Table S1) |
| $m^D$ $m^A$ | 1/pN | 1.5 | 7.6, 2.5, 1.5, 1, 1 for $F_{1/2}$ values of 1, 3, 5, 8, 25, respectively | Adjusted to match the experimental observation of negligeable FP mechanical switching in unloaded MTSs. See text for further information. |
| $k_{MS,0}^D$ $k_{MS,0}^A$ | 1/s | 1 | [.1,10] | Base value similar to parameter for GFP from Ganim et al.[8] (see Table S1). Sweep covers one order of magnitude around the base value. |
| $\Delta x_{MS}^D$ $\Delta x_{MS}^A$ | nm | .23 | .23 | Parameter for GFP from Ganim et al.[6] (see Table S1) |
| $F$ | pN | N/A | [0,30] | Encompasses estimated range of load magnitudes (~1-20 pN) based on molecular-scale force generation in the actin cytoskeleton driven by Myosin II motors[10,11] or F-actin polymerization[12], as well as estimates for the extension of molecular clutches (with characteristic protein stiffnesses) by actin retrograde flow[3,13] (Figure S1c, x-axis). Full range of parameter sweep extends to load magnitudes 1.5x as high for completeness. |
| $\tau$ | s | 1 | [.01,100] | Encompasses estimated range of load durations (~0.1-10 seconds) based on the unbinding lifetimes of bonds in mechanical linker proteins and transmembrane proteins under this range of load magnitudes, including alpha-Catenin:F-actin[14], Vinculin:F-actin[15], Talin:F-actin[16], Integrin:FN[17], E-cad trans-dimer[18], alpha-Actinin:F-actin[19] and Filamin:F-actin[19] (Figure S1c, y-axis). Full range of parameter sweep extends below and above estimated range by 1 order of magnitude for completeness. |

#### 2. Steady State Analysis of Kinetic Model

To characterize the kinetic model for FP mechanical switching in an MTS, a steady state analysis in the continuous deterministic limit was performed using 4 ordinary differential equations describing the rate of change of concentration of each species and 1 conservation equation for total concentration of MTS.

$$\frac{d[D1A1]}{dt} = -\left(k_{MS}^{D}(F) + k_{MS}^{A}(F)\right) \cdot [D1A1] + k_{unbind} \cdot ([D0A1] + [D0A1] + [D0A0])$$

$$\frac{d[D0A1]}{dt} = -(k_{MS}^{A}(F) + k_{unbind}) \cdot [D0A1] + k_{MS}^{D}(F) \cdot [D1A1]$$

$$\frac{d[D1A0]}{dt} = -(k_{MS}^{D}(F) + k_{unbind}) \cdot [D1A0] + k_{MS}^{A}(F) \cdot [D1A1]$$

$$\frac{d[D0A0]}{dt} = -(k_{unbind}) \cdot [D0A0] + k_{MS}^{A}(F) \cdot [D0A1] + k_{MS}^{D}(F) \cdot [D1A0]$$

$$[Total\ MTS] = [D1A1] + [D0A1] + [D1A0] + [D0A0]$$

Setting each ODE to zero and solving for the steady state concentrations of each species yields expressions for the steady state fraction ($\rho$) of each species:

$$\rho_{D1A1}(F) = \frac{[D1A1]_{ss}}{[Total\ MTS]} = \frac{k_{unbind}}{k_{MS}^{D}(F) + k_{MS}^{A}(F) + k_{unbind}} \tag{S6}$$

$$\rho_{D0A1}(F) = \frac{[D0A1]_{ss}}{[Total\ MTS]} = \frac{k_{MS}^{D}(F) \cdot k_{unbind}}{[k_{MS}^{A}(F) + k_{unbind}][k_{MS}^{D}(F) + k_{MS}^{A}(F) + k_{unbind}]} \tag{S7}$$

$$\rho_{D1A0}(F) = \frac{[D1A0]_{ss}}{[Total\ MTS]} = \frac{k_{MS}^{A}(F) \cdot k_{unbind}}{[k_{MS}^{D}(F) + k_{unbind}][k_{MS}^{D}(F) + k_{MS}^{A}(F) + k_{unbind}]} \tag{S8}$$

$$\rho_{D0A0}(F) = \frac{[D0A0]_{ss}}{[Total\ MTS]} = \frac{k_{MS}^{D}(F) \cdot k_{MS}^{A}(F) \cdot (k_{MS}^{D}(F) + k_{MS}^{A}(F) + 2k_{unbind})}{[k_{MS}^{A}(F) + k_{unbind}][k_{MS}^{D}(F) + k_{unbind}][k_{MS}^{D}(F) + k_{MS}^{A}(F) + k_{unbind}]} \tag{S9}$$

Additionally, the total steady state fraction of MTSs with mechanically switched donor or acceptor can also be obtained:

$$\rho_{D0}(F) = \frac{[D0A1]_{ss} + [D0A0]_{ss}}{[Total\ MTS]} = \frac{k_{MS}^{D}(F)}{k_{MS}^{D}(F) + k_{unbind}} \tag{S10}$$

$$\rho_{A0}(F) = \frac{[D1A0]_{ss} + [D0A0]_{ss}}{[Total\ MTS]} = \frac{k_{MS}^{A}(F)}{k_{MS}^{A}(F) + k_{unbind}} \tag{S11}$$

#### 3. Computation of Three Channel FRET Measurements of MTS Ensemble

We focused on sensitized emission as the FRET imaging modality because it is widely used and there are existing approaches for calibrating, measuring, and analyzing the relative abundance of acceptor and donor fluorophores[25-27]. In sensitized emission-based FRET measurements, images are acquired in three channels (AA: acceptor excitation and acceptor emission; DD: donor excitation and donor emission; DA: donor excitation and acceptor emission)[28]. With calibration, the FRET efficiency, $E$, and FP stoichiometry, $S = n_D/(n_D + n_A)$, can be determined from the signal in these three channels[27]. To simulate measurements of MTSs with FP mechanical switching, we derive general expressions for the three channel FRET signals and resulting apparent FRET Efficiency, $E_{app}$, and Stoichiometry, $S_{app}$, for a population of MTSs. In a given population, each MTS exists in a state, $\psi_i$, based on the functional status of its donor and acceptor fluorescent protein ($\psi_i = \mathrm{D1A1, D1A0, D0A1, D0A0}$) and is subject to a force, $F_i$.

**Signal Contributions for Each Sensor State**. Sensors in the D1A1 state ($\psi_i = D1A1$) undergo intramolecular FRET with a FRET efficiency ($E_i$), which depends on molecular tension according to the FRET Eff vs force calibration of the tension sensor module[24,29], i.e. $E_i = f(F_i)$. Here, the FRET-force relationship for the original TSMod (mTFP1-(GPGGA)$_8$-mVenus; Figure S2c) was used to facilitate comparisons to experimental data in this work[24,29]. However, the framework here can be adapted to other calibrated tension sensor modules by modifying the FRET-force relationship (see Section VI for a complete discussion of this applicability). To our knowledge, forced-induced changes in the excitation or emission wavelengths of FPs have not been described. Therefore, to determine the signal contribution for each sensor state, we assumed that donor FPs that have undergone mechanical switching cannot be excited by any excitation light in the optical system, and that acceptor FPs that have undergone mechanical switching cannot be excited by any excitation light in the optical system and also cannot accept energy from donor FPs. As such, sensors with mechanically switched acceptors (D1A0) do not undergo FRET and behave like free donors. Sensors with mechanically switched donors (D0A1) do not undergo FRET and behave like free acceptors. Sensors with both donor and acceptor mechanically switched (D0A0) do not undergo FRET and do not affect any signals. Additionally, we assume the absence of intermolecular FRET between sensors, regardless of sensor state. Using previously defined photophysical and instrumental parameters[27], the signal contribution from each sensor state in each image channel is therefore defined in Table S3, where $E_i = f(F_i)$ is the FRET Efficiency (in the D1A1 state) for the ith sensor, $\phi_j$ is the fluorescence quantum yield of fluorophore $j$, $L_j$ is the excitation intensity for the excitation of fluorophore $j$, $\sigma_k^j$ is the absorption cross section of fluorophore $j$ when excited with excitation channel $k$, $\eta_k^j$ is the efficiency of detecting photons emitted by fluorophore $j$ in the detection channel $k$.

**Table S3: Signal Contributions from Each Sensor State**

| State | AA Signal Per Sensor | DD Signal Per Sensor | DA Signal Per Sensor | | |
|---|---|---|---|---|---|
| | | | Sensitized Emission | Bleedthrough of Donor | Direct Acceptor Excitation |
| D1A1 | $L_A\sigma_{Aex}^A\phi_A\eta_{Adet}^{Aem}$ | $(1-E_i)\cdot(L_D\sigma_{Dex}^D\phi_D\eta_{Ddet}^{Dem})$ | $E_i\cdot(L_D\sigma_{Dex}^D\phi_A\eta_{Adet}^{Aem})$ | $(1-E_i)\cdot(L_D\sigma_{Dex}^D\phi_D\eta_{Adet}^{Dem})$ | $L_D\sigma_{Dex}^A\phi_A\eta_{Adet}^{Aem}$ |
| D1A0 | 0 | $L_D\sigma_{Dex}^D\phi_D\eta_{Ddet}^{Dem}$ | 0 | $L_D\sigma_{Dex}^D\phi_D\eta_{Adet}^{Dem}$ | 0 |
| D0A1 | $L_A\sigma_{Aex}^A\phi_A\eta_{Adet}^{Aem}$ | 0 | 0 | 0 | $L_D\sigma_{Dex}^A\phi_A\eta_{Adet}^{Aem}$ |
| D0A0 | 0 | 0 | 0 | 0 | 0 |

These expressions can be simplified by combining the photophysical and instrumental parameters into 4 constants[27]. These (or similar) constants are routinely determined by three channel FRET calibration procedures and enable the determination of FRET efficiency from sensitized emission[25-27]. The donor bleedthrough constant, $\alpha^{BT}$, relates to the bleedthrough of photons emitted by the donor into the DA channel, and the acceptor direction excitation constant, $\delta^{DE}$, relates to the photons from the direct excitation of the acceptor in the DA channel. They are given below:

$$\alpha^{BT} \equiv \frac{\eta_{Ddet}^{Dem}}{\eta_{Adet}^{Dem}}, \qquad \delta^{DE} \equiv \frac{L_D \sigma_{Dex}^{A}}{L_A \sigma_{Aex}^{A}}$$

Additionally, a factor for the different detection efficiencies in both channels, $\gamma^M$, and a factor for the different excitation efficiencies, $\beta^X$ are defined as follows:

$$\gamma^M \equiv \frac{\phi_A \eta_{Adet}^{Aem}}{\phi_D \eta_{Ddet}^{Dem}}, \qquad \beta^X \equiv \frac{L_A \sigma_{Aex}^{A}}{L_D \sigma_{Dex}^{D}}$$

Using these four constants and an intensity scaling constant representing the signal intensity of a single acceptor fluorophore in the AA channel, $C_{AA} \equiv L_A \sigma_{Aex}^{A} \phi_A \eta_{Adet}^{Aem}$, we simplify the signal contribution from each sensor state in each image channel, yielding the expressions in Table S4.

**Table S4: Signal Contributions for Each Sensor State with Simplified Parameters**

| State | AA Signal Per Sensor | DD Signal Per Sensor | DA Signal Per Sensor | | |
|---|---|---|---|---|---|
| | | | Sensitized Emission | Bleedthrough of Donor | Direct Acceptor Excitation |
| D1A1 | $C_{AA}$ | $(1-E_i)\cdot\frac{C_{AA}}{\gamma^M\beta^X}$ | $E_i\cdot\frac{C_{AA}}{\beta^X}$ | $(1-E_i)\cdot\frac{\alpha_{BT}C_{AA}}{\gamma^M\beta^X}$ | $\delta_{DE}\cdot C_{AA}$ |
| D1A0 | 0 | $\frac{C_{AA}}{\gamma^M\beta^X}$ | 0 | $\frac{\alpha_{BT}C_{AA}}{\gamma^M\beta^X}$ | 0 |
| D0A1 | $C_{AA}$ | 0 | 0 | 0 | $\delta_{DE}\cdot C_{AA}$ |
| D0A0 | 0 | 0 | 0 | 0 | 0 |

From the signal contributions for each sensor state in each image channel, we derive the total signal in each image channel ($I_{AA}^{TOT}, I_{DD}^{TOT}, I_{DA}^{TOT}$) for a population of sensors existing in these four states.

$$I_{AA}^{TOT} = \sum_{D1A1} C_{AA} + \sum_{D0A1} C_{AA} \tag{S12}$$

$$I_{DD}^{TOT} = \sum_{D1A1} (1-E_i)\cdot\frac{C_{AA}}{\gamma^M\beta^X} + \sum_{D1A0} \frac{C_{AA}}{\gamma^M\beta^X} \tag{S13}$$

$$I_{DA}^{TOT} = \sum_{D1A1}\left(E_i\cdot\frac{C_{AA}}{\beta^X} + (1-E_i)\cdot\frac{\alpha_{BT}C_{AA}}{\gamma^M\beta^X} + \delta_{DE}\cdot C_{AA}\right) + \sum_{D1A0}\frac{\alpha_{BT}C_{AA}}{\gamma^M\beta^X} + \sum_{D0A1}\delta_{DE}\cdot C_{AA} \tag{S14}$$

where $E_i$ is the FRET Efficiency (in the D1A1 state) for the ith sensor, and $\sum_{\psi_m}[\dots]$ denotes the sum over all sensors in state $\psi_m$.

**Computation of Corrected FRET.** In three channel FRET measurements, the DA channel is subject to bleedthrough of photons emitted by the donor and photons resulting from the direct excitation of the acceptor. We do not consider crosstalk between donor and acceptor here because microscope hardware is typically specified to make these effects negligeable[25-27]. Therefore, the corrected FRET intensity,

$I_{DA\,corr}^{TOT}$ (also indicated using the variable $F_C$ in other sensitized emission FRET formalisms) is defined as follows:

$$I_{DA\,corr}^{TOT} \equiv I_{DA}^{TOT} - \widehat{\alpha_{BT}} \cdot I_{DD}^{TOT} - \widehat{\delta_{DE}} \cdot I_{AA}^{TOT} \tag{S15}$$

where $\widehat{\alpha_{BT}}$ is the estimated donor bleedthrough constant and $\widehat{\delta^{DE}}$ is the estimated acceptor direction excitation constant, which are determined experimentally using samples containing only donor fluorophores or only acceptor fluorophores[25-27]. As cytosolic FPs are not mechanically loaded, and these constants are routinely determined using standard three channel FRET calibration procedures[25-27], we assume $\widehat{\alpha_{BT}} = \alpha^{BT}$ and $\widehat{\delta^{DE}} = \delta^{DE}$, i.e. no estimation error in these calibration constants. Using this result, the expression for $I_{DA\,corr}^{TOT}$ can be further simplified:

$$\begin{aligned} I_{DA\,corr}^{TOT} &\equiv I_{DA}^{TOT} - \widehat{\alpha_{BT}} \cdot I_{DD}^{TOT} - \widehat{\delta_{DE}} \cdot I_{AA}^{TOT} \\ &= \left[\sum_{D1A1}\left(E_i \cdot \frac{C_{AA}}{\beta^X} + (1-E_i)\cdot\frac{\alpha_{BT}C_{AA}}{\gamma^M\beta^X} + \delta_{DE}\cdot C_{AA}\right) + \sum_{D1A0}\frac{\alpha_{BT}C_{AA}}{\gamma^M\beta^X} + \sum_{D0A1}\delta_{DE}\cdot C_{AA}\right] - \alpha^{BT}\left[\sum_{D1A1}(1-E_i)\cdot\frac{C_{AA}}{\gamma^M\beta^X} + \sum_{D1A0}\frac{C_{AA}}{\gamma^M\beta^X}\right] \\ &\qquad - \delta^{DE}\left[\sum_{D1A1}C_{AA} + \sum_{D0A1}C_{AA}\right] \\ &= \sum_{D1A1}E_i\cdot\frac{C_{AA}}{\beta^X} \end{aligned}$$

Note that this demonstrates that sensors with non-functional donor and/or acceptor (sensors in the D1A0, D0A1, or D0A0 states) do not affect the corrected FRET signal.

**Computation of Apparent FRET Efficiency and Stoichiometry.** To estimate the apparent FRET Efficiency and Stoichiometry of a FRET sensor, two additional correction factors, $\widehat{\gamma^M}$ and $\widehat{\beta^X}$ , must be determined for the microscope setup and fluorescent protein pair[27]. When these constants are determined, the apparent FRET Efficiency and Stoichiometry are evaluated as follows:

$$E_{app} \equiv \frac{I_{DA\,corr}^{TOT}}{I_{DA\,corr}^{TOT} + \widehat{\gamma^M} I_{DD}^{TOT}} \tag{S16}$$

$$S_{app} \equiv \frac{I_{DA\,corr}^{TOT} + \widehat{\gamma^M} I_{DD}^{TOT}}{I_{DA\,corr}^{TOT} + \widehat{\gamma^M} I_{DD}^{TOT} + I_{AA}^{TOT}/\widehat{\beta^X}} \tag{S17}$$

The constants $\widehat{\gamma^M}$ and $\widehat{\beta^X}$ are routinely determined using standard three channel FRET calibration procedures[25-27]. These calibrations use cytosolic FRET constructs, which are unloaded, so we assume $\widehat{\gamma^M} = \gamma^M$ and $\widehat{\beta^X} = \beta^X$, i.e. no estimation error in these calibration constants.

**Expressions for Apparent FRET Efficiency and Stoichiometry for MTS Population with All Sensors Under the Same Tension and No Imaging Noise.** We now derive expressions for the apparent FRET Efficiency and Stoichiometry of a sensor population in which all sensors are subject to the same force, i.e. $F_i = F_0$

for all sensors. Therefore, all sensors have the same FRET Efficiency in the D1A1 state, i.e. $E_i = E_0$ for all sensors in the D1A1 state, where $E_0 = f(F_0)$ as defined previously. For a mixed population of MTSs containing a specified number of sensors in each state ($n_{D1A1}, n_{D1A0}, n_{D0A1}, n_{D0A0}$) and having a non-zero number of sensors in the D1A1 state ($n_{D1A1} > 0$), the apparent FRET efficiency and stoichiometry can be simplified to the expressions given by Equations S18 and S19 in this section. First, we derive apparent FRET efficiency, $E_{app}$:

$$\begin{aligned}
E_{app} &\equiv \frac{I_{DA\,corr}^{TOT}}{I_{DA\,corr}^{TOT} + \widehat{\gamma^M} I_{DD}^{TOT}} \\
&= \frac{\left[\sum_{D1A1} E_i \cdot \frac{C_{AA}}{\beta^X}\right]}{\left[\sum_{D1A1} E_i \cdot \frac{C_{AA}}{\beta^X}\right] + \gamma^M \left[\sum_{D1A1} (1 - E_i) \cdot \frac{C_{AA}}{\gamma^M \beta^X} + \sum_{D1A0} \frac{C_{AA}}{\gamma^M \beta^X}\right]} \\
&= \frac{E_0 \cdot n_{D1A1} \cdot \frac{C_{AA}}{\beta^X}}{E_0 \cdot n_{D1A1} \cdot \frac{C_{AA}}{\beta^X} + \gamma^M \left([n_{D1A1} \cdot (1 - E_0) + n_{D1A0}] \frac{C_{AA}}{\gamma^M \beta^X}\right)} \\
&= \frac{E_0 \cdot n_{D1A1}}{E_0 \cdot n_{D1A1} + [n_{D1A1} \cdot (1 - E_0) + n_{D1A0}]} \\
&= \frac{E_0}{1 + \left(\frac{n_{D1A0}}{n_{D1A1}}\right)}
\end{aligned}$$

$$E_{app} = \frac{E_0}{1 + \left(\frac{n_{D1A0}}{n_{D1A1}}\right)} \qquad \text{(S18)}$$

where $E_0 = f(F_0)$. This demonstrates that $E_{app}$ depends only on the number of sensors in the D1A1 state ($n_{D1A1}$) and sensors with non-functional acceptor ($n_{D1A0}$), and not on the number of sensors with non-functional donor ($n_{D0A1}$) or both FPs non-functional ($n_{D0A0}$). Therefore, $E_{app}$ is only affected by acceptor mechanical switching. This is similar to previous work showing that free donors, but not free acceptors, affect $E_{app}$ when mixed with a population of sensors in the D1A1 state[27]. Indeed, for the case of a sensor whose FRET is insensitive to force, Equation S18 is equivalent to the expression derived by Coullomb et al.[27] for $E_{app}$ for a mixed population of ideal sensors ($n_0^D = n_{D1A1}$) and free donors ($n_{free}^D = n_{D1A0}$ ).

Next, we derive apparent Stoichiometry, $S_{app}$:

$$\begin{aligned}
S_{app} &\equiv \frac{I_{DA\,corr}^{TOT} + \widehat{\gamma^M} I_{DD}^{TOT}}{I_{DA\,corr}^{TOT} + \widehat{\gamma^M} I_{DD}^{TOT} + \frac{I_{AA}^{TOT}}{\widehat{\beta^X}}} \\
&= \frac{\left[\sum_{D1A1} E_i \cdot \frac{C_{AA}}{\beta^X}\right] + \gamma^M \left[\sum_{D1A1} (1 - E_i) \cdot \frac{C_{AA}}{\gamma^M \beta^X} + \sum_{D1A0} \frac{C_{AA}}{\gamma^M \beta^X}\right]}{\left[\sum_{D1A1} E_i \cdot \frac{C_{AA}}{\beta^X}\right] + \gamma^M \left[\sum_{D1A1} (1 - E_i) \cdot \frac{C_{AA}}{\gamma^M \beta^X} + \sum_{D1A0} \frac{C_{AA}}{\gamma^M \beta^X}\right] + \frac{1}{\beta^X} \left[\sum_{D1A1} C_{AA} + \sum_{D0A1} C_{AA}\right]} \\
&= \frac{E \cdot n_{D1A1} \cdot \frac{C_{AA}}{\beta^X} + \gamma^M \left([n_{D1A1} \cdot (1 - E) + n_{D1A0}] \frac{C_{AA}}{\gamma^M \beta^X}\right)}{E \cdot n_{D1A1} \cdot \frac{C_{AA}}{\beta^X} + \gamma^M \left([n_{D1A1} \cdot (1 - E) + n_{D1A0}] \frac{C_{AA}}{\gamma^M \beta^X}\right) + \frac{(n_{D1A1} + n_{D0A1}) \cdot C_{AA}}{\beta^X}}
\end{aligned}$$

$$= \frac{E \cdot n_{D1A1} + n_{D1A1} \cdot (1-E) + n_{D1A0}}{E \cdot n_{D1A1} + n_{D1A1} \cdot (1-E) + n_{D1A0} + (n_{D1A1} + n_{D0A1})}$$

$$= \frac{(n_{D1A1} + n_{D1A0})}{(n_{D1A1} + n_{D1A0}) + (n_{D1A1} + n_{D0A1})}$$

$$S_{app} = \frac{n_{D1A1} + n_{D1A0}}{(n_{D1A1} + n_{D1A0}) + (n_{D1A1} + n_{D0A1})} \tag{S19}$$

This demonstrates that $S_{app}$ depends on the number of sensors in the D1A1 state ($n_{D1A1}$) and the number of sensors with either non-functional acceptor ($n_{D1A0}$) or non-functional donor ($n_{D0A1}$). Therefore, as expected, $S_{app}$ can be affected by both acceptor and donor mechanical switching. This is again consistent with previous work showing that free donor or free acceptor can both affect $S_{app}$ when mixed with a population of sensors in the D1A1 state[27]. By combining the total number of functional donor ($n_D = n_{D1A1} + n_{D1A0}$) and acceptor FPs ($n_A = n_{D1A1} + n_{D0A1}$), we also show that $S_{app} = \frac{n_D}{n_D + n_A}$, which means that $S_{app}$ remains a true readout of functional FP stoichiometry in the presence of mechanical switching of the acceptor and/or donor.

To validate the derived expressions for $E_{app}$ and $S_{app}$ (Equations S18 and S19), we confirmed exact agreement with values of $E_{app}$ and $S_{app}$ computed directly from simulated raw channel intensities using Equations S12-S17. Validation results for different populations of sensor states ($n_{D1A1}, n_{D1A0}, n_{D0A1}, n_{D0A0}$) and forces $F$ are shown in Figure S3. The validations shown in Figure S3 were performed with FRET calibration constants similar to those for the mTFP1-mVenus FRET pair and microscope setup used in our experimental work ($\widehat{\alpha_{BT}} = 0.75, \widehat{\delta^{DE}} = 0.25, \widehat{\gamma^{M}} = 1.65, \widehat{\beta^{X}} = 0.6061$) and with intensity values ($C_{AA} = 100$) similar to those in our experimental system. However, the derived expressions hold for all FRET calibration constants and raw channel intensity values.

### B. Results

#### 1. Development of a Framework to Assess FP Mechanical Switching in MTS

To visualize MTS data in the presence of FP mechanical switching, we sought to adapt a recently developed framework for plotting three-channel FRET data using the apparent FRET efficiency, $E_{app}$, and apparent stoichiometry, $S_{app}$[27]. MTSs have a continuously variable FRET efficiency in the D1A1 state, $E_0(F)$, that depends on the molecular tension according to a FRET efficiency versus tension calibration curve[24,29] (Figure S4a). Therefore, in the absence of FP mechanical switching, increasing tension leads to a decrease in $E_{app}$ with a constant $S_{app} = 0.5$ (Figure S4b). In this case, $E_{app}$ for an ensemble of MTSs under a single tension $F$ remains equal to the calibrated value, $E_0(F)$. In the presence of FP mechanical switching, deviations from $S_{app} = 0.5$ occur. Acceptor mechanical switching increases $S_{app}$ and decreases $E_{app}$ moving the ($E_{app}$, $S_{app}$) point up and left (Figure S4c, dots indicate different distributions of D1A1 and D1A0 states for 3 pN). Donor mechanical switching decreases $S_{app}$ and does not affect $E_{app}$, moving the ($E_{app}$, $S_{app}$) point down (Figure S4d, dots indicate different distributions of D1A1 and D0A1 states for 3 pN). To establish references, we define tension isoclines as the curves containing ($E_{app}$, $S_{app}$) points for an MTS ensemble under a single tension $F$ (and thus having a single $E_0$) with all levels of acceptor mechanical switching only or donor mechanical switching only.

Taken together, this establishes a framework for visualizing three channel FRET measurements of MTSs using $E_{app}$ and $S_{app}$. This framework applies generally to MTSs containing any type of non-functional acceptor or donor FPs. We assess its suitability specifically for detecting FP mechanical switching in MTSs in the following sections.

#### 2. Effect of Acceptor Mechanical Switching in Dynamic MTSs

We next assessed the suitability of this framework for detecting FP mechanical switching in MTSs that undergo dynamic binding, loading, and unbinding. To do so, we used the kinetic model of FP mechanical switching in dynamic MTSs to obtain the steady state fractions of sensors in each of the four possible states for an MTS population subject to a specified load magnitude and duration and then computed the resulting ($E_{app}$, $S_{app}$) value. First, we analyzed computed FRET measurements for populations of MTSs undergoing acceptor mechanical switching only.

To assess the ability to detect acceptor mechanical switching inside MTS, we compared readouts for an MTS with no acceptor mechanical switching or other loss-of-function (Figure S5c, brown line), an MTS with force-independent acceptor loss-of-function such as due to photobleaching (Figure S5c, grey line), and an MTS with acceptor mechanical switching (Figure S5c, black line; base parameters in Table S2). We looked at ($E_{app}$, $S_{app}$)-curves for load magnitude $F$ from 0 to 30 pN and a load duration $\tau$ of 1 second. In the absence of acceptor mechanical switching (Figure S5c, brown line), the ($E_{app}$, $S_{app}$)-curve starts at the point ($E_{app} = E_0(F = 0) = 0.286$ , $S_{app} = 0.5$), corresponding to an unloadable tension sensor module, and as the load magnitude increases $E_{app}$ decreases while $S_{app}$ remains constant. Thus, an MTS without acceptor mechanical switching exhibits a horizontal ($E_{app}$, $S_{app}$) signature at $S_{app} = 0.5$. In the case of force-independent acceptor loss-of-function (Figure S5c, grey line), the ($E_{app}$, $S_{app}$)-curve starts at a lower $E_{app}$ and a higher $S_{app}$ than that of the unloaded tension sensor module. As load magnitude increases, $E_{app}$ decreases but $S_{app}$ does not change. Thus, an MTS with force-independent acceptor loss-of-function exhibits a horizontal ($E_{app}$, $S_{app}$) signature at $S_{app} > 0.5$. For the case of acceptor mechanical switching (Figure S5c, black line), the ($E_{app}$, $S_{app}$)-curve again starts at the point corresponding to the unloaded tension sensor module. While $E_{app}$ again decreases with load magnitude, $S_{app}$increases with load magnitude for loads near and above the FP's threshold force ($F \geq F^A_{1/2}$). Thus, acceptor mechanical switching exhibits an up/left slanting ($E_{app}$, $S_{app}$) signature. Therefore, these data indicate that acceptor mechanical switching has a unique ($E_{app}$, $S_{app}$) signature that can be distinguished from cases of no acceptor mechanical switching as well as force-independent acceptor loss-of-function.

A large set of natural and engineered FPs with different structures, photophysical properties, and mechanical stabilities exist, and the integration of FPs into a fusion protein as well as the use of FPs in the cellular environment can alter these properties[8,22-24]. As such, we next assessed the sensitivity of the acceptor mechanical switching signature to mechanical switching parameters. Increasing the acceptor force threshold $F^A_{1/2}$ increases the load magnitude and $E_{app}$ point at which $S_{app}$ begins to increase (Figure S5d). After the departure from $S_{app} = 0.5$, all ($E_{app}$, $S_{app}$)-curves exhibit monotonic decreases in $E_{app}$ and increases in $S_{app}$with increasing load magnitude. In comparison, increasing the acceptor kinetic rate constant $k^A_{MS,0}$ increases the slope of the ($E_{app}$, $S_{app}$) curve (Figure S5e). Again, all ($E_{app}$, $S_{app}$)-curves exhibit monotonic decreases in $E_{app}$ and increases in $S_{app}$with increasing load magnitude. Therefore, these data indicate that the ($E_{app}$, $S_{app}$)-curve has differential sensitivity to acceptor mechanical switching parameters, but that the overall up/left slanting ($E_{app}$, $S_{app}$) trend applies for all cases of acceptor mechanical switching in the loading regimes where it occurs.

Taken together, these data indicate that acceptor mechanical switching has a unique up/left slanting ($E_{app}$, $S_{app}$) signature that is robust to FP mechanical switching parameters and can be distinguished from cases of no acceptor mechanical switching as well as force-independent acceptor loss-of-function.

#### 3. Effect of Donor Mechanical Switching in Dynamic MTSs

To assess the ability to detect donor mechanical switching inside MTS, we compared readouts for an MTS with no donor mechanical switching or other loss-of-function (Figure S6c, brown line), an MTS with force-independent donor loss-of-function such as due to photobleaching (Figure S6c, grey line), and an MTS with donor mechanical switching (Figure S6c, black line; base parameters in Table S2). We looked at ($E_{app}$, $S_{app}$)-curves for load magnitude $F$ from 0 to 30 pN and a load duration $\tau$ of 1 second. In the absence of donor mechanical switching (Figure S6c, brown line), the ($E_{app}$, $S_{app}$)-curve starts at ($E_{app} = E_0(F = 0) = 0.286$ , $S_{app} = 0.5$), corresponding to an unloadable tension sensor module, and as the load magnitude increases $E_{app}$ decreases while $S_{app}$ remains constant. Thus, an MTS without donor mechanical switching exhibits a horizontal ($E_{app}$, $S_{app}$) signature at $S_{app} = 0.5$. In the case of force-independent donor loss-of-function (Figure S6c, grey line), the ($E_{app}$, $S_{app}$)-curve starts at the same $E_{app}$ but a lower $S_{app}$ than that of the unloaded tension sensor module. As load magnitude increases, $E_{app}$ decreases but $S_{app}$ does not change. Thus, an MTS with force-independent donor loss-of-function exhibits a horizontal ($E_{app}$, $S_{app}$) signature at $S_{app} < 0.5$. For the case of donor mechanical switching (Figure S6c, black line), the ($E_{app}$, $S_{app}$)-curve again starts at the point corresponding to the unloaded tension sensor module. While $E_{app}$ again decreases with load magnitude, $S_{app}$ decreases with load magnitude for loads near and above the FPs threshold force ($F \geq F^D_{1/2}$). Thus, donor mechanical switching exhibits a down/left slanting ($E_{app}$, $S_{app}$) signature. Furthermore, the response these data have to donor mechanical switching parameters resembles the response described for acceptor mechanical switching in the previous section. Briefly, donor force threshold ($F^D_{1/2}$) controls the $E_{app}$ value at which the ($E_{app}$, $S_{app}$)-curve diverges from $S_{app} = 0.5$ and the donor kinetic rate constant ($k^D_{MS,0}$) controls the slope of the ($E_{app}$, $S_{app}$)-curve, but in all cases the down/left slanting ($E_{app}$, $S_{app}$) signature for donor mechanical switching remains (Figure S6d-e).

Taken together, these data indicate that donor mechanical switching has a unique down/left ($E_{app}$, $S_{app}$) signature that is robust to FP mechanical switching parameters and can be distinguished from cases of no donor mechanical switching as well as force-independent donor loss-of-function.

#### 4. FP Mechanical Switching in MTSs is Sensitive to both Load Magnitude and Duration

The mechanical switching of single FPs in load-bearing proteins was sensitive to both load magnitude and duration (Figure S1), so we next assessed if FRET measurements of MTSs with acceptor or donor mechanical switching were sensitive to both loading parameters. To assess the effect of load duration at a given load magnitude, we varied the load duration $\tau$ over a range from 0.01 to 100 seconds at constant load magnitude $F$. For acceptor mechanical switching, the ($E_{app}$, $S_{app}$)-curve moves up/left with increasing load duration, tracking along the tension isocline (Figure S7a). For donor mechanical switching, the ($E_{app}$, $S_{app}$)-curve moves vertically down with increasing load duration, similarly tracking along the tension isocline (Figure S7b).

For comparison, we assessed the effect of load magnitude at a given load duration. To do so, we varied the load magnitude $F$ over a range from 0 to 30 pN at constant load duration $\tau$ (Figure S7c-d). For the case of acceptor mechanical switching, $S_{app}$ always increases and $E_{app}$ always decreases in response to increases in either load magnitude (Figure S7c) or duration (Figure S7a). Thus, both produce up/left

($E_{app}$, $S_{app}$)-curves, with slopes being steeper for load duration variation compared to load magnitude variation. For the case of donor mechanical switching, $S_{app}$ always decreases in response to increases in either load magnitude (Figure S7d) or duration (Figure S7b), and $E_{app}$ either decreases or remains constant, for changes in load magnitude or duration, respectively.

Taken together, this data suggests that the ($E_{app}$, $S_{app}$) signatures for FP mechanical switching in MTSs are sensitive to changes in both load magnitude and duration and that they respond to these loading parameters differently.

**5. Dominant mechanical switching in one FP is detectable in presence of weaker mechanical switching in the other FP**

We next assessed how the ability to detect mechanical switching in one FP is affected by the presence of weaker levels of mechanical switching in the other FP. To do so, we modeled one FP exhibiting mechanical switching with the base parameters (Table S2) and the other FP exhibiting lower levels of mechanical switching (lower rate constant $k_{MS,0}^{D}/k_{MS,0}^{A}$ or higher force threshold $F_{1/2}^{D}/F_{1/2}^{A}$). For acceptor mechanical switching in the presence of donor mechanical switching with lower rate constant ($k_{MS,0}^{D} < k_{MS,0}^{A}$) but equal force threshold ($F_{1/2}^{D} = F_{1/2}^{A}$), increasing amounts of donor mechanical switching (increasing $k_{MS,0}^{D}$) leads to a reduction in $S_{app}$ (Figure S8a, blue and orange lines) compared to case of acceptor mechanical switching only (Figure S8a, black line). The up/left ($E_{app}$, $S_{app}$)-curve is preserved but with a reduced slope. Further increases in donor mechanical switching up to the level of acceptor mechanical switching causes $S_{app}$ to return to 0.5 and the ($E_{app}$, $S_{app}$)-curve to have zero slope (Figure S8a, yellow line; data signatures of identical mechanical switching parameters for donor and accepter are discussed further in Section IV). Donor mechanical switching in the presence of acceptor mechanical switching with lower rate constant ($k_{MS,0}^{A} < k_{MS,0}^{D}$) but equal force threshold ($F_{1/2}^{A} = F_{1/2}^{D}$) displays a similar behavior but with effects in the opposite direction (Figure S8b). Therefore, in both cases, ($E_{app}$, $S_{app}$)-curves for dominant acceptor or donor mechanical switching in the presence of lower levels of mechanical switching in the other FP retain their up/left or down/left signatures, respectively.

We next investigated differences in the FP force threshold. For acceptor mechanical switching in the presence of donor mechanical switching with higher force threshold ($F_{1/2}^{D} > F_{1/2}^{A}$) but equal constant ($k_{MS,0}^{D} = k_{MS,0}^{A}$), effects on the ($E_{app}$, $S_{app}$)-curves are only seen at load magnitudes near and above $F_{1/2}^{D}$ (Figure S8c). Above these load magnitudes, $S_{app}$ reduces with increased load magnitude, approaching 0.5. A similar effect in the opposite direction is observed for the case of dominant donor mechanical switching (Figure S8d). Despite changes in the ($E_{app}$, $S_{app}$)-curve at higher loads, the deviations in $S_{app}$ from 0.5 as well as the shape of the ($E_{app}$, $S_{app}$)-curve at lower loads remained unchanged. As such, they remain reliable indicators of the FP that exhibits dominant mechanical switching.

Taken together, this data demonstrates that dominant acceptor or donor mechanical switching in the presence of lower levels of mechanical switching in the other FP are detectable from their ($E_{app}$, $S_{app}$) signatures.

### IV. Visualization of Experimental Data

To facilitate visualization of experimental data, we next conducted simulations that account for phenomena likely to be present in experimental data, including variability in protein loading dynamics and intrinsic noise due to inherent stochasticity of kinetic processes. Protein loading dynamics likely vary within and between cells. For instance, F-actin flow speeds and polymerization rates (two sources of protein loading in cells) vary spatially with distance from the cell edge[30]. Additionally, traction stresses and vinculin molecular tension vary between and within focal adhesions, and heterogeneity of traction stresses across cell populations has also been reported[13,24,31-33]. Together, this suggests subcellular heterogeneity in protein loading dynamics (i.e. differences in protein loading dynamics between populations of mechanical proteins in different cells and at different subcellular locations in the same cell). Therefore, we conducted simulations where the load magnitude and/or duration for each ensemble were drawn from distributions.

#### A. Model Formulation

##### 1. Stochastic Simulations

We simulated ensembles of MTSs consisting of $n_{sensors}$ sensors that are subject to dynamic loading and undergo FP mechanical switching according to the kinetic model for FP mechanical switching in MTSs described in Section III.A. All MTSs in a given ensemble exhibit the same FP mechanical switching parameters and are subject to the same dynamic loading conditions, parameterized by the load magnitude $F$ and a characteristic load duration $\tau$ ($\tau \equiv 1/k_{unbind}$). Stochastic simulations were performed using the Gillespie Algorithm[34], starting with all MTSs in the D1A1 state and running until the number of sensors in each of the 4 MTS states reached a steady state. Then, the state of each sensor was sampled, and this information was used to compute the experimentally observable readouts of three channel FRET as described in Section III.A.

##### 2. Simulations with Distributions in Load Magnitude and Duration Between Ensembles

For each parameter combination, we simulated $N_{sims} = 1000$ MTS ensembles each containing $n_{sensors} = 50$ total sensors. The apparent FRET Eff ($E_{app}$) and Stoichiometry ($S_{app}$) was computed for each simulation and displayed in ES histograms. As protein loading dynamics likely vary within and between cells[13,24,31-33], we conducted simulations where the load magnitude and/or duration for each ensemble were drawn from distributions. As little is known about the distribution of forces on proteins inside cells, and because we sought to interpret the data for both synthetic MTSs and MTSs for naturally occurring proteins, we chose simple generic distributions. Specifically, the $k$-th ensemble is assigned a load magnitude $F_k$ that is drawn from a uniform distribution on the interval 0 to 10 pN, i.e. $F_k \sim Uniform(0{,}10)$. Likewise, the characteristic load duration $\tau_k$ was drawn from a log-uniform distribution from $10^{-0.5}$ to $10^{0.5}$ s, i.e. $\log_{10}(\tau_k) \sim Uniform\big(\log_{10}(-0.5), \log_{10}(0.5)\big)$. As previously defined, the unloading rate constant was $k_{unbind,k} = 1/\tau_k$. Within each single ensemble, all MTSs are subjected to the same loading conditions.

#### B. Results

To facilitate data visualization, we investigated signatures of FP mechanical switching in MTSs in the presence of variability in the protein loading conditions as well as intrinsic noise due to inherent stochasticity of kinetic processes. We considered four scenarios: no FP mechanical switching, only acceptor mechanical switching (with base parameters in Table S2), only donor mechanical switching (with base parameters in Table S2), and both acceptor and donor mechanical switching at equal levels (with base parameters in Table S2). For each scenario, we investigated the ($E_{app}$, $S_{app}$)-distribution for variable load magnitude, variable load duration, and variable load magnitude and load duration. The histograms presented in this section match all trends described in Section III, but the histograms are expected to provide more realistic data signatures for comparison to experimental data as they account for phenomena likely present in experimental data, including variability in protein loading dynamics and intrinsic noise due to inherent stochasticity of kinetic processes.

We first considered variations in load magnitude only. Here, the ($E_{app}$, $S_{app}$)-distribution in the absence of FP mechanical switching is horizontal along $S_{app} = 0.5$ to the left of the unloaded $E_{app} = 0.286$ (Figure S9a). For acceptor only mechanical switching, the ($E_{app}$, $S_{app}$)-distribution has an up/left sloping signature (Figure S9d). For donor only mechanical switching, the ($E_{app}$, $S_{app}$)-distribution has a down/left sloping signature (Figure S9g). These results match the trends described in detail in Figures S5-S6 and Section III.B.2-3. For both acceptor and donor mechanical switching at equal levels, the ($E_{app}$, $S_{app}$)-distribution is centered horizontally along $S_{app} = 0.5$ with increasing spread in $S_{app}$ at lower $E_{app}$ (Figure S9j). This case of acceptor and donor mechanical switching at equal levels (Figure S9j) is clearly distinguishable from the case of no FP mechanical switching (Figure S9a) by the significantly increased spread in $S_{app}$ at lower $E_{app}$.

We next considered variations in load duration only. Here, the ($E_{app}$, $S_{app}$)-distribution in the absence of FP mechanical switching occupies a single point at $S_{app} = 0.5$ and the $E_{app} = E_0(F)$ (Figure S9b), and the cases of acceptor only (Figure S9e) or donor only (Figure S9h) mechanical switching have up/left or vertical down ($E_{app}$, $S_{app}$)-distributions, as described in detail in Figure S7 and Section III.B.4. For both acceptor and donor mechanical switching at equal levels, the ($E_{app}$, $S_{app}$)-distribution lies on and to the left of an ($E_{app}$, $S_{app}$) point with spreads in both $E_{app}$ and $S_{app}$ (Figure S9k).

We lastly considered variations in both load magnitude and load duration (Figure S9c,f,i,l). For all four cases of FP mechanical switching, the ($E_{app}$, $S_{app}$)-distributions closely resemble the corresponding distributions for variations in load magnitude only, except that there is increased spread in $S_{app}$ at lower $E_{app}$ for the three cases of FP mechanical switching (Figure S9f,i,l).

Taken together, these data provide unique data signatures for experimental data containing no FP mechanical switching, only acceptor mechanical switching, only donor mechanical switching, and both acceptor and donor mechanical switching at equal levels in the presence of cell-to-cell and/or subcellular heterogeneity in protein loading as well as intrinsic noise.

### V. Force-Dependent Bonds and Variable Loading Rates

Mechanical forces affect the lifetime of most intermolecular bonds[35]. Many of the mechanical proteins that transmit forces within the cytoskeleton or between the cytoskeleton and the extracellular environment exhibit force-sensitive bonds, and force-sensitive bond dynamics play an important role in mechanosensitive processes[2,5]. For example, the mechanical linker protein vinculin exhibits a force-activated bond with F-actin[15], and its turnover at focal adhesions is stabilized by molecular tension[1]. Furthermore, cells transmit forces to the ECM and sense mechanical properties of the ECM through molecular linkages at FAs[2,5,36]. These linkages couple the ECM to the actin cytoskeleton, transmitting forces from retrograde flowing actin[2,5,36]. Importantly, alterations in ECM stiffness are thought to be sensed through changes in the loading rate of molecular linkages in FAs[5]. Therefore, force-sensitive bond dynamics and loading rate are two important aspects of protein loading dynamics. In this section we extend our model of FP mechanical switching in MTSs to assess effects of force-sensitive bond dynamics and loading rate.

#### A. Model Formulation

#### 1. Force-Sensitive Unbinding Rate Constants

Three generic bond models are commonly used to represent the major responses of bonds to mechanical force: insensitivity, destabilization, and stabilization. An **ideal bond**, identical to $k_{unbind}$ in the preceding sections, represents a bond that is insensitive to load magnitude $F$ on the scale of those experienced by load-bearing proteins inside cells:

$$k_{unbind}(F) = k_{off,0} \tag{S20}$$

A **slip bond** represents a destabilizing bond whose lifetime decreases with load magnitude $F$ on the scale of those experienced by load-bearing proteins inside cells:

$$k_{unbind}(F) = k_{off,0}e^{F/F_b} \tag{S21}$$

where $k_{off,0}$ is the intrinsic unbinding rate constant and $F_b$ determines the sensitivity of the bond to force. Lastly, a two-pathway **catch-slip bond** is often used to represent a stabilizing bond whose lifetime initially increases with load magnitude $F$ before decreasing at higher load magnitudes. We define a rate constant that fits this form as follows:

$$k_{unbind}(F) = k_{off,0}\left(0.9e^{-2F/F_b} + 0.1e^{F/F_b}\right) \tag{S22}$$

where $k_{off,0}$ is the intrinsic unbinding rate constant and $F_b$ determines the sensitivity of the bond to force.

#### 2. Incorporation of Force-Sensitive Bonds into the Model of FP Mechanical Switching in MTSs

An additional model was implemented exactly as described in Section III but with replacing the unbinding rate constant for the MTS, $k_{unbind}$, with one of the three $k_{unbind}(F)$ functions (Equations S20-S22) in the steady state analysis (Equations S6-S9). $E_{app}$ and $S_{app}$ were computed as in Section III using Equations S18-S19.

#### 3. Loading Rate Control

These methods apply to the data in Section V.B.2 only. Instead of subjecting each MTS to the same load magnitude $F$ as before, each MTS was now subjected to the same loading rate $dF/dt$. All MTSs in an ensemble exhibit the same FP mechanical switching parameters and have the same unbinding rate constant $k_{unbind}(F)$. Stochastic simulations were conducted to analyze this model. Specifically, MTS ensembles were simulated according to the methods and parameters in Section IV, except for the following modifications. For each sensor $i$ in an ensemble, the force $F_i$ was updated at each timestep $dt$ using the equation $F_i = (dF/dt) \cdot dt$. Upon unbinding, which occurred with a rate $k_{unbind}(F_i)$, $F_i$ was reset to 0, corresponding to the unloaded state, followed by FP recovery (if applicable) and immediate re-binding and loading. In this scenario, each MTS inside a single ensemble can have a different $F_i$ value, and thus different $E_i$ value. As a result, $E_{app}$ and $S_{app}$ must be computed directly from the channel intensities using Equations S12-S17. Additionally, due to the enhanced complexity, no analytical expressions for average steady state species abundances or $E_{app}$ and $S_{app}$ were derived. Instead, ($E_{app}$, $S_{app}$)-curves were plotted using the averages from stochastic simulations. For each parameter combination, we simulated $N_{sims} = 100$ MTS ensembles each containing $n_{sensors} = 50$ total sensors.

#### B. Results

##### 1. Effect of Force-Sensitive Bond Dynamics on FP Mechanical Switching

To assess how FP mechanical switching in an MTS is affected by force-dependent bond dynamics, we investigated bonds whose durations are insensitive (ideal; Equation S20), destabilized (slip; Equation S21), or stabilized (catch-slip; Equation S22) by forces on the scale of those experienced by load-bearing proteins inside cells (Figure S10a-c). Acceptor or donor mechanical switching parameters were held constant at the base values from Table S2.

We first assessed effects associated with the functional form of the three bond models. To do so, we compared bonds with identical intrinsic rate constants ($k_{off,0} = 1$ sec) and with equal force-sensitivities for the slip and catch-slip bonds ($F_b = 5$ pN) (Figure S10d). The up/left slanting ($E_{app}$, $S_{app}$) signature of acceptor mechanical switching for the ideal bond has already been described (Section III). At low and intermediate loads, the slip and catch-slip bonds both match the up/left slanting ($E_{app}$, $S_{app}$) signature of the ideal bond (Figure S10e). Over this range of loads, slip bonding reduces the amount of FP mechanical switching by reducing the loading duration, while catch bonding increases the amount of FP mechanical switching by increasing the loading duration. At higher loads, both the slip and catch-slip bond exhibit decreased $S_{app}$ (i.e. a return toward 0.5) due to large reductions in bond duration at high forces, leading to rapid MTS unbinding and recovery of FP function. The sensitivity of FP mechanical switching to force-sensitive bonds is thus consistent with the previously determined importance of load duration (Figure S1 and Figure S7). For donor mechanical switching, a similar effect of slip and catch-slip bonding is observed, although with $S_{app}$ moving to lower values before increasing back toward 0.5 at higher loads (Figure S10f). Together, these results indicate that FP mechanical switching is sensitive to force-sensitive bond dynamics at load magnitudes near the characteristic force sensitivity of the bond, with slip bonding reducing FP mechanical switching and catch bonding enhancing FP mechanical switching.

To comprehensively assess effects of force-dependent bond dynamics on the ($E_{app}$, $S_{app}$) signatures, we next looked at slip (Figure S10g-i) and catch-slip (Figure S10j-l) bonds with a wide range of characteristic force sensitivities ($F_b$ from 1 to 25 pN). Slip and catch-slip bonds that rapidly destabilize at load magnitudes below the characteristic force threshold of the FP ($F_b \ll F_{1/2}^{A}$ or $F_b \ll F_{1/2}^{A}$) do not permit FP mechanical switching (e.g. orange line in Figure S10h-j,i-k). For slip and catch-slip bonds with higher $F_b$ values, FP mechanical switching is enhanced. At these higher $F_b$ values, slip and catch-slip bonds resemble the signatures of ideal bonds: up/left slanting ($E_{app}$, $S_{app}$) signatures for acceptor mechanical switching (Figure S10h,k) and down/left slanting ($E_{app}$, $S_{app}$) signatures for donor mechanical switching (Figure S10i,l), up to the load magnitude at which both slip and catch-slip bonds begin to rapidly destabilize causing decreases in $S_{app}$ toward 0.5 at the highest load magnitudes. We again see differences in the absolute position of the ($E_{app}$, $S_{app}$) data for ideal, slip, and catch-slip bonds at load magnitudes near $F_b$, confirming the response of FP mechanical switching to force-sensitive bond dynamics. Lasty, for sufficiently high values of $F_b$, slip and catch-slip bonds approach the behavior of an ideal bond, which is expected because as $F_b$ tends to infinity both bond models become ideal bonds.

Lastly, we assessed the effect of force-dependent bonds on ES-histograms of MTS populations subject to variable load magnitudes (analogous to Figure S9a,d,g,j), which represent effects of cell-to-cell and/or subcellular heterogeneity in protein loading as well as intrinsic noise (see Section IV). Specifically, we simulated populations of MTSs with ideal, slip, or catch-slip bonds subjected to a load magnitude ranging from 0 to 10 pN (uniformly distributed) with the cases of neither, acceptor only, donor only, or both FPs undergoing FP mechanical switching (Figure S11). Compared to the ideal bond (Figure S11d,g,j), the slip bond reduces the amount of FP mechanical switching for cases of acceptor only (Figure S11e), donor only (Figure S11h), and acceptor and donor (Figure S11k) mechanical switching, as indicated by $S_{app}$ values closer to 0.5 at intermediate and lower $E_{app}$ values. In contrast, the catch-slip bond increases the amount of FP mechanical switching at intermediate forces for cases of acceptor only (Figure S11f), donor only (Figure S11i), and acceptor and donor (Figure S11l) mechanical switching, as indicated by $S_{app}$ values farther away from 0.5 at intermediate $E_{app}$ values compared to ideal and slip bonds. As weaking of the catch-slip bond occurs above the maximum load magnitude (10 pN) simulated here, a restoration of $S_{app}$ values back toward 0.5 is not observed here but would exist for weaker catch-slip bonds and/or higher maximum forces.

Together, these analyses identify the effects of force-sensitive bonds on FP mechanical switching. Across low and intermediate loads, MTSs with slip or catch-slip bonding retain the general ($E_{app}$, $S_{app}$) signatures for acceptor (up/left) and donor (down/left) mechanical switching reported for ideal bonds (Sections III and IV). However, in this force regime, slip bonding reduces the amount of FP mechanical switching in comparison to ideal bonds by reducing the loading duration, while catch bonding increases the amount of FP mechanical switching by increasing the loading duration. Lastly, at high loads, both slip and catch-slip bonds lead to reductions in FP mechanical switching compared to ideal bonds due to large reductions in load duration. Overall, these results suggest that perturbations to force-sensitive bonding of an MTS can lead to drastic changes in the amount of FP mechanical switching within the MTS.

##### 2. Effect of Changes in Loading Rate on FP Mechanical Switching

To assess the effect of load rate, ensembles of MTS were simulated in which each MTS was loaded at a constant rate $dF/dt$ and was unloaded upon unbinding/rebinding (Figure S12a). As there are no direct measurements of protein loading rates inside cells, a range of loading rates (0.1 to 100 pN/s) was chosen to encompass recent estimates for the loading rate of integrin-based linkages (1-8 pN/s based on actin retrograde flow speeds and talin domain unfolding tensions)[37]. For these simulations, acceptor or donor mechanical switching parameters were held constant at the base values from Table S2. Unbinding rate constants for ideal, slip, or catch-slip bonds were used, all with identical intrinsic rate constants ($k_{off,0} = 1$ sec) and with equal force-sensitivities for the slip and catch-slip bonds ($F_b = 5$ pN) (Figure S12b). For acceptor mechanical switching in an MTS with an ideal bond, increasing the loading rate produces an up/left movement of the ($E_{app}$, $S_{app}$) data (simulated MTS populations for 4 different loading rates shown separately in Figure S12c, and averages are plotted together in Figure S12f). This indicates that acceptor mechanical switching increases with loading rate across all probed rates when MTSs have an ideal bond. This high degree of FP mechanical switching occurs for an ideal bond because higher load magnitudes are reached at higher loading rates when bond lifetime (and thus load duration) is independent of load rate. For the slip bond, increases in loading rate produce a flatter up/left movement of the ($E_{app}$, $S_{app}$) data across lower loading rates, i.e. $S_{app}$ remains closer to 0.5 as $E_{app}$ decreases, and $S_{app}$ returns to 0.5 at higher loading rates due to reduced load duration (Figure S12d,f). For the catch-slip bond, increases in loading rate produce a steeper up/left movement of the ($E_{app}$, $S_{app}$) data across low and intermediate loading rates (compared to both the ideal and slip bonds), with $S_{app}$ again moving back toward 0.5 at the highest loading rates due to reduced load duration (Figure S12e,f). These trends arise for force-sensitive bonds because the distributions of load magnitude and duration are now set by the competition between loading rate and unbinding rate. For the slip bond this results in the highest levels of FP mechanical switching at lower loading rates. For the catch-slip bond this results in the highest levels of FP mechanical switching at intermediate loading rates. For donor mechanical switching, increased loading rate has analogous effects for all three types of bonds (ideal, slip, or catch-slip), although with $S_{app}$ moving to lower values before increasing back toward 0.5 (Figure S12g-j).

Overall, consistent with the previously demonstrated sensitivity to variations in load magnitude and/or duration (Figure S7), this data indicates that FP mechanical switching in MTSs is sensitive to changes in loading rate, and that the sensitivity to loading rate depends on the force-sensitivity of the bond mediating loading. As ECM stiffness is one factor that controls the loading rate of proteins in FAs[5], this data also suggests the existence of regimes where FP mechanical switching in MTS is sensitive to substrate stiffness.

### VI. Assumptions and Limitations

Here we discuss model assumptions and limitations. First, we cover those of the kinetic model of FP mechanical switching in MTSs. Then, we cover those of sensitized emission-based FRET measurements.

##### 1. Kinetic Model of Protein Loading Dynamics and FP Mechanical Switching

As little is known about the time-dependent loading profiles of proteins inside cells, we considered as a starting point two simple loading profiles, constant load magnitude or constant loading rate. In the first case, MTSs are subjected to a constant load magnitude $F$ for a load duration $\tau$, driven by stochastic unbinding from a loading source ( $\tau \equiv 1/k_{unbind}$). This step function loading profile is analogous to the force-clamp loading profile commonly used in single molecule experiments in vitro[6]. In the second case, MTSs are subjected to linear ramp loading (at constant loading rate $dF/dt$), again with stochastic unbinding from the loading source. This is analogous to the force-ramp loading profile used in single molecule experiments in vitro[38]. The loading profiles of mechanical linkages in cells are likely more complex than the step or ramp profiles used here, possibly involving forces or extensions that vary nonlinearly with time and/or saturate. Although differences in these loading profiles will affect the extent of FP mechanical switching, the simplified models presented here are sufficient for assessing how protein loading dynamics affect FP mechanical switching and thereby provide insight into the response of FP mechanical switching in more complex loading conditions.

Furthermore, the model focuses on the competing timescales of FP mechanical switching and protein loading dynamics. As such, following the unloading of an MTS, we used the simplifying assumption that FPs that underwent mechanical switching are immediately restored to the functional state. We note that this is consistent with the rapid recovery of FPs in unloaded MTSs and/or the exchange of unloaded MTSs with a cytoplasmic pool of MTSs in the D1A1 state. In reality, a mechanical protein at a specific structure in the cell (e.g. a focal adhesion or actin fiber) engages/disengages from a loading source (e.g. actin), potentially undergoing multiple loading cycles before exchanging from the disengaged state with a pool of freely diffusing cytoplasmic proteins. In this context, the characteristic load duration ($\tau \equiv 1/k_{unbind}$) represents the duration over which the MTS continuously experiences loads and neither FP recovery nor MTS turnover occurs. For a protein that rapidly turns over following disengagement, like a rapidly diffusing actin binding protein, $\tau$ corresponds more closely to the turnover timescale of the protein. On the other hand, for a protein that undergoes many cycles of engagement/disengagement (loading/unloading), like a slowly diffusing trans-membrane protein, $\tau$ corresponds more closely to the disengagement timescale of the protein, which would be driven by the unbinding rate constant for the shortest-lived bond in the mechanical linkage (e.g. ECM:integrin:talin:vinculin:F-actin). Overall, the model is sufficient to assess the competing timescales of FP mechanical switching versus MTS loading dynamics. In the future, more complex models explicitly incorporating binding, unbinding, and turnover as separate processes could be used to explore the interactions between more processes, like MTS rebinding versus FP recovery or sensor turnover.

To investigate subcellular and cell-to-cell heterogeneity in protein loading[13,24,32,33], we implemented distributions in load magnitude and duration between MTS populations. As little is known about the distribution of forces on proteins inside cells, we chose simple generic distributions for these two parameters. Other load magnitude distributions, such as those with long tails at higher forces, will produce similar results for cases where the FP force thresholds ($F_{1/2}^{D}$ and $F_{1/2}^{A}$) are close to the mode of the distribution. Alternatively, if the FP force thresholds are considerably higher than the mode of the distribution, long tails at higher forces could result in rare FP mechanical switching events occurring exclusively at very high load magnitudes.

For the constant magnitude loading profile, we used the simplifying assumption that all MTSs in a single population (corresponding to a measured pixel) are each engaged and loaded to the same load magnitude and duration values. However, MTSs within single ensembles could experience variation in loading. For instance, mixed populations of engaged/loaded and disengaged/unloaded talin proteins have been reported[39]. In this case, each MTS will have a unique $E_0(F)$, meaning $E_{app}$ and $S_{app}$ must be computed directly from raw three channel FRET intensities (Equations 12-17), as was done for the constant loading rate simulations. An implication of sub-ensemble load variation is that MTSs under higher load magnitudes and longer load durations will preferentially undergo FP mechanical switching. This highlights that the presence of FP mechanical switching is likely to impact estimation of load magnitude from $E_{app}$ for all cases of acceptor and/or donor mechanical switching, i.e. not just for acceptor mechanical switching as would already be expected for a system with excess donors. However, we do not expect the trends for detecting acceptor and/or donor mechanical switching reported here to be affected by this.

Lastly, we assumed that the entire load applied across the MTS is felt by both FPs, consistent with both FPs being in series in the line of loading, and that mechanical switching of the two FPs are independent processes. This ignores possible changes to the applied force caused by deformations or unfolding of one FP, as well as potential state-dependent interactions between the FPs.

#### 2. Sensitized emission-based FRET measurements of MTS populations containing mechanically switched FPs

Next, we discuss assumptions and limitations related to measurements of FRET-based MTSs. First, to determine the signal contribution for each sensor state in three channel sensitized emission-based FRET measurements, we made assumptions about the effect of FP mechanical switching on the photophysical properties of FPs. To our knowledge, forced-induced changes in the excitation or emission wavelengths of FPs have not been described. Therefore, we assumed that donor FPs that have undergone mechanical switching cannot be excited by any excitation light in the optical system, and that acceptor FPs that have undergone mechanical switching cannot be excited by any excitation light in the optical system and also cannot accept energy from donor FPs. If this assumption is not met, a substantially more complex formalism is needed.

Furthermore, the FRET Efficiency-force relationship for sensors in the D1A1 state was modeled according to the original TSMod, mTFP1-$(GPGGA)_8$-mVenus, to facilitate comparisons to experimental data elsewhere in this paper using this module. However, the framework here can be applied immediately to TSMods based on other FPs and other genetically encoded FRET-based MTS using calibrated, unstructured tension sensing elements (e.g. repeats of GGSGGS[24]), whose FRET-force relationships are loading rate-independent and have no hysteresis[40]. Additionally, the framework should also be adaptable to threshold-like tension sensing elements that operate at or near equilibrium with calibrated FRET-force relationships that are loading rate-independent and have no hysteresis, such as those based on a single ultra-fast unfolding/refolding transition (e.g. HP35 and HP35st[41]). In contrast, the framework is not readily applicable for tension sensing elements exhibiting loading rate-dependence or hysteresis.

Lastly, there are limitations associated with very high levels of FP mechanical switching that may affect experimental data interpretation. First, if all sensors in some MTS ensembles (pixels) exist in the D0A0 state due to high levels of both acceptor and donor mechanical switching, some of the data will be undetectable and lead to an underestimation of FP mechanical switching in the system. Second, if there are especially low numbers of sensors in the D1A1 state, low signal-to-noise could cause uninterpretable $E_{app}$ values in real experiments. However, it should be noted that these very high levels of FP mechanical switching do not pose a limitation to the mathematical model itself. The concentration of sensors in the D1A1 state mathematically remained non-zero in the ODE models (Sections III and V), and the parameter values explored in the stochastic models resulted in non-zero numbers of sensors in the D1A1 state (Sections IV and V). If the number of sensors in the D1A1 state reaches zero, $E_{app}$ and $S_{app}$ can still be computed directly from raw three channel FRET intensities (Equations S12-S17), as the simplified expressions for $E_{app}$ and $S_{app}$ (Equations S18-S19) become invalid.

### VII. Conclusions

In this supplemental note we developed and applied mathematical models of fluorescent protein (FP) mechanical switching in load-bearing proteins and MTSs. To investigate the process of FP mechanical switching in the context of protein loading in cells, we first modeled the reversible mechanical switching of a single FP integrated into a load-bearing protein and assessed the extent of FP mechanical switching across load magnitudes and durations estimated for protein loading in cells (Section II). This model indicated that FP mechanical switching can occur at the load magnitudes and durations estimated for protein loading in cells for a range of FP parameters ($F_{1/2} \leq F$ and $1/k_{ms,0} \leq \tau$), and that the process exhibits sensitivity to both load magnitude and duration. Next, to assess how FP mechanical switching affects FRET-based MTSs, we extended the model of FP mechanical switching to an MTS containing a donor and an acceptor FP (Section III). Using three channel FRET measurements analogous to experimentally accessible readouts, we developed and applied a framework for detecting FP mechanical switching in MTSs. We found that acceptor mechanical switching has a unique up/left slanting ($E_{app}$, $S_{app}$) signature that is robust to FP mechanical switching parameters and can be distinguished from force-independent acceptor loss-of-function. Donor mechanical switching has a unique down/left ($E_{app}$, $S_{app}$) signature that is robust to FP mechanical switching parameters and can be distinguished from force-independent donor loss-of-function. Both acceptor and donor FP mechanical switching signatures remain detectable in the presence of lower levels of mechanical switching in the other FP. Also, the ($E_{app}$, $S_{app}$) signature for FP mechanical switching in MTSs are sensitive to changes in both load magnitude and duration, and they respond to these loading parameters differently. To facilitate the data visualization, we conducted simulations that account for phenomena expected in real experimental data, including subcellular and cell-to-cell variation in protein loading dynamics and intrinsic noise in FP mechanical switching (Section IV). These data provide unique data signatures for comparison to experimental data. Lastly, consistent with the effects of loading dynamics on FP mechanical switching, we also demonstrate that FP mechanical switching is sensitive to force-sensitive bond dynamics and changes in loading rate (Section V), both of which are thought to underly stiffness sensing by FAs[5]. Overall, our model of FP mechanical switching in MTSs establishes a framework for the detection of FP mechanical switching in MTSs. Using this framework, existing and new MTSs can be assessed for the existence of FP mechanical switching, which should become a new step in the development and interpretation of all MTSs. Furthermore, as FP mechanical switching in MTSs is sensitive to various aspects of protein loading dynamics, we expect that FP mechanical switching could be leveraged in the to probe force-sensitive protein function in the cellular context.

### Note S2: Steered Molecular Dynamics Simulations of FPs

#### I. Introduction

Our experimental findings suggest that mVenus has a lower mechanical stability than mTFP1. As an independent means of testing this, we performed steered molecular dynamics (SMD) on the two FPs.

#### II. Methods

Structures of mTFP1, mVenus, and αGFP were all obtained using AlphaFold[42]. SMD simulations were prepared on a laptop computer using the QwikMD plugin[43] of VMD 1.9.4a53[44]. Default "Easy Run" settings were used (T=27 °C, 0.15 mol/L salt concentration, and implicit solvent). The C-terminal residue was selected as the pulling residue, and the N-terminal residue was selected as the anchoring residue. The prepared simulation files were then transferred to the Duke Compute Cluster (DCC) at Duke University. Each simulation was then run on the DCC on 24 cores with 2GB of RAM per core. The completed simulation results were then transferred back to a laptop computer and re-loaded in VMD. Force-extension curves were calculated within the "Advanced Analysis" tab of the QwikMD plugin and subsequently analyzed using MATLAB 2020b. All simulation results were visually inspected to ensure that beta-strand pullout occurred within the simulation. The unfolding force for each simulation was characterized as the maximum force between the initial timepoint and the user-identified timepoint at which the first beta strand was pulled out of the beta barrel. Visual snapshots were prepared in VMD. The initial state (i) snapshot was selected as a very early timepoint, the Δα snapshot was chosen as a timepoint just before the maximum force values shown via red circles in Figures S15-S16, and the Δαβ snapshot was selected as a timepoint where the force was close to zero and the full removal of one β strand from the β-barrel could be clearly seen.

#### III. Results

We performed SMD on FPs using an implicit solvent at pulling speeds ranging from 0.3 to 10 nm/ns using the QwikMD plugin of the Visual Molecular Dynamics software[44]. Simulations started with a relaxed FP structure at zero-force, and then pulled the N- and C-termini of the FP apart until at least one beta strand was pulled out of the FP's beta barrel. Rupture force was defined as the maximum force experienced by the FP before full pullout of the first beta-strand.

To validate our approach, we first simulated mechanical unfolding of αGFP, which has previously been studied using SMD by Saeger et al.[45] as well as by experimental and computational means in other studies[6,8]. The αGFP simulations had an average maximum rupture force of 412±64 pN (Figure S15 and Movie S1), which resembles rupture forces obtained via SMD on similar timescales in prior work[45,46]. Note that it is typical for force estimates obtained from SMD to be higher by roughly an order of magnitude than experimental estimates[6,8] because the pulling speeds used in SMD are much higher (~nm/ns) than typical experimental pulling speeds (~nm/ms), and molecular unfolding is a stochastic, time-dependent process. Such high SMD pulling speeds are necessary because MD simulations performed with readily accessible computational resources can generally only access ns timescales.

We then simulated mechanical unfolding of mTFP1 and mVenus (Figure S16 and Movies S2-S3). Consistent with our experimental results, mTFP1 exhibited a significantly higher ($p$ = 0.047, paired T-test) maximum force (409±61 pN) than mVenus (340±39 pN) (Figure S17). Notably, despite large differences in primary sequence between the two FPs, the same structural transitions occurred, albeit at different average forces. In both cases, the FP's handle was unfurled (called the Δα state based on prior work[8]), followed by the removal of the FP's N-terminal beta-strand (called the Δαβ state based on prior work[8]).

### Note S3: Supporting Tables for Statistical Tests

**Table S5. P-value from Wilcoxon Rank Sum Test for ABDTS and ABDTL in Live Condition in Figure 2.**
Data were considered not normal, so the two samples were compared by the Wilcoxon Rank Sum Test.

| Level | - Level | p-Value |
|---|---|---|
| ABDTS_Live_None | ABDTL_Live_None | <.0001 |

**Table S6. P-value from Wilcoxon Rank Sum Test for ABDTS and ABDTL in Fixed Condition in Figure S14.**
Data were considered not normal, so the two samples were compared by the Wilcoxon Rank Sum Test.

| Level | - Level | p-Value |
|---|---|---|
| ABDTS_Fix_None | ABDTL_Fix_None | <.0001 |

**Table S7. P-values from Steel-Dwass Test for ABDTS and ABDTL Data in Figure S18.**
Data were considered not normal and therefore were compared with a Kruskal-Wallis test (non-parametric one-way ANOVA on ranks). The Kruskal-Wallis test was significant (p<.0001), so Steel-Dwass non-parametric multiple comparisons test was conducted.

| Level | - Level | p-Value |
|---|---|---|
| ABDTS_Fix_DMSO | ABDTL_Fix_DMSO | <.0001 |
| ABDTS_Fix_DMSO | ABDTL_Fix_LatA | <.0001 |
| ABDTS_Fix_LatA | ABDTL_Fix_DMSO | <.0001 |
| ABDTL_Fix_LatA | ABDTL_Fix_DMSO | <.0001 |
| ABDTS_Fix_LatA | ABDTL_Fix_LatA | 0.0546 |
| ABDTS_Fix_LatA | ABDTS_Fix_DMSO | <.0001 |

**Table S8. P-values from Steel-Dwass Test for VinTS Variants on Glass in Figure 3.**
Data were considered not approximately normal and therefore were compared with a Kruskal-Wallis test (non-parametric one-way ANOVA on ranks). The Kruskal-Wallis test was significant (p<.0001), so Steel-Dwass non-parametric multiple comparisons test was conducted.

| Level | - Level | p-Value |
|---|---|---|
| VinTS E1015A_Glass | VinTS_Glass | 0.8863 |
| VinTS I997A_Glass | VinTS E1015A E1021A_Glass | 0.9578 |
| VinTS E1021A_Glass | VinTS_Glass | 0.0732 |
| VinTS I997A_Glass | VinTS E1021A_Glass | 0.0067 |
| VinTS I997A_Glass | VinTS_Glass | <.0001 |
| VinTS E1021A_Glass | VinTS E1015A_Glass | 0.0003 |
| VinTS E1015A E1021A_Glass | VinTS E1021A_Glass | <.0001 |
| VinTS I997A_Glass | VinTS E1015A_Glass | <.0001 |
| VinTS E1015A E1021A_Glass | VinTS_Glass | <.0001 |
| VinTS E1015A E1021A_Glass | VinTS E1015A_Glass | <.0001 |

**Table S9. P-value from Wilcoxon Rank Sum Test for VinTS Variants on PA Gels in Figure S20.**
Data were considered not normal, so the two samples were compared by the Wilcoxon Rank Sum Test.

| Level | - Level | p-Value |
|---|---|---|
| VinTS E1015A E1021A_PAGel | VinTS_PAGel | 0.3266 |

### Supplemental References


1 Rothenberg, K. E., Scott, D. W., Christoforou, N. & Hoffman, B. D. Vinculin Force-Sensitive Dynamics at Focal Adhesions Enable Effective Directed Cell Migration. *Biophys J* **114**, 1680-1694, doi:10.1016/j.bpj.2018.02.019 (2018).

2 Hoffman, B. D., Grashoff, C. & Schwartz, M. A. Dynamic molecular processes mediate cellular mechanotransduction. *Nature* **475**, 316-323, doi:10.1038/nature10316 (2011).

3 Roca-Cusachs, P., Iskratsch, T. & Sheetz, M. P. Finding the weakest link: exploring integrin-mediated mechanical molecular pathways. *J Cell Sci* **125**, 3025-3038, doi:10.1242/jcs.095794 (2012).

4 Hoffman, B. D. & Yap, A. S. Towards a Dynamic Understanding of Cadherin-Based Mechanobiology. *Trends Cell Biol* **25**, 803-814, doi:10.1016/j.tcb.2015.09.008 (2015).

5 Elosegui-Artola, A., Trepat, X. & Roca-Cusachs, P. Control of Mechanotransduction by Molecular Clutch Dynamics. *Trends Cell Biol* **28**, 356-367, doi:10.1016/j.tcb.2018.01.008 (2018).

6 Ganim, Z. & Rief, M. Mechanically switching single-molecule fluorescence of GFP by unfolding and refolding. *Proc Natl Acad Sci U S A* **114**, 11052-11056, doi:10.1073/pnas.1704937114 (2017).

7 Wolfenson, H., Bershadsky, A., Henis, Y. I. & Geiger, B. Actomyosin-generated tension controls the molecular kinetics of focal adhesions. *J Cell Sci* **124**, 1425-1432, doi:10.1242/jcs.077388 (2011).

8 Dietz, H. & Rief, M. Exploring the energy landscape of GFP by single-molecule mechanical experiments. *Proc Natl Acad Sci U S A* **101**, 16192-16197, doi:10.1073/pnas.0404549101 (2004).

9 Bustamante, C., Chemla, Y. R., Forde, N. R. & Izhaky, D. Mechanical processes in biochemistry. *Annu Rev Biochem* **73**, 705-748, doi:10.1146/annurev.biochem.72.121801.161542 (2004).

10 Finer, J. T., Simmons, R. M. & Spudich, J. A. Single myosin molecule mechanics: piconewton forces and nanometre steps. *Nature* **368**, 113-119, doi:10.1038/368113a0 (1994).

11 Molloy, J. E., Burns, J. E., Kendrick-Jones, J., Tregear, R. T. & White, D. C. Movement and force produced by a single myosin head. *Nature* **378**, 209-212, doi:10.1038/378209a0 (1995).

12 Footer, M. J., Kerssemakers, J. W., Theriot, J. A. & Dogterom, M. Direct measurement of force generation by actin filament polymerization using an optical trap. *Proc Natl Acad Sci U S A* **104**, 2181-2186, doi:10.1073/pnas.0607052104 (2007).

13 Gardel, M. L. *et al.* Traction stress in focal adhesions correlates biphasically with actin retrograde flow speed. *J Cell Biol* **183**, 999-1005, doi:10.1083/jcb.200810060 (2008).

14 Buckley, C. D. *et al.* Cell adhesion. The minimal cadherin-catenin complex binds to actin filaments under force. *Science* **346**, 1254211, doi:10.1126/science.1254211 (2014).

15 Huang, D. L., Bax, N. A., Buckley, C. D., Weis, W. I. & Dunn, A. R. Vinculin forms a directionally asymmetric catch bond with F-actin. *Science* **357**, 703-706, doi:10.1126/science.aan2556 (2017).

16 Owen, L. M., Bax, N. A., Weis, W. I. & Dunn, A. R. The C-terminal actin-binding domain of talin forms an asymmetric catch bond with F-actin. *Proc Natl Acad Sci U S A* **119**, e2109329119, doi:10.1073/pnas.2109329119 (2022).

17 Kong, F., Garcia, A. J., Mould, A. P., Humphries, M. J. & Zhu, C. Demonstration of catch bonds between an integrin and its ligand. *J Cell Biol* **185**, 1275-1284, doi:10.1083/jcb.200810002 (2009).

18 Rakshit, S., Zhang, Y., Manibog, K., Shafraz, O. & Sivasankar, S. Ideal, catch, and slip bonds in cadherin adhesion. *Proc Natl Acad Sci U S A* **109**, 18815-18820, doi:10.1073/pnas.1208349109 (2012).

19 Ferrer, J. M. *et al.* Measuring molecular rupture forces between single actin filaments and actin-binding proteins. *Proc Natl Acad Sci U S A* **105**, 9221-9226, doi:10.1073/pnas.0706124105 (2008).
20 Yao, M. *et al.* Mechanical activation of vinculin binding to talin locks talin in an unfolded conformation. *Sci Rep* **4**, 4610, doi:10.1038/srep04610 (2014).
21 Yao, M. *et al.* Force-dependent conformational switch of alpha-catenin controls vinculin binding. *Nat Commun* **5**, 4525, doi:10.1038/ncomms5525 (2014).
22 Perez-Jimenez, R., Garcia-Manyes, S., Ainavarapu, S. R. & Fernandez, J. M. Mechanical unfolding pathways of the enhanced yellow fluorescent protein revealed by single molecule force spectroscopy. *J Biol Chem* **281**, 40010-40014, doi:10.1074/jbc.M609890200 (2006).
23 Snapp, E. L. Fluorescent proteins: a cell biologist's user guide. *Trends Cell Biol* **19**, 649-655, doi:10.1016/j.tcb.2009.08.002 (2009).
24 LaCroix, A. S., Lynch, A. D., Berginski, M. E. & Hoffman, B. D. Tunable molecular tension sensors reveal extension-based control of vinculin loading. *Elife* **7**, doi:10.7554/eLife.33927 (2018).
25 Chen, H., Puhl, H. L., Koushik, S. V., Vogel, S. S. & Ikeda, S. R. Measurement of FRET efficiency and ratio of donor to acceptor concentration in living cells. *Biophys J* **91**, L39-L41, doi:10.1529/biophysj.106.088773 (2006).
26 Gates, E. M., LaCroix, A. S., Rothenberg, K. E. & Hoffman, B. D. Improving Quality, Reproducibility, and Usability of FRET-Based Tension Sensors. *Cytometry A* **95**, 201-213, doi:10.1002/cyto.a.23688 (2019).
27 Coullomb, A. *et al.* QuanTI-FRET: a framework for quantitative FRET measurements in living cells. *Sci Rep* **10**, 6504, doi:10.1038/s41598-020-62924-w (2020).
28 LaCroix, A. S., Rothenberg, K. E., Berginski, M. E., Urs, A. N. & Hoffman, B. D. Construction, imaging, and analysis of FRET-based tension sensors in living cells. *Methods Cell Biol* **125**, 161-186, doi:10.1016/bs.mcb.2014.10.033 (2015).
29 Grashoff, C. *et al.* Measuring mechanical tension across vinculin reveals regulation of focal adhesion dynamics. *Nature* **466**, 263-266, doi:10.1038/nature09198 (2010).
30 Ponti, A., Machacek, M., Gupton, S. L., Waterman-Storer, C. M. & Danuser, G. Two distinct actin networks drive the protrusion of migrating cells. *Science* **305**, 1782-1786, doi:10.1126/science.1100533 (2004).
31 Rajagopalan, P., Marganski, W. A., Brown, X. Q. & Wong, J. Y. Direct comparison of the spread area, contractility, and migration of balb/c 3T3 fibroblasts adhered to fibronectin- and RGD-modified substrata. *Biophys J* **87**, 2818-2827, doi:10.1529/biophysj.103.037218 (2004).
32 Trepat, X. *et al.* Physical forces during collective cell migration. *Nat Phys* **5**, 426-430, doi:10.1038/Nphys1269 (2009).
33 De La Pena, A., Mukhtar, M., Yokosawa, R., Carrasquilla, S. & Simmons, C. S. Quantifying cellular forces: Practical considerations of traction force microscopy for dermal fibroblasts. *Exp Dermatol* **30**, 74-83, doi:10.1111/exd.14166 (2021).
34 Gillespie, D. T. Exact stochastic simulation of coupled chemical reactions. *The Journal of Physical Chemistry* **81**, 2340-2361, doi:10.1021/j100540a008 (1977).
35 Evans, E. Probing the relation between force--lifetime--and chemistry in single molecular bonds. *Annu Rev Biophys Biomol Struct* **30**, 105-128, doi:10.1146/annurev.biophys.30.1.105 (2001).
36 Wolfenson, H., Yang, B. & Sheetz, M. P. Steps in Mechanotransduction Pathways that Control Cell Morphology. *Annu Rev Physiol* **81**, 585-605, doi:10.1146/annurev-physiol-021317-121245 (2019).
37 Liu, J. *et al.* Tension Gauge Tethers as Tension Threshold and Duration Sensors. *ACS Sens* **8**, 704-711, doi:10.1021/acssensors.2c02218 (2023).
38 Staple, D. B., Hanke, F. & Kreuzer, H. J. Dynamics of single-molecule force-ramp experiments: The role of fluctuations. *Physical Review E* **77**, 021801, doi:10.1103/PhysRevE.77.021801 (2008).

39 Ringer, P. *et al.* Multiplexing molecular tension sensors reveals piconewton force gradient across talin-1. *Nat Methods* **14**, 1090-1096, doi:10.1038/nmeth.4431 (2017).
40 Ham, T. R., Collins, K. L. & Hoffman, B. D. Molecular Tension Sensors: Moving Beyond Force. *Curr Opin Biomed Eng* **12**, 83-94, doi:10.1016/j.cobme.2019.10.003 (2019).
41 Fischer, L. S., Rangarajan, S., Sadhanasatish, T. & Grashoff, C. Molecular Force Measurement with Tension Sensors. *Annu Rev Biophys* **50**, 595-616, doi:10.1146/annurev-biophys-101920-064756 (2021).
42 Jumper, J. *et al.* Highly accurate protein structure prediction with AlphaFold. *Nature* **596**, 583-589, doi:10.1038/s41586-021-03819-2 (2021).
43 Ribeiro, J. V. *et al.* QwikMD - Integrative Molecular Dynamics Toolkit for Novices and Experts. *Sci Rep* **6**, 26536, doi:10.1038/srep26536 (2016).
44 Humphrey, W., Dalke, A. & Schulten, K. VMD: visual molecular dynamics. *J Mol Graph* **14**, 33-38, 27-38, doi:10.1016/0263-7855(96)00018-5 (1996).
45 Saeger, J., Hytonen, V. P., Klotzsch, E. & Vogel, V. GFP's mechanical intermediate states. *PLoS One* **7**, e46962, doi:10.1371/journal.pone.0046962 (2012).
46 Cheng, C.-L., Zhang, M.-Z. & Zhao, G.-J. Mechanical stability and thermal conductivity of β-barrel in green fluorescent protein by steered molecular dynamics. *RSC Advances* **4**, 6513-6516, doi:10.1039/C3RA42679C (2014).
